# Generalization in Sensorimotor Networks Configured with Natural Language Instructions

**DOI:** 10.1101/2022.02.22.481293

**Authors:** Reidar Riveland, Alexandre Pouget

## Abstract

One of humans’ most fundamental cognitive feats is the ability to interpret linguistic instructions in order to perform novel tasks without any explicit experience with the task. Yet, the computations that the brain might use to accomplish such a feat remains poorly understood. Here we use the latest advances in Natural Language Processing to create a neural model of generalization based on linguistic instructions. Models are trained on a set of commonly studied psychophysical tasks, and receive instructions embedded by a pre-trained language model. Our best models can perform a previously unseen task with a performance of 83% correct on average based solely on linguistic instructions (i.e. 0-shot learning). We found that language scaffolds sensorimotor representations such that activity for interrelated tasks share a common geometry with the semantic representations of instructions, allowing language to cue the proper composition of practiced skills in unseen settings. Finally, we show how this model can generate a linguistic description of a novel task it has identified using only motor feedback, which can subsequently guide a partner model to perform the task. Our models offer several experimentally testable predictions outlining how linguistic information must be represented in order to facilitate flexible and general cognition in the human brain.

## Introduction

In a laboratory setting, animals require numerous trials carefully presented over a period of weeks in order to acquire a new behavioral task. This is in part because the only means of communication with non-linguistic animals is simple positive and negative reinforcement signals. By contrast, it is common to give written or verbal instructions to human subjects which allows them to perform new tasks relatively quickly. For instance, a subject told to ‘respond in the direction of the stimulus presented with highest intensity’, would know what to do right away. Indeed, for simple tasks like this one, human subjects would perform nearly perfectly on the very first trial (i.e. 0-shot learning), even if they had no previous experience with the particular task. Alternatively, human subjects can also learn to correctly perform a task in the absence of linguistic instructions, e.g. through positive and negative reinforcement, and, once training is completed, subjects can typically explain how they solved the task with natural language.

The dual ability to use the information in an instruction to immediately perform a novel task and, conversely, produce a linguistic description of the demands of a task once it has been learned are two unique cornerstones of human communication. Yet, the computational principles and representational structures that underlie these abilities remain poorly understood.

One influential systems-level explanation of our ability to generalize performance posits that flexible inter-region connectivity in the prefrontal cortex (PFC) allows for the reuse of practiced sensorimotor representations in novel settings [1, 2, 3, 4, 5]. By recombining practiced stimulus-response patterns according to the ostensible demands of a previously unseen task, we can leverage well-established abilities and perform on new tasks in very few practice trials. More recently, multiple studies have observed that when subjects are required to flexibly recruit different stimulus-response patterns in order to switch between tasks, neural representations are organized according to the abstract structure of the task set [6, 7, 8, 9]. This disentangled geometry may allow for the easy reuse of task sub-components. Lastly, recent modeling work has begun to explore the neural mechanisms of compositional computation in a non-linguistic setting. For example, a recurrent neural network (RNN) trained on a set of psychophysical tasks will share dynamical motifs across tasks with similar demands (e.g. the same ring attractor to store a circular variable across tasks that require memory) [10]. By reusing underlying computational dynamics networks exhibit faster learning in novel settings. This work forms a strong basis for explanations of flexible cognition in humans but leaves open the essential question of how linguistic information can immediately reconfigure a sensorimotor network so that it performs a novel task well on the very first attempt. From the opposite perspective, few studies have investigated the linguistic and semantic processing hierarchy that directly informs sensorimotor control (although see [11], [12], [13], [14]). Overall it remains unclear what representational structure we should expect from brain areas that are responsible for integrating linguistic information in order to reorganize sensorimotor mappings on the fly.

These questions become all the more pressing given that recent advances in machine learning have led to artificial intelligent systems that exhibit human-like language skills, both within the language domain itself and in multi-modal settings [15, 16, 17]. Recent works have matched neural data recorded during passive listening and reading tasks to activations in autoregressive language models (e.g. GPT [18]), arguing that there is a fundamentally predictive component to language comprehension [19, 20]. However, comprehension during passive listening is only one of many interrelated notions of linguistic knowledge. Indeed, part of the essential utility of understanding language is not simply that we can exchange information between individuals, but further that each individual can correctly interpret and then execute the actions that this linguistic information suggests we take. Some high-profile machine learning models do show the ability to use natural language as a prompt to perform a linguistic task or render an image, but these models are designed with a specific engineering goal in mind and, especially in text-to-image models, this makes their outputs difficult to interpret in terms of a sensorimotor mapping that we might expect to occur in a biological system [21, 22, 15, 16, 17, 23]. Alternatively, recent work on multi-modal interactive agents may be far more interpretable in terms of the actions they take, but utilize a perceptual hierarchy that fuses vision and language at very early stages of processing, making them difficult to map onto functionally and anatomically distinct language and vision areas in human brains [24], [25], [26].

We therefore seek a setting where we can leverage the power of language models in a way that results in testable neural predictions detailing how the human brain processes natural language in order to generalize across sensorimotor tasks.

To that end, we train a recurrent neural network (Sensorimotor-RNN) model on a set of simple psychophysical tasks. On each trial, the model receives natural language instructions for the present task embedded through a transformer architecture pre-trained on one of several natural language processing objectives. Both the type of tasks we use and the neural network modeling of such tasks have a rich history in the experimental and computational neuroscience literature respectively [27, 28, 29, 30, 31, 10, 32].

We find that instruction embeddings produced by models trained explicitly to produce high quality embeddings of sentence level semantics allow Sensorimotor-RNNs to perform an entirely novel task at 83 correct on average. By contrast, models trained solely on next-word prediction performed relatively poorly despite massive advantages in model size and pre-training scope. Generalization in our models is supported by a representational geometry which captures task subcomponents and is shared between instruction embeddings and sensorimotor activity. In our best performing models, this structure emerges as activity ascends the language processing hierarchy. We also find that individual neurons modulate their tuning based on the semantics of instructions. We demonstrate how a network trained to interpret linguistic instructions can invert this understanding and produce a linguistic description of a previously unseen task based on the information in motor feedback signals. Communicating these instructions to a partner model results in a performance of 82% across tasks compared to 30% performance when directly sharing latent task representations, demonstrating the importance of natural language as an information encoding scheme that is shared between otherwise independent sets of neurons. We end by discussing how these results can guide research on the neural basis of language-based generalization in the human brain.

## Results

### Instructed models and task set

To examine how linguistic information leads to generalization across sensorimotor tasks, we train Sensorimotor-RNNs on a set of 50 interrelated psychophysical tasks that require various cognitive capacities which are well-studied in the literature [31]. These include perceptual decision-making [27, 30], inhibitory control [33], matching tasks [34, 35], confidence judgements [36], context-dependent tasks [28], time estimation, [29] and combinations of these and other skills. Two example tasks are presented in Fig. 1a, b as they might appear in a laboratory setting. For all tasks, models receive a sensory input and task-identifying information and must output motor response activity appropriate for the task (Fig. 1c). Input stimuli are encoded by two 1-D maps of neurons, each representing a different input modality, with periodic Gaussian tuning curves to angles (over [0, 2*π*]). As a result, a specific input value, i.e., a specific angle, is encoded by a hill of activity across the 1D maps. Output responses are encoded in the same way, via hills of activity, encoding specific output angles. Inputs also include a single fixation unit which tells the model when to maintain fixation during the trial, which is simulated by high activity in the fixation output of the model. After the input fixation is off the model can break its own fixation and respond to the input stimuli. Our 50 tasks are roughly divided into 5 groups, ‘Go’, ‘Decision Making’, ‘Comparison’, ‘Duration’ and ‘Matching’, where within group tasks share similar sensory input structures but may require divergent responses. For instance, in the decision making task (DM), the network must respond in the direction of the stimulus with the highest contrast immediately after the extinction of the fixation point whereas in the anti-decision making task (AntiDM), the network must instead respond in the direction of the stimulus with the weakest contrast (Fig. 1a). Given the possibility of such divergent responses, networks must properly infer the task demands for a given trial from task-identifying information in order to perform all tasks simultaneously (see Methods for task details, see Supplementary Info. 16 for example trials of all tasks).

**Figure 1.**
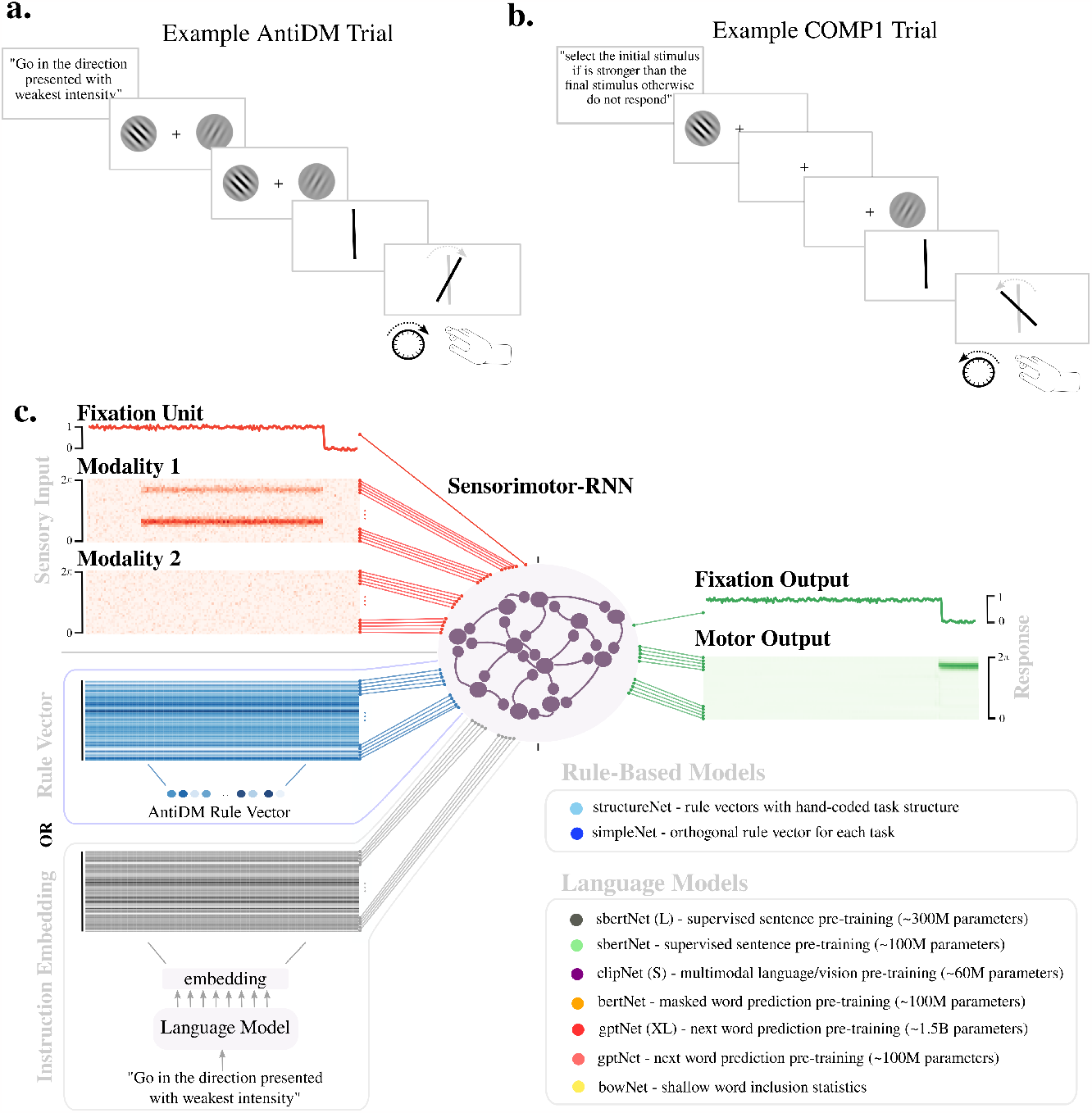
Tasks and Models. **a, b**. Illustration of example trials as they might appear in a laboratory setting. The trial is instructed, then stimuli are presented with different angles and strengths of contrast. The agent must then respond with the proper angle during the response period. **a**. An example ‘AntiDM’ trial where the agent must respond to the angle presented with the least intensity. **b**. An example ‘COMP1’ trial where the agent must respond to the first angle if it is presented with higher intensity than the second angle other repress response. **c**. Diagram of model inputs and outputs. Sensory inputs (Fixation Unit, Modality 1, Modality 2) are shown in red and model outputs (Fixation Output, Motor Output) are shown in green. Models also receive a rule vector (blue) or the embedding that results from passing task instructions through a pre-trained language model (grey). A list of models tested is provided in the inset.

In our models, task-identifying input is either non-linguistic or linguistic. We use two non-linguistic control models. First, in simpleNet, the identity of a task is represented by one of 50 orthogonal rule vectors which index each task. Second, structureNet uses a set of 10 orthogonal structure vectors which each represent a dimension of the task set (e.g. respond weakest vs. strongest direction, respond in the same vs. opposite direction, pay attention only to stimuli in the first modality etc.), and tasks are encoded using combinations of these vectors (see Supplementary Info. 14 for full set of structure combinations). These controls represent two extremes in terms of knowledge about the task set, where the task-identifying input to structureNet fully captures all of the relevant relationships amongst tasks, and, conversely, task-identifying input to simpleNet encodes none of this structure.

Instructed Models use a pre-trained transformer architecture [37] to embed natural language instructions for the tasks at hand. These embeddings are then passed to the Sensorimotor-RNN as task-identifying information (Fig. 1a, Instructed Models). For each task there is a corresponding set of 20 unique instructions (15 training, 5 validation, see Supplementary Info. 13 for full instruction set).

We test various types of language models that share the same basic architecture but differ in their size and also their pre-training objective, which endows them with different kinds of linguistic knowledge. First, we tested two autoregressive models, a standard and a large version of GPT2 which we call gpt and gpt (XL) respectively. These models are trained to predict the next word in a text given all the preceding context [18]. This allows the model to complete texts from prompts with striking fluency [38]. Previous work has demonstrated that gpt can account for continuous next-word prediction and degree of surprise for unexpected words in ECoG data from participants listening to narrative stories [20], whereas activations from the hidden layers of gpt (XL) are predictive of fMRI and EEG data taken from language areas while humans read sentences [19]. This validation against neural data makes it an important benchmark for computational models of language processing in humans. The bert model receives bi-directional inputs and is trained to identify masked words within a piece of text [39]. bert also uses a simple unsupervised sentence level objective, in which the network is given two sentences and must determine whether they follow each other in the original text. s-bert [40] is trained like bert but receives additional tuning on the Stanford Natural Language Inference task [41], a hand-labeled dataset detailing the logical relationship between two candidate sentences (see Methods). Lastly, we also use the language embedder from CLIP, a multi-modal model which learns a joint embedding space of images and their text captions [42].

gpt, bert, and s-bert all use a baseline size of ∼ 100M parameters. gpt (XL) is an order of magnitude larger with ∼ 1.5 billion parameters while CLIP uses ∼ 60M parameters. We also use a slight larger version of s-bert with ∼ 300M parameters. We call a Sensorimotor-RNN using a given language model to embed instructions languageModelNet and append a letter indicate its size relative to the baseline model size of ∼ 100M parameters where appropriate (e.g. clipNet (S) and sbertNet (L), see Fig. 1c. inset). For each language model we apply a pooling method to the last hidden state of the transformer and pass this fixed length representation through a set of linear weights that results in a 64-dimensional instruction embedding across all models (see Methods). This last set of linear weights trains with the Sensorimotor-RNN on our task set whereas transformer weights are frozen, unless specified otherwise. Finally, as a control we also test a Bag of Words (bow) embedding scheme that only uses word count statistics to embed each instruction.

First, we verify our models can perform all tasks simultaneously. For non-linguistic models, high performance across all tasks is simply a question of the representational power of the Sensorimotor-RNN, as the contextual information about each task is already perfectly separated. The problem is more complicated with linguistic inputs because models must first infer the common semantic content between 15 distinct instruction formulations for each task before it can properly select the relevant mapping in the Sensorimotor-RNN. We find that all of our instructed models can process linguistic instructions to perform the entire task set simultaneously except for gptNet, in which case performance asymptotes below the 95% threshold for some tasks. Hence, we relax the performance threshold to 85% for models which use gpt to embed instructions (Supplementary Info. Fig. 1, see Methods for training details).

To test whether our model can generalize performance to new linguistic instructions for the learned tasks, we tested all architectures on a set of validation instructions, with no further training (Supplementary Info. Fig. 2). In general, sbertNet (L) and sbertNet are our best performing models, achieving an average performance of 97% and 94% respectively on validation instructions, demonstrating that these networks infer the proper semantic content even for entirely novel instructions.

### Generalization to novel tasks

We next examine how training networks to perform tasks with natural language instructions can help those networks generalize performance to novel tasks. To this end, we trained individual networks on 45 tasks and then tested network performance when exposed to each of the 5 held out tasks. Results are shown in Fig. 2. We use unequal variances t-tests to make comparisons amongst the generalization performance of different models. Fig. 2. shows p-values for the most relevant comparisons and a full matrix of comparisons across all models can be found in Supplementary Info. 3, 4.

**Figure 2.**
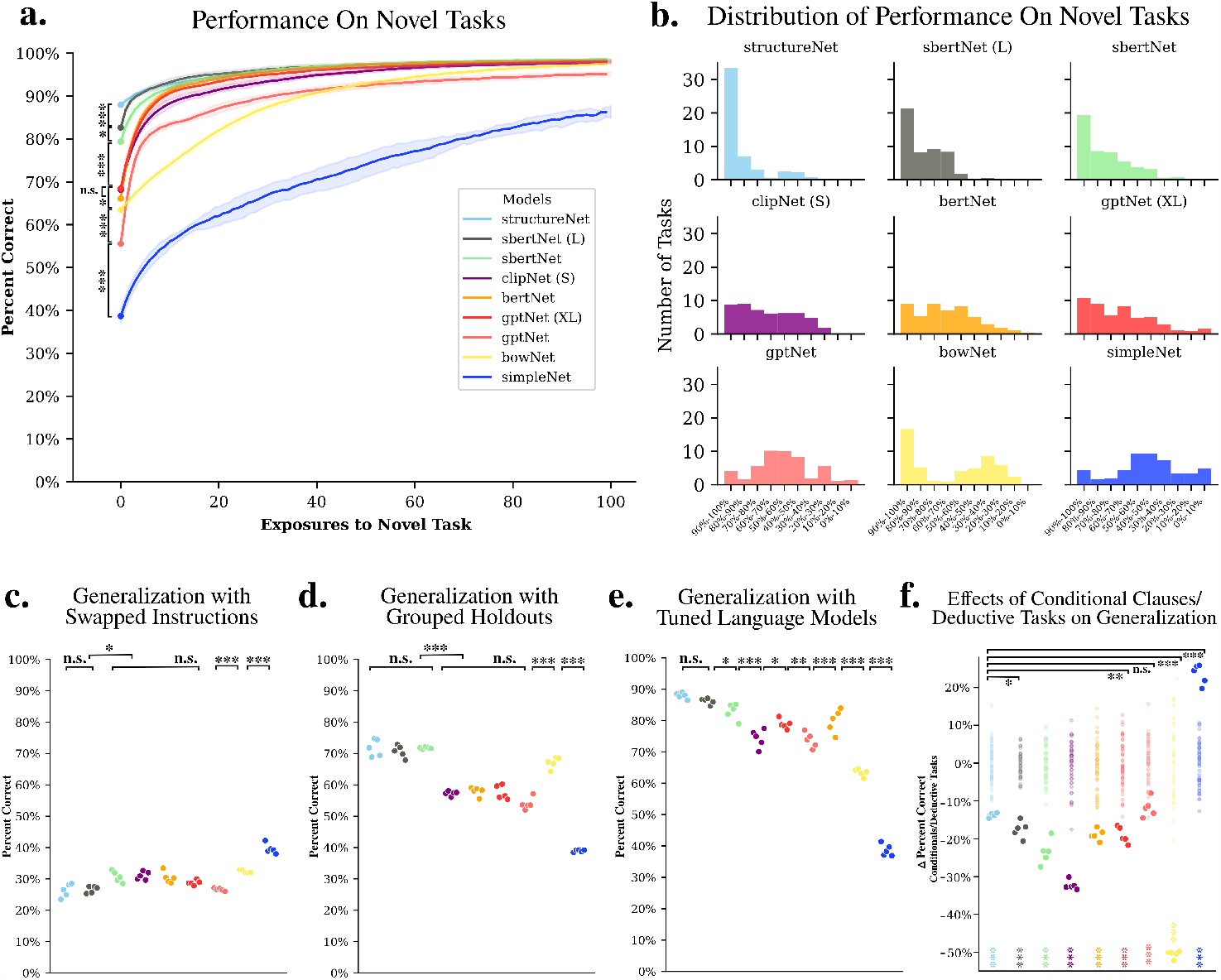
Model Performance on Novel Tasks. **a**. Learning curves for the first 100 exposures to held out tasks averaged over all tasks for models with five random initialization. Shaded region shows plus and minus one standard deviation across different random initializations. Stars indicate significant differences among performance on first exposure to novel task according to unequal variance t-test (for all subplots, n.s. = p>0.05, ∗= p<0.05, ∗∗= p<0.01, ∗∗∗= p< 0.001). A full table of pairwise comparisons can be found in Supplementary Info. Fig. 3. **b**. Distribution of generalization performance (i.e. first exposure to novel task) across models. **c**. Generalization performance for tasks where instructions are swapped at test time. Each point indicates average performance across tasks for a random initialization of model parameters. **d**. Generalization performance for models where tasks from the same family are held out during training. **e**. Generalization performance for models where the last layers of language models are allowed to fine-tune to the loss from sensorimotor tasks. **f**. Average difference in performance between tasks that use standard imperative instructions and those that use instructions with conditional clauses and require a simple deductive reasoning component. Colored stars on bottom show p-values for unequal variance t-test between a the null distribution constructed using random splits of the task set (transparent points), and stars at the top of plot indicate p-value results from t-test comparing difference amongst models (see Methods and Supplementary Info. Fig. 6. for full comparisons).

Our uninstructed control model simpleNet performs at 39% on average on the very first presentation of the novel tasks (0-shot generalization). This serves as a baseline for generalizing performance, as orthogonal task rules contain no information about how tasks relate to one another. Note that given how we score trials and the continuous nature of motor outputs (see Methods), a random network would effectively perform at 0% correct. However, despite the full orthogonality of task rules provided to simpleNet, exposure to the task set allows models to learn patterns that are common to all tasks (e.g. always repress response during fixation, always respond with a coherent angle etc.). In these cases, even though models have no knowledge about specific task demands they can nonetheless make better guesses. Therefore, 39% is not random level performance per se, but rather baseline performance achieved by the default response mode of a network trained and tested on a task set with some common requirements for responding (we also test a version of simpleNet with added layers between task-identifying information and the Sensorimotor-RNN and found that this added representational power fails to induce a statistically significant increase in performance (*t* = 0.58, *p >* 0.5, Supplementary Info. Fig. 4.5).

Next we tested generalization in networks that received linguistic instructions. gptNet, exhibits a 0-shot generalization of 57%. This is a significant improvement over simpleNet (*t* = 8.32, *p <* 0.001). Strikingly though, the relatively large size and extensive pre-training of gpt produces generalization performance which remains well below bowNet, which achieves a performance of 64% on unseen tasks (*t* = 3.34, *p <* 0.001). Perhaps even more striking is that increasing the size of gpt by an order of magnitude to the 1.5 billion parameters used by gptXL only results in modest gains over bowNet, with gptNet (XL) achieving 68% on held out tasks (*t* = 2.04, *p* = 0.047). By contrast, clipNet (S), which uses 4% the number of parameters utilized by gptNet (XL), is nonetheless able to achieve the same performance (68% correct, *t* = 0.146, *p* = 0.88). Likewise, bertNet achieves a generalization performance that lags only 2% behind gptNetXL in the mean (a difference which, again, was not statistically significant, *t* = 1.12, *p* = 0.262). Next we tested language models which learn explicitly about sentence level semantics during their pre-training. Specifically, sbert starts from the pre-trained weights of bert but also learns from an additional supervised linguistic objective designed to produce a high quality sentence embedding space. [40, 41]. sbertNet can perform an unseen task at 79% correct on average. Finally, our best performing model, sbertNet (L), can execute a never before seen task with a performance of 83% correct on average, lagging just a few percentage points behind structureNet (88% correct), which receives the structure of the task set hand-coded in its rule vectors.

Fig. 2b. shows a histogram of the number of tasks for which each model achieves a given level of performance. Again, sbertNet (L) manages to perform over 20 tasks set nearly perfectly in the zero-shot setting (for individual task performance for all models across tasks see Supplementary Info. Fig. 3). To validate that our best performing models truly leveraged the semantics of instructions to generalization performance, we presented the network with the sensory input for one held out task, while providing the linguistic instructions for a different held out task. Models that truly infer the proper sensorimotor response based on the semantic content of instructions should be most penalized by this manipulation. As predicted, swapping instructions destroys the capacity to generalize in all of our instruction models (Fig. 2c). We also test a more stringent hold-out procedure where we purposefully choose 4-6 tasks from the same family of tasks to hold-out during training, forcing models to infer an entire set of interrelated sensorimotor mappings from language instructions (Fig. 2d). Overall, performance decreases in this more difficult setting, though our best performing models still shows strong generalization, with sbertNet (L) and sbertNet achieving 71% and 72% correct on novel tasks, which was not significantly different from structureNet at 72% (*t* = 0.629, *p* = 0.529; *t* = 0.064, *p* = 0.948 for sbertNet (L) and sbertNet respectively). To test whether a rudimentary form of linguistic grounding [43] can improve performance we allow the weights of language models to tune according to the loss experienced during sensorimotor training (see Methods for tuning details). This manipulation improves the generalization performance across all models, and for our best performing model sbertNet (L) we see that generalization is as strong as for structureNet (86%, *t* = 1.204, *p* = 0.229). Finally, following [31] we also tested models in a setting where task type information for a given task was represented as a composition of information for related tasks in the training set (e.g. AntiDMMod1 = (*rule*(AntiDMMod2) − *rule*(DMMod2)) + *rule*(DMMod1)). In this setting we did find that the performance of simpleNet improved (60% correct). However, when we embed task instructions using pre-trained language models and combine these embeddings according to the same compositional rules, our linguistic models dramatically outperformed simpleNet. This suggests that training in the context of language can lead to an organization of computational components which more readily allows a simple compositional scheme to successfully configure task responses (see Supplementary Info. 5 for full results and compositional encodings).

The discrepancy in performance between our instructed models across all these conditions is striking given that each model used to embed instructions exhibits impressive language skills in their own right. This suggests that in order to represent linguistic information such that it can successfully configure sensorimotor networks, it isn’t sufficient to simply use any very powerful language processing system. Rather the problem of generalizing based on instructions requires a specific type of linguistic processing. For our models, success can be delineated by examining the extent to which they are exposed to knowledge about sentence-level semantics in their linguistic pre-training. Our best performing models sbertNet (L) and sbertNet are explicitly trained to produce good sentence embeddings, whereas our worst performing model gptNet is only tuned to the statistics of upcoming words. Both clipNet (S) and bertNet, are exposed to some form of sentence level knowledge, though not in the supervised sense that sbert models are. clipNet (S) is interested in sentence-level representations, but trains these representations using the statistics of corresponding vision representations. bertNet performs a two-way classification of whether or not input sentences are adjacent in the training corpus, though it is primarily involved in predicting masked words. That the 1.5B parameters of gptNet (XL) doesn’t allow it to separate its performance from these comparatively small models speaks to the fact that model size isn’t the determining factor. Rather, what allows models to process instructions in a way that induces generalization is the extent to which they possess explicit knowledge of sentence-level semantics, and can organize their instruction embeddings according to those semantics. Indeed, though Bag of Words removes key elements of linguistic meaning (e.g. syntax), and is ultimately a crude way to represent language, the simple use of word occurrences results in an embedding space where the information that is encoded using this method is primarily information about the similarities and differences between the sentences. For instance, simply representing the inclusion or exclusion of the words ‘stronger’ or ‘weaker’ is highly informative about the meaning of the instruction. This results in some degree of generalization for bowNet.

With our set up we can also begin to explore which features of language make it difficult for our models to configure the proper sensorimotor response. 30 of our tasks require processing instructions with a conditional clause structure (e.g. ‘COMP1’) as opposed to a simple imperative (e.g. ‘AntiDM’). Importantly, tasks that are instructed using conditional clauses also require a simple form of deductive reasoning in order to formulate a correct response (if p then q else s). Neuroimaging literature exploring the relationship between such deductive processes and language areas have reached differing conclusions, with some early studies showing that deduction recruits regions that are canonically thought to support syntactic computations [44, 45, 46], and follow-up studies claiming that deduction can be reliably dissociated from language areas [47, 48, 49, 50]. One theory for this variation in results is that baseline tasks used to isolate deductive reasoning in earlier studies used linguistic stimuli that required only superficial processing [51, 52].

To explore this issue, we calculated the average difference in model performance between tasks with and without conditional clauses/deductive reasoning requirements (Fig. 2f). All our models performed worse on these tasks relative to a set of random shuffles. Interestingly, however, we also saw an additional effect between structureNet, and our instructed models, which performed worse than structureNet by a statistically significant margin (see Supplementary Info. 6 for full comparisons). This is a crucial comparison because structureNet performs deductive tasks without relying on language. Hence, the decrease in performance between structureNet and instructed models strongly suggests that some of the total effect we see in instructed models is due to the difficulty inherent in parsing syntactically more complicated language. The implication is that we may indeed see increased engagement of linguistic areas in deductive reasoning tasks, but this may simply be due to the increased syntactic demands of corresponding instructions (rather than processes that recruit linguistic areas to explicitly aid in the deduction), and hence controlling for syntactic complexity is key to isolating deductive processes. This result largely agrees with two reviews of the deductive reasoning literature which concluded that the effects in language areas seen in early studies were likely due to the syntactic complexity of test stimuli [51, 52].

Next, we move to an analysis of the instruction embedding spaces produced by different language models, the resultant structure in Sensorimotor-RNN activations across different tasks, and how shared geometry between both these spaces aids in generalization.

### Shared structure in language and sensorimotor networks support generalization

In order to assess what properties of language embeddings lead to strong generalization across sensorimotor tasks, we first examined the representations that emerge in Sensorimotor-RNNs and in the final layer of our language models across different tasks.

First, we note that like in other multitasking models, units in our Sensorimotor-RNNs exhibit functional clustering, where similar subsets of neurons show high variance across similar sets of tasks (Supplementary Info. Fig. 7). Moreover, we find that models can learn unseen tasks by only training Sensorimotor-RNN input weights and keeping the recurrent dynamics constant (Supplementary Info. Fig. 8). Past work has shown that clustering and the ability to recruit new skills while recurrent dynamics are fixed are characteristic of networks that are able to factorize their computations and reuse the same set of underlying neural resources across different settings [31, 10].

We then examined the geometry that exists between the neural representations of related tasks. We plotted the first 3 PCs of Sensorimotor-RNN hidden activity at stimulus onset in simpleNet, gptNetXL, sbertNet (L), and structureNet performing modality specific DM and AntiDM tasks. Here models receive input for a decision-making task in both modalities but must only account for the stimuli in the modality relevant for the current task. Importantly, AntiDMMod1 is held out of training in the following examples. We observe that a pattern of abstract relations emerges among tasks depending on how instructions are embedded, and afterwards offer a quantitative analysis of this abstract structure across all held out tasks and all models. In addition, we plot the PCs of either the rule-vectors or the instruction embeddings in each task. Results are shown in Fig. 3.

**Figure 3.**
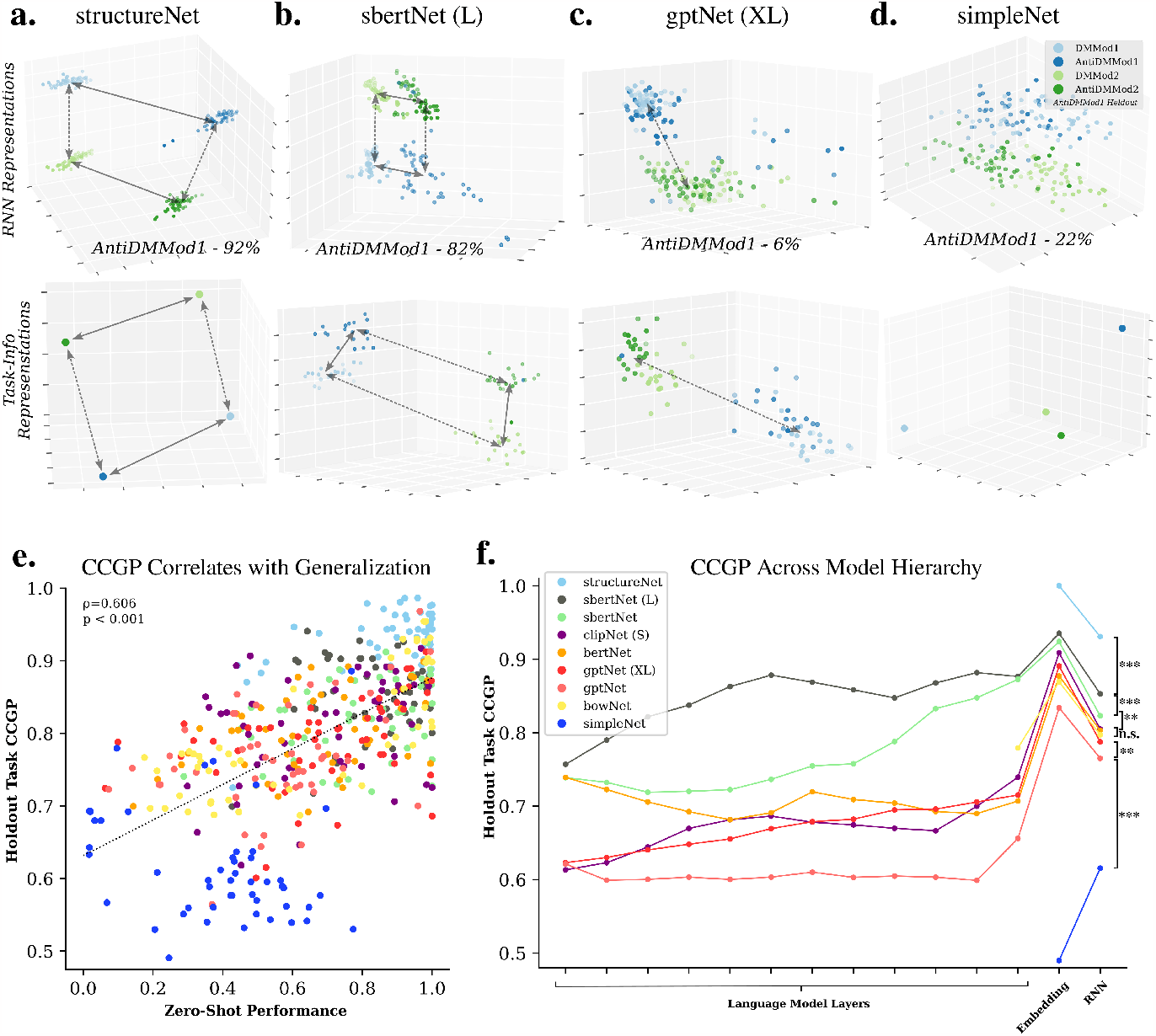
Structured Representations in Instructed Models. **a.-d**. First three PC’s of sensorimotor hidden activity and task-info representations for models trained with AntiDMMod1 held out. Solid arrows represent an abstract ‘Pro’ vs. ‘Anti’ axis, and dashed arrows represent an abstract ‘Mod1’ vs. ‘Mod2’ axis. **a**. structureNet **b**. sbertNet (L) **c**. gptNet (XL) **d**. simpleNet **e**. Correlation between held out task CCGP and zero-shot performance. **f**. CCGP scores for held out tasks for each layer in the model hierarchy. Significance scores indicate p-value results from unequal variance t-tests performed amongst models distributions of CCGP scores on held outs tasks for Sensorimotor-RNN (n.s. = p>0.05, ∗= p<0.05, ∗∗= p<0.01, ∗∗∗= p< 0.001; see Supplementary Info. 9 for full comparisons as well as t-test results for embedding layer CCGP scores).

For structureNet, we see that hidden activity is factorized along task relevant axes, namely a ‘Pro’ vs. ‘Anti’ direction in activity space that is consistent whether applied to ‘Mod1’ or ‘Mod2’ (solid arrows), and a ‘Mod1’ vs. ‘Mod2’ direction that is consistent across ‘Pro’ vs. ‘Anti’ conditions (dashed arrows). Importantly, this structure is maintained even for ‘AntiDMMod1’ which has been held out of training, allowing structureNet to achieve a performance of 92% correct on this unseen task. This factorization is also reflected in the PCs of rule embeddings. Overall, these are results we would expect from a model where the compositional structure of the task set is hand-coded in the rule vectors. Strikingly, sbertNet (L) also organizes its representations in a way that captures the essential compositional nature of the task set using only the structure that it has inferred from the semantics of instructions. This is the case for language embeddings, which maintain abstract axes across AntiDMMod1 instructions (again, held out of training) and instructions for the training task. As a result, sbertNet (L) is able to use these relevant axes for AntiDMMod1 Sensorimotor-RNN representations, leading to a generalization performance of 82%. By contrast, gptXLNet fails to properly infer a distinct ‘Pro’ vs. ‘Anti’ axes in either Sensorimotor-RNN representations or language embeddings, despite the fact that it uses a large and powerful language model to process instructions (Fig. 3b. Column 2). This results in a zero-shot performance on AntiDMMod1 of 6%. Finally, we find that although tasks in the training set are discriminable for simpleNet, its orthogonal rule vectors preclude it from leveraging any structured knowledge for the held out task, resulting in a performance of 22%.

To more precisely quantify the structure that may exist between representations of tasks and instructions across the entire task set and all models we measure the Cross Condition Generalization Performance (CCGP) of these representations [6]. CCGP measures the ability of a linear decoder trained to differentiate one set of conditions (e.g. ‘DMMod2’ and ‘AntiDMMod2’) to generalize to an analogous set of test conditions (e.g. ‘DMMod1’ and ‘AntiDMMod1’). Intuitively, this captures the extent to which models have learned to place sensorimotor activity along abstract task axes (e.g. the ‘Anti’ dimension). This measure serves as an important bridge between our models and representations in biological brains, as high CCGP scores and related measures have been observed in experiments that required human participants to flexibly switch between different interrelated tasks [7, 2]. In particular, single-cell recordings in MFC exhibit high CCGP for related task demands, which allows for rapid switching between different task types and output modalities [7].

Firstly, we measured CCGP scores amongst representations in Sensorimotor-RNNs for tasks that have been held out of training (see Methods for details). We found a strong correlation between tasks which display high CCGP and a models’ performance on that task in a zero-shot setting (Fig. 3d.). This strongly suggests that representing held out tasks in a way that preserves the geometry with tasks in the training set is a crucial factor aiding in generalization. We can verify this again using a task switching test. If language models’ ability to properly relate linguistic information contained in novel and practiced instructions is truly responsible for inducing the proper structure for held out task representations, then swapping instructions should destroy this structure and sharply reduce CCGP scores for Sensorimotor-RNNs. As expected, CCGP scores for all instructed models is dramatically reduced in this setting (Supplementary Info. Fig. 9).

Using our models not only can we examine how abstract representations emerge in Sensorimotor-RNNs responsible for executing stimulus-response mappings, but go a step further and ask to what extent these relations emerge in the language system responsible for processing instructions. CCGP decoding scores for different layers in our model are shown in Fig. 3d. For each instructed model, scores for 12 transformer layers (or the last 12 layers for sbertNet (L) and gptNet (XL)), the 64-dimensional embedding layer, and the Sensorimotor-RNN task representations are plotted. We also plot CCGP scores for the rule embeddings used in our non-linguistic models and find that structureNet and simpleNet achieve perfect CCGP and chance level performance respectively. Among models, there is a notable discrepancy in how abstract structure as measured by CCGP emerges in the language processing hierarchy. Autoregressive models (gptNetXL, gptNet), bertNet and clipNet (S), show a low CCGP throughout language model layers followed by a jump in CCGP in the embedding layer. This is predominantly because weights feeding into the embedding layer are tuned during sensorimotor training to ensure all Sensorimotor-RNNs receive task information as a 64-dimensional vector. The implication of this spike is that most of the useful representational processing that comes with embedding instructions using these models actually does not occur in the pre-trained language model per se, but rather in the linear readout which is exposed to task structure via training. By contrast, our best performing models sbertNet and sbertNet (L) employ language representations where high CCGP scores emerge gradually in the intermediate layers of their respective language models. Again, we emphasize the weights of language models are fixed during training on sensorimotor tasks. That the semantic representations obtained from sbertNet and sbertNet (L) pre-training are structured in a way that is also useful for organizing downstream sensorimotor dynamics is a striking result. Further, because semantic representations already have such a structure, most of the compositional inference involved in generalization can occur in the comparatively powerful language processing hierarchy. Since sbertNet and sbertNet (L) pre-training induces high CCGP throughout the intermediate stages of language processing, representations are already well organized in the last layer of language models, and a linear readout in the embedding layer is sufficient for the Sensorimotor-RNN to correctly infer the geometry of the task set and generalize well.

This analysis strongly suggests that models exhibiting generalization do so by relating practiced and novel tasks in a structured geometry that captures the relevant subcomponents of each task. Models are able to correctly infer these relations because they are guided by language models which themselves embed task instructions according to the same basic structure. Hence, successful models can leverage the linguistic relations between practiced and novel instruction sets to cue sensorimotor activity for an unseen task in the proper region of RNN activity space relative to this geometry, thereby composing practiced behaviors to perform well in a novel setting.

### Semantic modulation of single unit tuning properties

In the previous section we outlined the structured representational scheme that allows our best performing instructed models to generalize performance to unseen tasks. With our artificial models we can go a step further and examine tuning profiles of individual neurons in our Sensorimotor-RNNs, allowing us to make predictions for single-unit activity in human experiments. As one would expect, we found that individual neurons are tuned to a variety of task relevant variables, such as the position of the input in a ‘Go’ task, or the difference in intensity between the input stimuli in a decision making task. Critically, however, we find neurons where this tuning varies predictably within a task group and is modulated by the semantic content of instructions in a way that reflects task demands.

For instance, for the ‘Go’ family of tasks, unit 42 shows direction selectivity that shifts by *π* between ‘Pro’ and ‘Anti’ tasks, reflecting the relationship of task demands in each context (Fig. 4a). This flip in selectivity is observed even for the AntiGo task, which was held out during training. Hence, the observed reversal is induced solely by the linguistic instructions, with no additional training. Again, note that tuning curves are produced by the same set of stimuli, and the only factor responsible for the variation in tuning is the input instructions for the task. We make sure to use the full range of instructions for each task.

**Figure 4.**
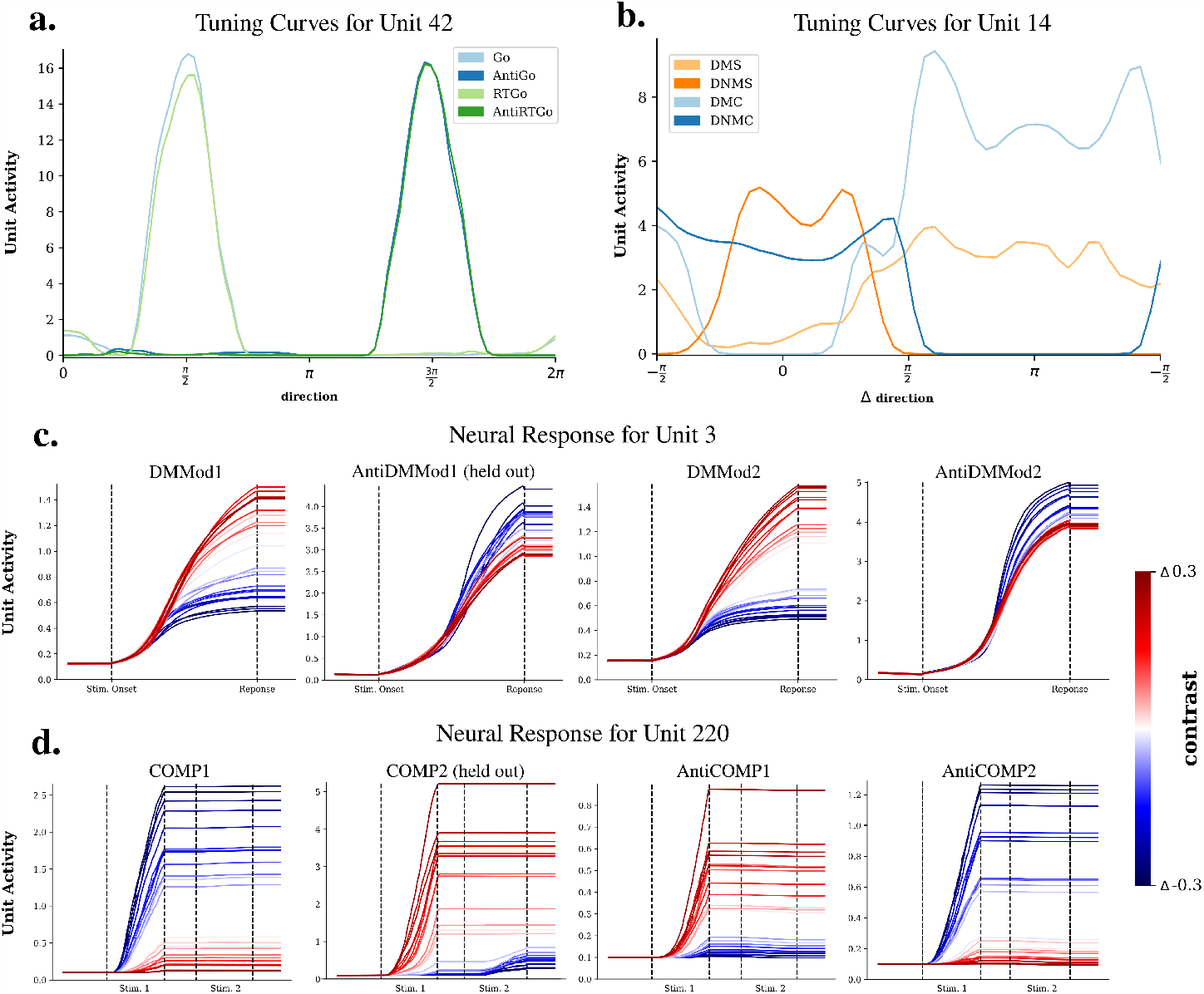
Semantic Modulation of Single Unit Tuning Properties. **a.)** Tuning curves for example sbertNet (L) Sensorimotor-RNN unit that modulates tuning according to task demands in the ‘Go’ family. **b.)** Tuning curves for example sbertNet (L) Sensorimotor-RNN unit in the ‘Matching’ family of tasks plotted in terms of difference in angle between two stimuli. **c)** Full activity traces for modality specific ‘DM’ and ‘AntiDM’ tasks for different levels of relative stimulus strength. **d)** Full activity traces for tasks in the ‘Comparison’ family of tasks for different levels of relative stimulus strength.

For the ‘Matching’ family of tasks, Unit 14 modulates activity between ‘Match’ (DMS, DMC) and ‘Non-Match’ (DNMS, DNMC) conditions. In ‘Non-Match’ trials the activity of this unit increases as the distance between the two stimuli increases. By contrast, for ‘Matching’ tasks this neuron is most active when the relative distance between the two stimuli is small. Hence, in both cases this neuron modulates its activity to represent when the model should respond, changing selectivity to reflect opposing task demands between ‘Match’ and ‘Non-match’ trials. Again this is true even for DMS which has been held out of training.

We can also examine how the content of instructions affects the temporal evolution of neural activity for single units across the relevant trial epochs. Fig. 4c. shows traces of Unit 3 activity in modality specific versions of DM and AntiDM tasks (AntiDMMod1 is held out of training) for different levels of contrast (*contrast* = *str*_*stim*1_ − *str*_*stim*2_). In all tasks, we observe ramping activity where the rate of ramping is relative to the strength of evidence (in this case, contrast). This motif of activity has been reported in various previous studies [53, 54]. However, in our models we observe that an evidence accumulating neuron can swap the sign of its integration in response to a change in linguistic instructions, which allows models to meet opposing demands of ‘Standard’ and ‘Anti’ versions of the task. Moreover, this modulation of neural tuning is accomplished in a 0-shot setting, when the pre-synaptic weights of the neuron have never explicitly tuned to the AntiDMMod1 task.

Finally, we see a similar pattern in the time course of activity for trials in the ‘Comparison’ family of tasks (Fig. 4d.). These tasks add a level of complexity to the classic ‘Decision-Making’ tasks; in the COMP1 task, the network must respond in the direction of the first stimulus if it has higher intensity than the second stimulus, and must not respond otherwise. In COMP2, it must respond to the second stimulus if the second stimulus is higher intensity, and not respond otherwise. For ‘Anti’ versions, the demands of stimulus ordering are the same except the model has to choose the stimuli with the weakest contrast. The added syntactic complexity makes these tasks particularly demanding. Yet we still find individual neurons that modulate their tuning with respect to task demands, even for held out tasks (in this case ‘COMP2’). For example unit 82, is active when the network should repress response. For ‘COMP1’ this unit is highly active with negative contrast (i.e. *str*_*stim*2_ *> str*_*stim*1_), but flips this sensitivity in ‘COMP2’ and is highly active with positive contrast (i.e. *str*_*stim*1_ *> str*_*stim*2_). Importantly, this relation is reversed when the goal is to select the weakest stimuli, in which case the unit shows high activity for a relatively strong versus a relatively weak first stimulus in ‘AntiCOMP1’ and ‘AntiCOMP2’ tasks respectively. Hence, despite these subtle syntactic differences in instruction sets, the language embedding can reverse the tuning of this unit in a task-appropriate manner.

### Linguistic communication between networks

So far we have offered an account of what representations emerge in a network that understands the sensorimotor implications of linguistic instructions. We now seek to model the complementary human ability to describe a particular sensorimotor skill with words once it has been acquired. To do this we invert the language-to-sensorimotor mapping our models learn during training so that they can provide a linguistic description of a task based only on the state of sensorimotor units. First, we construct an output channel, which we call the Production-RNN (Fig. 5a-c; blue), to map the information contained in the Sensorimotor-RNN to the linguistic instructions. To train this network, we used a self-supervised procedure: when the network is presented with sensory inputs and linguistic instructions that drive sensorimotor activity, these same instructions are used as targets to train the Production-RNN (Fig. 5a, see Methods for training details). We then pose the following problem to our networks: after training, suppose the network is tested on a sequence of trials in which it is presented with a sensory input and the correct motor feedback for a task, but without the linguistic instructions that normally provide the essential information about task demands. Our goal is to determine whether models can use the information in the motor feedback signal to provide an appropriate linguistic instruction for the task and verify that these instructions can effectively guide the performance of another network. In order to drive networks to perform a task in the absence of instructions, we begin by randomly initializing activity in the embedding layer of the model. We then present the network with a series of example trials for a specific task along with motor feedback and update embedding layer activity in order to reduce the error between the output of the model and the motor feedback signal (Fig. 5b.). All model weights remain frozen during this procedure. Once the activity in the embedding layer drives sensorimotor units to achieve a performance criteria over the series of example trials, we use the Production-RNN to decode a linguistic description of the current task. Finally, to evaluate the quality of these instructions we input them to a partner model and measure partner model performance across tasks with produced instructions (Fig. 5c.). All instructing and partner models used in this section are instances of sbertNet (L) (see Methods for details).

**Figure 5.**
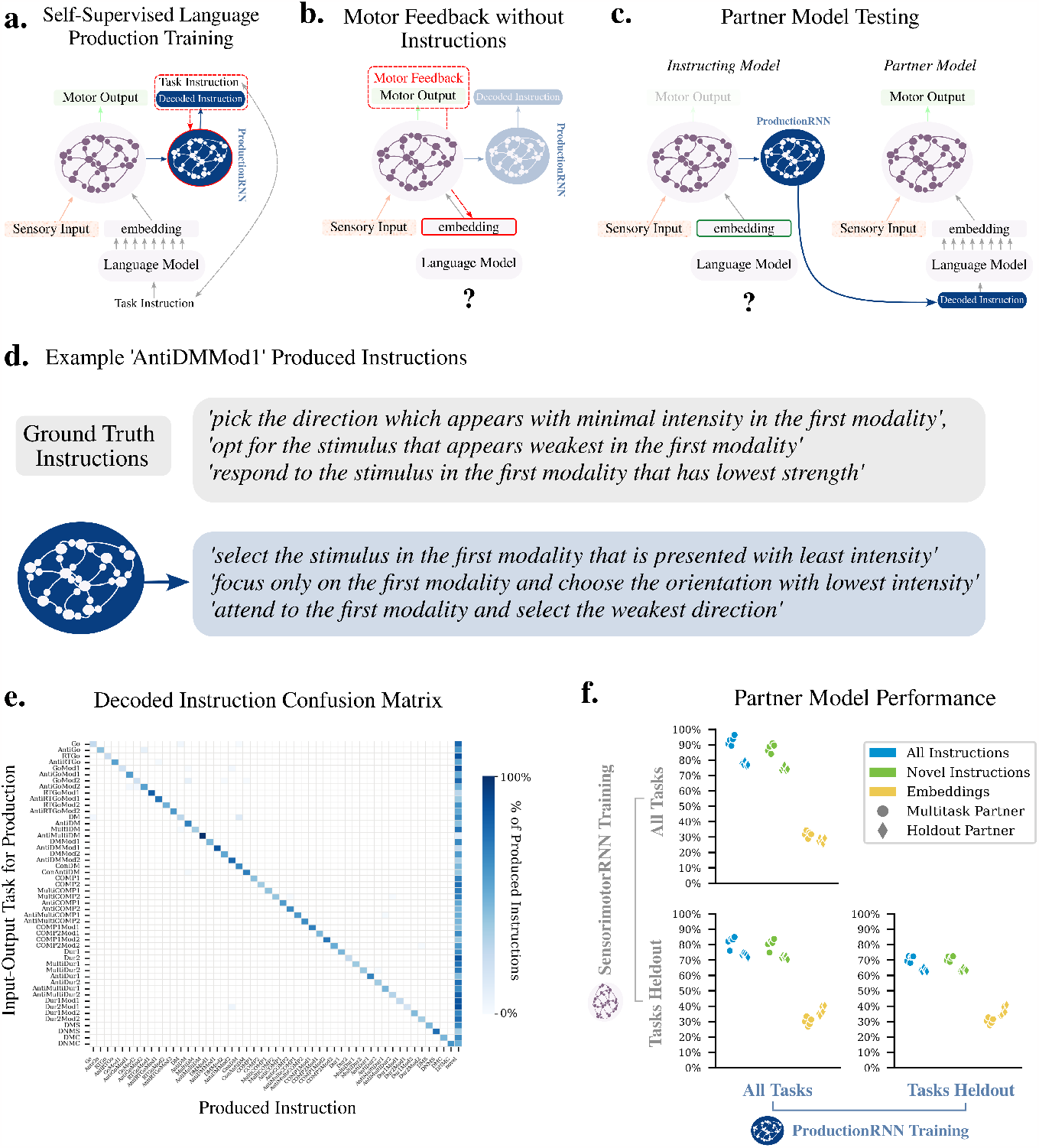
Communication Between Networks. **a**. Illustration of self-supervised training procedure for the language production network (blue). Red dotted line indicates gradient flow. **b**. Illustration of motor feedback used to drive task performance in the absence of linguistic instructions. **c**. Illustration of the partner model evaluation procedure used to evaluate the quality of instructions generated from the instructing model **d**. Three example instructions produced from sensorimotor activity evoked by embeddings inferred in **b** for an AntiDMMod1 task **e**. Confusion matrix of instructions produced again using the method described in **b. f**. Performance of partner models in different training regimes given produced instructions or direct input of embedding vectors. Each point represents the average performance of a partner model across tasks using instructions from a decoders train with different random initializations. Dots indicate the partner model was trained on all tasks whereas diamonds indicate performance on held out tasks. Axes indicate the training regime of the instructing model. Full statistical comparisons of performance can be found in Supplementary Info. 11.3.1

First, we show some example decoded instructions for the AntiDMMod1 task (Fig. 5d, see Supplementary Info. 15 for all decoded instructions). To visualize decoded instructions across the task set we plot a confusion matrix where both Sensorimotor-RNN and Production-RNN are trained on all tasks (Fig. 5e). On the y-axis is the task that determines the input-output pairs used to infer an embedding as depicted in Fig. 5b. The x-axis indicates whether the instruction produced from the resulting sensorimotor activity was included in one of the instruction sets used during self-supervised training or was a ‘Novel’ instruction formulation i.e. a decoded instruction that was not included in the training set for the Production-RNN (see Methods for details). Darker tones indicate a higher overall percentage of produced instruction fell into a particular category. Novel instructions make up 53% of decoded instructions across all tasks. Of course, the novelty of these instructions says nothing about how well they convey information about the current task. To test this, we evaluate a partner model’s performance on instructions that the first network has generated by inferring task demands and communicating these demands via a linguistic output (Fig. 5c.). Results are shown in Fig. 5f, top left.

When the partner model is trained on all tasks, performance on all decoded instructions is 93% on average across tasks (blue dots). Communicating instructions to partner models with tasks held out of training also results in good performance (78%) meaning that partner models can generalize to unseen tasks based on produced instructions (blue diamonds). Importantly, this information is present even for ‘novel’ instructions, where average performance is 88% for partner models trained on all tasks and 75% for partner models with hold-out tasks (green bars).

Given that the instructing and partner models share the same architecture, one might expect that it is more efficient for the two networks to use direct neural communication, instead of language-based communication. For instance, we can forgo the language component of communication altogether and simply copy the embedding inferred by one model into the input of the partner model. This results in only 31% correct performance on average, compared to 93% for linguistic communication, and 28% performance when testing partner networks on tasks that have been held out of training (yellow marks). The reason for this drop off is straightforward: though both instructing and partner networks share the same basic architecture and the same task competencies, they nonetheless have different synaptic weights. Hence, using a neural representation tuned for the set of weights within the one agent won’t necessarily produce good performance in the other.

We also want to determine the extent to which our models can leverage their ability to generalize to unseen tasks in order to produce useful instructions for novel tasks, again only using information gained from motor feedback. We begin with an instructing model Sensorimotor-RNN with tasks held out of training, after which we freeze the weights in this network and expose the system to example trials of all tasks to train the Production-RNN. We emphasize here that during training the Production-RNN attempts to decode from sensorimotor hidden states induced by instructions for tasks the network has never experienced before (as in Fig 5a), whereas during test time instructions are produced from sensorimotor states that emerge entirely as a results of minimizing a motor error (as in Fig. 5b,c). This contrast in train and test conditions makes the task of decoding non-trivial. Yet, we nonetheless find that such a system can produce quality instructions even from tasks which the Sensorimotor-RNN was never exposed to during training (Fig 5.f bottom left). In this setting a partner model trained on all tasks performs at 82% correct across tasks, while partner models with tasks held out of training perform at 73%. Here 77% of produced instructions are novel so we see a very small decrease of 1% when we test the same partner models only on novel instructions. Like above, context representations induce a relatively low performance of 30% and 37% correct for partners trained on all tasks and with tasks held out respectively.

Lastly, we test our most extreme setting where tasks have been held out for both Sensorimotor-RNNs and Production-RNNs (Fig. 5f., bottom right). Here again we find that produced instructions induce a performance of 71% and 63% for partner models trained on all tasks and with tasks held out respectively. Though this is a decrease in performance from our previous set-ups, the fact that models can produce sensible instructions at all in this double held out setting is striking. Previously we showed that the production network could produce valid, novel instructions for tasks where instructions for that task were included in the training set (Fig 5f. first column, green bars). Here we go a step further and ask the system to infer the proper combination and ordering of words such that the resulting sentence effectively explains a task which the system itself has never seen nor tried to describe before, and again, to produce this instruction in a setting where the only information about task demands comes in the form of motor feedback. The fact that the system succeeds to any extent in this setting speaks to strong inductive biases introduced by training the Sensorimotor-RNNs in the context of rich interrelations in and amongst instructed tasks.

## Discussion

In this study we use the latest advances in natural language processing to build tractable models of some quintessential human language skills: the ability to interpret instructions to guide actions in novel settings and the ability to produce a description of a task once it has been learned. We have shown that recurrent neural networks can learn to perform a set of psychophysical tasks simultaneously using a pre-trained language transformer to embed a natural language instruction for the current task. Our best performing models can leverage these embeddings to perform a brand new task on the very first trial, with an average performance of 83% correct based solely on a linguistic description. Instructed models that generalize performance do so by leveraging the shared structure of instruction embeddings and tasks representations, such that an inference about the relations between practiced and novel instructions leads to a good inference about what sensorimotor transformation is required for the unseen task. Finally, once a network understands how to interpret instructions to perform a sensorimotor task, it can invert this process and provide a linguistic description for a previously unseen task based only on the sensorimotor contingency it observes, in the absence of any linguistic input. We find that communicating through language, as opposed to directly via latent neural representations, is essential for effectively passing information between independent agents each with their own set of synaptic weights.

Our models make several predictions for what neural representations to expect in brain areas that integrate linguistic information in order to exert control over sensorimotor areas. Our first prediction comes from the CCGP analysis of our model hierarchy, which suggests that when human subjects must generalize across (or switch between) a set of related tasks based on instructions, the neural geometry observed amongst sensorimotor mappings should also be present in semantic representations of instructions that emerge from the language processing hierarchy. This prediction is well grounded in the existing experimental literature where multiple studies have observed the type of abstract structure we find in our Sensorimotor-RNNs also exists in sensorimotor areas of biological brains [6, 55, 56, 2]. In Sensorimotor-RNNs that were best able to maintain this structure in a novel setting, we found the analogous structure emerged from language modules responsible for inferring task demands based on linguistic information. Hence, we predict that the emergence of this task related structure in language areas is an essential component of instructed action in humans, and should guide experiments examining the interface between language processing and sensorimotor control. One intriguing candidate for an area that may support this structured embedding of instructions is the language selective subregion of left Inferior Frontal Gyrus (lIFG). This area is sensitive to both lexico-semantic and syntactic aspects of sentence comprehension, is implicated in tasks that require semantic control (e.g. a match to sample task where the participants must judge if the meaning of stimuli are related), indicating that representations in this area can contribute to decision-making processes, and most intriguingly, lies anatomically adjacent to another functional subregion of lIFG which is part of the Multiple Demands Network, a circuit responsible for executive control and flexible cognition [57, 58, 59, 60]. We also predict that individual units involved in implementing sensorimotor mappings should modulate their tuning properties on a trial by trial basis according to the semantics of the input instructions. This theoretical insight may be especially useful to interpret high resolution imaging data and multi-unit recordings in humans, our only example of organisms with sophisticated linguistic competencies. Finally, given that grounding linguistic knowledge in the sensorimotor demands of the task set improved performance across models (Fig. 2e), we predict that during learning the highest level of the language processing hierarchy should likewise be shaped by the embodied processes that accompany linguistic inputs, for example motor planning or affordance evaluation [61].

One notable negative result of our study is the relatively poor generalization performance of gptNet and, in particular, gptNetXL, which used at least an order of magnitude more parameters than other models to process instructions. This is particularly striking given that activity in these models is predictive of many behavioral and neural signatures of human language processing in both passive listening and reading tasks [19, 20]. These studies argue that there is an essential predictive component of word-level language processing and that these computations are captured by autoregressive language models. Nonetheless, embedding instructions with these systems failed to engender the characteristic human ability to generalize based on instructions, and also didn’t produce the structured language embeddings that we found in our best performing models. Given this, a synthesis of these results may lead to a holistic understanding of the entire language hierarchy. For example, future imaging studies may be guided by the representations in both autoregressive models and our best performing models to more precisely delineate a full gradient of brain areas involved in each stage of language processing, from low-level next word prediction to higher level structured sentence representations to the sensorimotor control that language informs.

In addition to shedding light on the unique human ability to generalize based on instructions, our results also open the possibility of future work comparing flexible cognition and compositional representations in non-linguistic subjects like non-human primates (NHPs). Comparison of task switching (without linguistic instructions) between humans and NHPs do indicate that both use abstract rule representations, although humans are able to make switches much more rapidly [62]. One intriguing parallel in our analyses is the use of compositional rules vectors (Supplementary Info. 5). Even in the case of non-linguistic simpleNet, using these vectors boosted performance in unseen tasks, which suggests that simply learning the task set engenders agents with some notion of abstract rules even in the absence of language. Importantly, however, this compositionality is much stronger for our best performing instructed models which essentially recover all of the generalization performance seen in instructed trials. This suggests that language endows agents with a much more flexible organization of task subcomponenents which can be readily combined in a broader variety of contexts.

Our results also highlight the advantages of linguistic communication. Our networks can compress the information they have gained through experience of motor feedback and transfer that knowledge to a partner network such that the partner can forgo this learning process and simply follow a given instruction to perform the task. Further, they can communicate knowledge to a partner network even for tasks that the instructing network itself has never previously experienced. Though rudimentary in our example, the ability to endogenously produce a description of how to accomplish a task after a period of practice (motor feedback, in our case) is a hallmark human language skill and underlies much of the utility of linguistic communication in a community of speakers. The failure to transfer performance by simply sharing latent representations demonstrates that in order to effectively communicate information in a group of otherwise independent networks of neurons, it needs to pass through a representational medium that is equally interpretable by all members of the group. In humans and for our best performing instructed models, this medium is language.

A series of works in reinforcement learning has investigated using language and language-like schemes to aid agent performance. Agents receive language information in a variety of ways such as through step-by-step descriptions of action sequences [63, 64, 65, 66, 67], learning policies conditioned on a language goal [68, 69, 70, 71], by shaping a reward function given language inputs [72, 73] or by leveraging the predictive power of pre-trained models to aid directly in planning and action selection [74, 75]. These studies often deviate from natural language and receive linguistic inputs that are parsed or simply refer to environmental objects, or conversely output actions in the form of a language which is hand coded to interface with the action space of the agent’s virtual environment. Further, unlike our models, none of these agents demonstrate any clear connection to the experimental literature, and their task environments make them poor candidates to develop testable neural predictions. Some larger versions of the pre-trained language models we use to embed instructions also display instruction following behavior in their own domains e.g. GPT-3, PALM [21], LaMDA[22], [15], and InstructGPT [16] in the modality of language and DALL-E and Stable Diffusion [23] in a language to image modality. The semantic and syntactic understanding displayed in these models is impressive. However, the output of these models are difficult to interpret in terms of a motor output and so it is again difficult to formulate the computation used in these models as a prediction about the dynamics of a sensorimotor system tuned to interpret language. DALL-E’s language to image mapping shows some cross-modal language understanding, but it has yet to be demonstrated that the model uses language as a genuine instruction as opposed to merely a prompt (for example, can the model infer control from language as in ‘draw a black dog or a blue fish but not both’?). Finally, recent work has sought to engineer instruction following agents that can function in complex or even real world environments[24, 25, 26]. While these models exhibit impressive behavioral repertoires, they rely on perceptual systems that fuse linguistic and visual information at very early stages in processing. This early fusing makes them difficult to compare to language representations in human brains, which emerge from a functionally and anatomically independent set of areas specialized for processing language. In all, none of these models offer a testable representational account of *how* language might be used to induce generalization over sensorimotor mappings in the brain.

Our models by contrast make tractable predictions for what population and single unit neural representations are required to support generalization and can guide future experimental work examining the interplay of linguistic and sensorimotor skills in humans. Many cognitive competencies are as of yet unaddressed by our task set and our instruction sets fall short of capturing every aspect of complexity exhibited in the full range of human language. However, by developing interpretable models that can both understand instructions as guiding a particular sensorimotor response, and communicate the results of sensorimotor learning as an intelligible linguistic instruction, we have begun to explain the power of language in encoding and transferring knowledge in networks of neurons.

## Methods

### Model Architecture

#### Sensorimotor-RNN

The base model architecture and task structure used in this paper follows [31]. All networks of sensorimotor units, denoted SensorimotorRNN are Gated Recurrent Units [77] using ReLU non-linearities with 256 hidden units each. Broadly, inputs to the networks are made up of two components 1) sensory inputs, *X*_*t*_ and 2) task identifying information, *I*_*t*_. We initialize hidden activity in the GRU as *h*^0^ ∈ ℝ ^256^ with values set to 0.1. All networks of sensorimotor units use the same hidden state initialization, so we omit *h*^0^ in network equations. At each time step, a read-out layer Linear_out_ decodes motor activity, *ŷ*_*t*_, from the activity of recurrent hidden units, *h*_*t*_, according to:

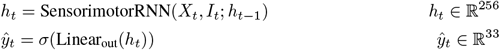

where *σ* denotes the sigmoid function. Sensory inputs *X*_*t*_ are made up of 3 channels, two sensory modalities *x*_mod1,*t*_ and *x*_mod2,*t*_, and a fixation channel *x*_fix,*t*_. Each of these inputs is continuously valued. Both *x*_mod1,*t*_, *x*_mod2,*t*_ ∈ ℝ ^32^ and stimuli in these modalities are represented as hills of activity with peaks determined by units’ preferred directions around a one-dimensional circular variable. Preferred directions for each of the 32 inputs units are evenly spaced between 0 and 2*π*. For an input at direction *θ*, the activity of a given input unit *u*_*i*_ with preferred direction *θ*_*i*_ is

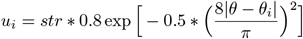

where *str* is the coefficient describing stimulus strength. The fixation channel *x*_fix,*t*_ ∈ ℝ ^1^ is a single unit simulating a fixation cue for the network. In all, sensory input *X*_*t*_ = (*x*_*mod*1,*t*_, *x*_*mod*2,*t*_, *x*_*fix,t*_) ∈ ℝ ^65^. Accordingly, motor output, *ŷ*_*t*_ consists of both a 32-dimensional ring representing directional responses to the input stimulus as well as a single unit representing model fixation, so that *ŷ*_*t*_ ∈ ℝ ^33^

For all models, task identifying information *I*_*t*_ ∈ ℝ^64^. Further, task identifying information is presented throughout the duration of a trial and remains constant such that *I*_*t*_ = *I*_*t′*_ ∀*t, t*′ . For all models, task identifying info *I*_*t*_ and sensory input *X*_*t*_ are concatenated as inputs to the SensorimotorRNN, so that the input dimensionality of all recurrent networks is 129.

#### Non-Linguistic Models

For simpleNet, we generate a set of 64-dimensional orthogonal task rules by constructing an orthogonal matrix using the Python package scipy.stats.ortho_group, and assign rows of this matrix to each task type. This results in a set of 50 orthogonal 64-dimensional vectors uniquely associated with each task type, which are used by simpleNet as task identifying information in our simulations. For structureNet we generate a set of 10 orthogonal, 64-dimensional vectors in the same manner, and each of these represents a dimension of the task set (e.g. respond weakest vs. strongest direction, respond in the same vs. opposite direction, pay attention only to stimuli in the first modality etc.). Rule vectors for tasks are simple combinations of each of these 10 basis vectors. For a full description of structure rule vectors see Supplementary Info. 14.

In order to assess whether added representational power might increase generalization in these models, we test simpleNetPlus and structureNetPlus, which use an additional hidden layer with 128 units and ReLU non-linearities to process orthogonal tasks rules *I*_*t*_ into a vector *Ī* _*t*_ which is used by SensorimotorRNN as task identifying information.

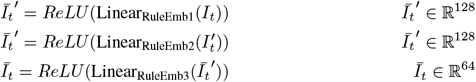

Results for these models are included in Supplementary Info. 4.5.

#### Pre-trained Transformers

For our instructed models, task identifying information is computed by embedding natural language instructions. The main language modules we test use pre-trained transformer architectures to produce *I*.

Importantly, transformers differ in the type of pre-training objective used to tune the model parameters and therefore also differ in the types of linguistic representations each model possesses. gpt is trained on a traditional language modeling objective i.e. predict next word given a context of words [18]. gptXL follows the same objective but trains for longer on a larger dataset [38]. Both models are fully autoregressive. bert, by contrast, takes bi-directional language inputs (future and past words) and is tasked with predicting masked words that appear in the middle of input phrases. Additionally, bert is trained on a simple sentence prediction task where the model must determine if input sentence 1 is followed by input sentence 2 in the training corpus. This training endows bert with some rudimentary notion of sentence level semantics. Extending this principle, sbert is explicitly trained to produce fixed length embeddings of whole sentences [40]. It takes pre-trained bert networks and uses them in a siamese architecture [76] which allows the weights of the model to be tuned in a supervised fashion according to the Stanford Natural Language Inference dataset [41]. Natural language inference is a three-way categorization task where the network must infer the logical relationship between sentences: whether a premise sentence implies, contradicts, or is unrelated to a hypothesis sentence. The sentence representations produced by these networks have been shown to match human judgements of sentence similarity across several datasets. Finally, CLIP is trained to jointly embed images and language [42]. It uses data from captioned images and is asked to properly categorize which text and images pairs match or are mismatched in the dataset via a contrastive loss. Hence, CLIP simultaneously shapes the language and image embedding spaces in complementary ways. For our models, we discard the image recognition network and simply use the language embedder.

Importantly, the natural output of a transformer is a matrix of size dim_trans._ × 𝒯, the inherent dimensionality of the transformer by the length of the input sequence. In order to create an embedding space for sentences it is standard practice to apply a pooling method to the transformer output which produces a fixed length representation for each instruction.

For gpt, gptXL, bert, and sbert we use an average pooling method. Suppose we have an input instruction *w*_1_ … *w* _*𝒯*_ . Following standard practice with pre-trained language models, the input to our transformers are tokenized with special ‘[cls]’ and ‘[eos]’ tokens at the beginning and end of the input sequence. We then compute *I* as follows,

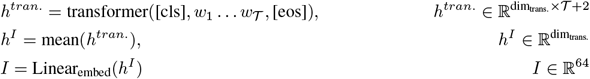

We chose this average pooling method for several reasons. Firstly, [40] found that average pooling resulted in highest performing sbert embeddings compared to other pooling methods. Said another way, average pooling produced the most effective sentence embeddings when fine-tuning pre-trained bert architectures on downstream tasks. Hence, we use averaged pooling for both sbert and bert models.

Another alternative would be to simply use the finally hidden representation of the [cls] token as a summary of the information in the entire sequence (given that bert architectures are bidirectional, this token will have access to the whole sequence).

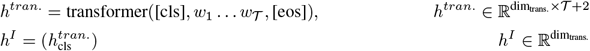

Where 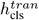denote the last hidden representation for the [cls] token. [40] found this pooling method performed worse than average pooling, so we don’t include these alternatives in our results. For gpt and gptXL we also tested a pooling method where the fixed length representation for a sequence was taken from the transformer output of the [eos] token, which has access to all the information in the input sequence for these fully autoregressive models. In this case

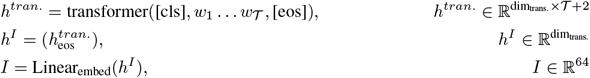

We found that gpt failed to achieve even a relaxed performance criteria of 85% across tasks using this pooling method and gptXL performed worse than with average pooling (Supplementary Info. 10). For clip models we use the same pooling method as in the original multi-modal training procedure which takes the outputs of the [cls] token as described above.

For all the above models we also test a version where the information from the pre-trained transformers is passed through an Multi-Layer Perceptron (MLP) with a single hidden layer of 256 hidden units and ReLU nonlinearities. We found that this manipulation reduced performance across all models, verifying that a simple linear embedding is beneficial to generalization performance. Because of the reduction in performance we omit these models from the main text and analyses (Supplementary Info. 4.4).

For gptNet, bert, and sbert, dim_trans._ = 768 and each model uses a total of ∼ 100M parameters; for sbert (L) dim_trans._ = 1024 and the model uses ∼ 300M parameters; gptNetXL dim_trans._ = 1600 and the model uses ∼ 1.5B parameters; for CLIP, dim_trans._ = 512 and model uses ~60M parameters. Full PyTorch implementations and model details can be found at the Huggingface library [79].

### BoW Model

We also use a Bag of Words embedding to test a comparatively simple language processing scheme. Here, instructions are represented as a vector of binary activations the size of the instruction vocabulary, where each unit indicates the inclusion or exclusion of the associated vocabulary word in the current instruction. For our instruction set |*vocab*| = 181. This vector is then projected through a linear layer into 64-dimensional space.

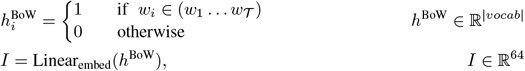

#### Blank Slate Language Models

Given that tuning the last layers of language models resulted in improved performance (Fig. 2e) we tested two additional models to determine if training a blank slate language model exclusively on the loss from sensorimotor tasks would improve performance. These models consist of passing BoW representations through an MLP and passing pre-trained BERT word embeddings through one layer of a randomly initialized BERT encoder. Both models performed poorly compared to pre-trained models (Supplementary Info. 4.5), confirming that language pre-training is essential to generalization.

### Tasks Sets

Tasks were divided into 5 interrelated subgroups: the ‘Go’ group, the ‘Decision-Making’ group, the ‘Matching’ group, and the ‘Comparison’ group and the ‘Duration’ group. Each group possesses its own internal structure which relates tasks to one another. Depending on the task, multiple stimuli may appear over the course of the stimulus epoch. Also depending on the task, models may be required to respond in a particular direction or repress response altogether. Unless otherwise specified, 0-mean Gaussian noise is added independently at each time step and to each input unit and the variance of this noise is drawn randomly from 𝕌 [0.1, 0.15]. The timing of stimuli differ among the tasks type. However, for all tasks trials can be divided into a preparatory, stimulus, and response epoch. The stimulus epoch can further be divided into three parts, stim1, delay and stim2, although these distinct parts aren’t used by all tasks. A trial lasts for a total of *T* = 150 times steps. Let *dur*_*epoch*_ denote the duration in simulated time steps of a given epoch. Then

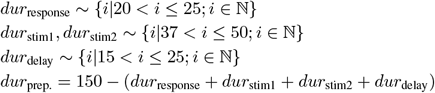

For tasks that don’t utilize a delay structure stim1, stim2, and delay epochs are grouped together in a single stimulus epoch where *dur*_stimulus_ = *dur*_stim1_ + *dur*_stim2_ + *dur*_delay_. Unless otherwise specified a fixation cue with a constant strength *str*_fix_ = 1 is activated throughout the prep. and stimulus epochs. For example trials of each task see Supplementary Info. 12.

#### ‘Go’ Tasks

The ‘Go’ family of tasks includes ‘Go’, ‘RTGo’, ‘AntiGo’, ‘AntiRTGo’ and modality specific versions of each task denoted with either ‘Mod1’ and ‘Mod2.’ In both the ‘Go’ and ‘AntiGo’ tasks, a single stimulus is presented at the beginning of the stimulus epoch. The direction of the presented stimulus is generated by drawing from a uniform distribution between 0 and 2*π*, i.e. *θ*_stim_ ∼ 𝕌 [0, 2*π*]. The stimulus will appear in either modality 1 or modality 2 with equal probability. The strength of the stimulus is given by *str*_stim_ ∼ 𝕌 [1.0, 1.2]. In the ‘Go’ task the target response is in the same direction as the presented stimulus i.e. *θ*_stim_ = *θ*_target_, while in the ‘Anti Go’ task the direction of the response should be in the opposite of the stimulus direction, *θ*_stim_ + *π* = *θ*_target_. For modality specific versions of each task, a stimulus direction is drawn in each modality *θ*_stim, mod1_ ∼ 𝕌 [0, 2*π*] and *θ*_stim, mod2_ ∼ 𝕌 [0, 2*π*] and for modality specific Go-type tasks

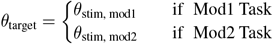

while for modality specific AntiGo-type tasks

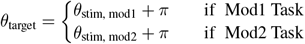

For ‘RT’ versions of the ‘Go’ tasks stimuli are only presented during the response epoch and the fixation cue is never extinguished. Thus the presence of the stimulus itself serves as the response cue and the model must respond as quickly as possible. Otherwise stimuli persist through the duration of the stimulus epoch.

#### ‘Decision-Making’ Tasks

The ‘Decision-Making’ family of tasks includes ‘DM’ (decision-making), ‘Anti DM’, ‘MultiDM’ (multi-sensory decision making), ‘AntiMultiDM,’ modality specific versions of each of these tasks and finally confidence based versions of ‘DM’ and ‘AntiDM.’ For all tasks in this group, two stimuli are presented simultaneously and persist throughout the duration of the stimulus epoch. They are drawn according to *θ*_stim1_ ∼ 𝕌 [0, 2*π*] and *θ*_stim2_ ∼ 𝕌 [(*θ*_stim1_ − 0.2*π, θ*_stim1_ − 0.6*π*) ∪ (*θ*_stim1_ + 0.2*π, θ*_stim1_ + 0.6*π*)]. A base strength applied to both stimuli is drawn such that *str*_*base*_ ∼ 𝕌 [1.0, 1.2]. A contrast is drawn from a discrete distribution such that *c* ∼ {−0.175, −0.15, −0.1, 0.1, 0.15, 0.175} so the stimulus strength associated with each direction in a trial are given by *str*_stim1_ = *str*_*base*_ + *c* and *str*_stim2_ = *str*_*base*_ − *c*.

For the ‘DM’ task, and for the the ‘Anti DM’ task,

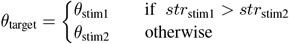

And for the the ‘Anti DM’ task,

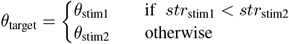

For these versions of the tasks the stimuli are presented in either modality 1 or modality 2 with equal probability. For the multi-sensory versions of each task, stimuli directions are drawn in the same manner and presented across both modalities so that *θ*_stim1, mod1_ = *θ*_stim1, mod2_ and *θ*_stim2, mod1_ = *θ*_stim2, mod2_. Base strengths are drawn independently for each modality. Contrasts for both modalities are drawn from a discrete distribution such that *c*_mod1_, *c*_mod2_ ∼ {0.2, 0.175, 0.15, −0.125, 0.125, −0.15, −0.175, −0.2} . If both |*c*_mod1_| −|*c*_mod2_| = 0 then contrasts are redrawn in order to avoid zero contrast trials during training. If both *c*_mod1_ and *c*_mod2_ have the same sign then contrasts are redrawn to ensure that the trial requires integrating over both modalities as opposed to simply performing a ‘DM’ task in a single modality. Criteria for target response are measured as the strength of a given direction summed over both modalities. So, for ‘MultiDM’

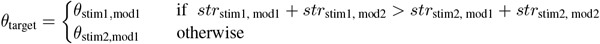

and for ‘AntiMultiDM’

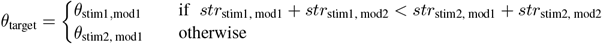

Stimuli for modality specific versions of each task are generated in the same way as multi-sensory versions of the task. Criteria for target response are the same as standard versions of ‘DM’ and ‘AntiDM’ tasks applied only to stimuli in the relevant modality.

In confidence based decision-making tasks (‘ConDM’ and ‘ConAntiDM’) the stimuli directions are drawn in the same way as above. Stimuli are shown in either modality 1 or modality 2 with equal probability. In each trial *str*_*base*_ = 1. The contrast and noise for each trial is based on the thresholded performance of a simpleNet model trained on all tasks except ‘ConDM’ and ‘ConAntiDM’. Once this model has been trained, we establish a threshold across levels of noise and contrasts for which the model can perform a ‘DM’ or an ‘AntiDM’ task at 95% correct. We then draw contrasts and noises for trials from above and below this threshold with equal probability during training. In trials where the noise and contrast levels fell below the 95% correct threshold the model must repress response, and otherwise perform the decision-making task (either ‘DM’ or ‘AntiDM’).

#### ‘Comparison’ Tasks

Our comparison task group includes ‘COMP1’, ‘COMP2’, ‘MultiCOMP1’, ‘MultiCOMP2’, ‘Anti’ versions of each of these tasks, as well as modality specific versions of ‘COMP1’ and ‘COMP2’ tasks. This group of tasks is designed to extend the basic decision-making framework into a setting with more complex control demands. These tasks utilize the delay structure in the stimulus epoch so that stim1 appears only during the stim1 epoch, followed by a delay, and finally stim2. This provides a temporal ordering on the stimuli. In ‘COMP1’ the model must respond to the first stimuli only if has greater strength than the second and otherwise repress a response i.e.

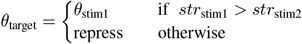

Likewise, in ‘COMP2’, the model must respond to the second direction if it presented with greater strength than the first otherwise repress response i.e.

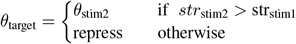

In ‘Anti’ versions of the task the ordering criteria is the same except for stimuli with least strength i.e. for ‘AntiCOMP1’

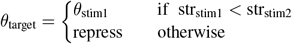

and for ‘AntiCOMP2’

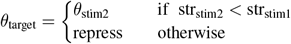

In multi-sensory settings the criteria for target direction is analogous to the multi-sensory decision-making tasks where strength is integrated across modalities. Likewise, for modality specific versions the criteria is only applied to stimuli in the relevant modality. Stimuli directions and strength for each of these tasks is drawn from the same distributions as the analogous task in the ‘Decision-Making’ family. However, during training we make sure to balance trials where responses are required and trials where models must repress response.

#### ‘Duration’ Tasks

The ‘Duration’ family of tasks includes ‘Dur1’, ‘Dur2’, ‘MultiDur1’, ‘MultiDur2’, ‘Anti’ versions of each of these tasks and modality specific versions of ‘Dur1’ and ‘Dur2’ tasks. These tasks require models to perform a time estimation task with the added demand or stimuli ordering determining relevance for response. Like in ‘Comparison’ tasks stim1 is presented followed by a delay and then stim2. For ‘Dur1’ trials

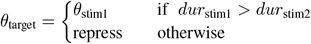

Likewise for ‘Dur2’

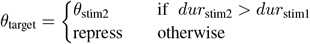

In ‘Anti’ versions of these tasks the correct response is in the direction of the stimulus with the shortest duration given the ordering criteria is met. Hence, for ‘AntiDur1’

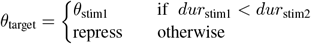

and for ‘AntiDur2’

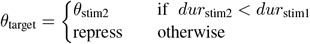

Across these tasks directions are drawn according to *θ*_stim1_ ∼ 𝕌 [0, 2*π*] and *θ*_stim2_ ∼ 𝕌 [(*θ*_stim1_ −0.2*π, θ*_stim1_ −0.6*π*) ∪ (*θ*_stim1_ + 0.2*π, θ*_stim1_ + 0.6*π*)]. Stimulus strengths are drawn according to *str*_stim1_, *str*_stim2_ ∼ 𝕌 [0.8, 1.2]. To set the duration of each stimulus we first draw *dur*_long_ ∼ {*i* |35 *< i* ≤50, *i*∈ ℕ } and *dur*_short_ ∼ {*i* |25 *< i* ≤ (*dur*_long_ −8), *i* ∈ ℕ } . During training, we determine which trials for a given task should and should not require a response in order to evenly balance repress and respond trials. We then assign *dur*_long_ and *dur*_short_ to either stim1 or stim2 so that the trial requires the appropriate response given the particular task type.

Again, criteria for correct response in the multi-sensory and modality specific versions of each tasks follow analogous tasks in the ‘Decision-Making’ and ‘Comparison’ groups where multi-sensory versions of the task require integrating total duration over each modality and modality specific tasks require only considering durations in the given task modality. For multi-sensory tasks we draw duration value *dur*_Long_ ∼ {*i* |75 *< i* ≤100, *i* ∈ℕ} and then split this value *dur*_long0_ = *dur*_Long_ ∗0.55 and *dur*_*long*1_ = *dur*_Long_ ∗0.45. We also draw a value *dur*_Short_ = *dur*_Long_ −∆*dur* where ∆*dur* ∼ {*i* |15 *< i* ≤25, *i* ∈ℕ . This value is then subdivided further into *dur*_short0_ = *dur*_long1_ + ∆*dur*_short_ where ∆*dur*_short_ ∼ {*i*| 19 *< i* ≤15, *i* ∈ℕ} and *dur*_short1_ = *dur*_Short_ −*dur*_short0_. Short and long durations can then be allocated to the ordered stimuli according to task type. Drawing durations in this manner ensures that like in ‘Decision-Making’ and ‘Comparison’ groups, correct answers truly require models to integrate durations over both modalities, rather than simply performing the task in a given modality to achieve correct responses.

#### ‘Matching’ Tasks

The ‘Matching’ family of tasks consists of ‘DMS’ (delay match to stimulus), ‘DNMS’ (delay non-match to stimulus), ‘DMC’ (delay match to category), ‘DMNC’ (delay non-match to category) tasks. For all tasks, stim1 is presented at the beginning of the stimulus epoch, followed by a delay, and the presentation of stim2. The stimulus strength is drawn according to *str*_stim1_, *str*_stim2_ ∼ 𝕌 [0.8, 1.2]. The input modality for any given trial is chosen at random with equal probability. In both ‘DMS’ and ‘DNMS’ tasks, trials are constructed as ‘matching stim’ trials or ‘mismatching stim’ trials with equal probability. In ‘matching stim’ trials *θ*_stim1_ ∼ 𝕌 [0, 2*π*] and *θ*_stim2_ = *θ*_stim1_. In ‘mismatch stim’ trials, *θ*_stim1_ ∼ 𝕌 [0, 2*π*] and *θ*_stim2_ ∼ 𝕌 [(*θ*_stim1_ −0.2*π, θ*_stim1_ −0.6*π*) ∪(*θ*_stim1_ + 0.2*π, θ*_stim1_ + 0.6*π*)]. For ‘DMS’, models must respond in the displayed direction if the stimuli match, otherwise repress response,

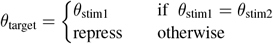

and for ‘DNMS’ models respond to the second direction if both directions are mismatched,

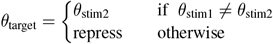

‘DMC’ and ‘DNMC’ tasks are organized in a similar manner. The stimulus input space is divided evenly into two categories such that cat1 = {*θ* : 0 *< θ* ≤*π}* and cat2 = {*θ* : *π < θ* ≤2*π}* . For ‘DMC’ and ‘DNMC’ tasks, trials are constructed as ‘matching cat.’ trials or ‘mismatching cat.’ trials with equal probability. In ‘matching cat.’ trials *θ*_stim1_ ∼ 𝕌 [0, 2*π*] and *θ*_stim2_ ∼ 𝕌 (cat_stim1_), where 𝕌 (cat_stim1_) is a uniform draw from the category of stim1. In ‘mismatch stim’ trials, *θ*_stim1_ ∼ 𝕌 [0, 2*π*] and *θ*_stim2_ ∼ 𝕌 (−cat_stim1_) where − cat_stim1_ is the opposite category as stim1. For ‘DMC’ the model must respond in the first direction if both stimuli are presented in the same category otherwise repress response,

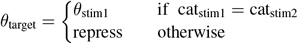

and for ‘DNMC’ the model should respond to the second direction if both stimuli are presented in opposite categories otherwise repress response,

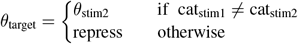

#### Target Output and Correct Criteria

The target output *y* ∈ℝ ^33 *×T*^ for a trial entails maintaining fixation in *y*_1_ = *y*_fix_ during the stimulus epoch, and then either responding in the correct direction or repressing activity in the remaining target response units *y*_2…33_ in the response epoch. Since the model should maintain fixation until response, target for fixation is set at *y*_fix_ = 0.85 during preparatory and stimulus epochs and *y*_fix_ = 0.05 in the response epoch. When a response is not required, as in the preparatory and stimulus epochs and with repressed activity in the response epoch, unit *i* takes on a target activity of *y*_*i*_ = 0.05. Alternatively, when there is a target direction for response,

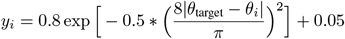

where *θ*_*i*_ is the preferred direction for unit *i*. Like in sensory stimuli, preferred directions for target units are evenly spaced values from [0, 2*π*] allocated to the 32 response units.

For a model response to count as correct, it must maintain fixation i.e. *ŷ*_fix_ *>* 0.5 during preparatory and stimulus epochs. When no response is required *ŷ*_*i*_ *<* 0.15. When a response is required, response activity is decoded using a population vector method and 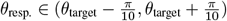. If the model fails to meet any of these criteria the trial response is incorrect.

### Model Training

Again following [31], model parameters are updated in a supervised fashion according to a masked Mean Squared Error Loss (mMSE) computed between the model motor response, *ŷ*_1…*T*_ = *ŷ* and the target *y*_1…*T*_ = *y* for each trial.

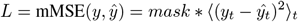

Here ∗denotes element-wise multiplication. Masks weigh the importance of different trial epochs. During preparatory and stimulus epochs mask weights are set to 1, during the first five time steps of the response epoch mask is set to 0, and during the remainder of the response epoch mask is set to 5. The mask value for the fixation is twice that of other values at all time steps.

For all models, we update Θ = {SensorimotorRNN, Linear_out_ } during training on our task set. For instructed models we additionally update Linear_embed_ in the process of normal training. We train models using standard PyTorch machinery and an Adam optimizer. An epoch consists of 2400 mini-batches, with each mini-batch consisting of 64 trials. Task type remains consistent within batches, but in the case of instructed models, the instruction used to inform each trial is drawn at random across the 64 trials in the batch. Tasks are randomly interleaved during training so that the networks can learn all tasks simultaneously. For all models we use the same initial learning rate as [31], *lr* = 0.001. We found that in the later phases of training model performance oscillated based on which latest task presented during training, so we decay the learning rate for each epoch by a factor of *γ* = 0.95 which allowed performance to converge smoothly. Following [31], models train until they reach a threshold performance of 95% across all tasks (and train for a minimum of 35 epochs). We found that training for gptNet tended to asymptote below performance threshold for multi-sensory versions of comparison tasks. This held true over a variety of training hyperparameters and learning rate scheduler regimes for training. Hence, we relax the performance threshold of gptNet to 85%. For each model type we train 5 models that start from 5 different random initializations. Where applicable results are averaged over these initializations.

#### Language Model Fine-Tuning

The pre-trained transformer architectures we use to embed task instructions are trained using a large corpus of natural language text according to wholly linguistic loss functions (except for model clipNet (S) which trains to jointly embed images and their text captions). In order to investigate the possibility that grounding linguistic representations more explicitly in the task set may improve performance, we allow the gradient from the motor loss experienced during sensorimotor training to fine-tune the weights in the final layers of the transformer language models. We do this for a relatively short period at the end of training. Specifically, during normal training we checkpoint a copy of our instructed models after training for 30 epochs. We then add the last 3 transformer layers to the set of trainable parameters, and reset the learning rates to *lr* = 1*e*^−4^ for Θ = {SensorimotorRNN, Linear_out_} and *lr*^lang^ = 3*e*^−4^ for Θ^lang^ = {Linear_embed_, transformer _− 3, −2, − 1_ } where transformer _−3, −2, − 1_ denotes the parameters of the last three layers of the relevant transformer architecture. In line with standard practice for language model fine-tuning, we used these reduced learning rates to avoid completely erasing pre-existing linguistic knowledge. Similarly for RNN parameters we found the above learning rate avoided catastrophic forgetting of sensorimotor knowledge while also allowing the RNN to adapt to updated language embeddings across all models. Autoregressive models were much more sensitive to this procedure, often collapsing at the beginning fine-tuning. Hence, for gptNetXL and gptNet we used *lr*^lang^ = 5*e*^−5^, which resulted in robust learning. Models train until they reach a threshold performance of 95% across training tasks or 85% correct for gptNet.

#### Hold-out Testing

During hold-out testing, we present models with 100 batches of one of the tasks that had been held out of training. For instructed model the only weights allowed to update during this phase are Θ = {SensorimotorRNN, Linear_out_, Linear_embed_} . All weights of simpleNet and structureNet are trainable in this context. In this holdout setting we found that in more difficult tasks for some of our more poorly performing models, the standard hyperparameters we used during training resulted in unstable learning curves for novel tasks. In order to stabilize performance and thereby create fair comparisons across models we used an increased batch size of 256. We then began with the standard learning rate of 0.001 and decreased this by increments of 0.0005 until all models showed robust learning curves. This resulted in a learning rate of 8*e* −4. All additional results shown in the Supplemental Information Section 4 follow this procedure.

### Cross Condition Generalization Performance Calculation

To calculate Cross Conditional Generalization Performance (CCGP) we train a linear decoder on a pair of tasks and then test that decoder on alternative pairs of task that have an analogous relationship. We group tasks into 8 dichotomies ‘Go’ vs. ‘Anti’, ‘Standard’ vs. ‘RT’, ‘Weakest’ vs. ‘Strongest’, ‘Longest’ vs. ‘Shortest’, ‘First Stim.’ vs. ‘Second Stim’, ‘Stim Match’ vs. ‘Category Match’, ‘Matching’ vs. ‘Non-Matching’, and ‘Mod1’ vs. ‘Mod2’. As an example, the ‘Go’ vs. ‘Anti’ dichotomy includes (‘Go’, ‘AntiGo’), (‘GoMod1’, ‘AntiGoMod1’), (‘GoMod2’, ‘AntiGoMod2’), (‘RTGo’, ‘AntiRTGo’), (‘RTGoMod1’, ‘AntiRTGoMod1’), (‘RTGoMod2’, ‘AntiRTGoMod2’) task pairs. For “RNN” task representations, we extracted activity at the time of stimulus onset for 250 example trials. For language representations, we input the instruction sets for relevant tasks to our language model and directly analyze activity in the “embedding” layer or take the sequence-averaged activity in each transformer layer. For non-linguistic models we simply analyze the space of rule vectors. We train a linear decoder to classify task activity according to one pair of tasks, and subsequently test the same classifier on the remaining set of pairs. Train and test conditions are determined by dichotomies identified across the task set (Supplementary Info. 12). To train and test decoders we use sklearn.svm.LinearSVC Python package. The CCGP score for a given task is the average decoding score achieved across all dichotomies where the task in question was either part of the train set or the test set. For model scores reported in the main text, we only calculate CCGP scores for models where the task in question has been held out of training. In this way we can capture the geometries that the models obtain in the zero shot setting. In Supplementary Info. 9 we report scores on tasks where models have been trained on all tasks, and for models where instructions have been switched for the holdout task.

For Fig. 3e. we calculate Pearson’s r correlation coefficient between performance on held out tasks and CCGP scores per task, as well as a p-value testing against the null hypothesis that these metrics are uncorrelated and normally distributed (using scipy.stats.pearsonr function). Full statistical tests for CCGP scores of both RNN and embedding layers from Fig. 3f. can be found in Supplementary Info. 9. Note that transformer language models use the same set of pre-trained weights amongst random initialization of Sensorimotor-RNNs so for Language Model Layers Fig. 3f plots the absolute scores of those language models.

### Conditional Clause/Deduction Task Analysis

We first split our task set into two groups (listed below): tasks that included conditional clauses and simple deductive reasoning components (30 tasks) and those where instructions include simple imperatives (20 tasks). We computed the difference in performance across the mean of generalization performance for each group across random initialization for each model (Fig 2f., solid dots). We compared these differences to a null distribution constructed by performing a set of 50 random shuffles of the task set into groups 30 and 20 tasks and computing differences in the same way, again using unequal variance t-tests. Because strucutreNet is a non-linguistic model we then compared performance of strucutreNet to our instructed models to disassociate the effects of performing tasks with a deductive reasoning component vs. processing instructions with more complicated conditional clause structure. Results of all statistical tests are reported in Supplementary Info. 6).

Simple Imperative Tasks: ‘Go’, ‘AntiGo’, ‘RTGo’, ‘AntiRTGo’, ‘GoMod1’, ‘GoMod2’, ‘AntiGoMod1’, ‘AntiGoMod2’, ‘RTGoMod1’, ‘AntiRTGoMod2’, ‘RTGoMod2’, ‘AntiRTGoMod2’, ‘DM’, ‘AntiDM’, ‘MultiDM’, ‘AntiMultiDM’, ‘DMMod1’, ‘DMMod2’, ‘AntiDMMod1’, ‘AntiDMMod2’

Conditional Clause/Deduction Tasks: ‘ConDM’, ‘ConAntiDM’, ‘Dur1’, ‘Dur2’, ‘MultiDur1’, ‘MultiDur2’, ‘AntiDur1’, ‘AntiDur2’, ‘AntiMultiDur1’, ‘AntiMultiDur2’, ‘Dur1Mod1’, ‘Dur1Mod2’, ‘Dur2Mod1’, ‘Dur2Mod2’, ‘COMP1’, ‘COMP2’, ‘MultiCOMP1’, ‘MultiCOMP2’, ‘AntiCOMP1’, ‘AntiCOMP2’, ‘AntiMultiCOMP1’, ‘AntiMultiCOMP2’, ‘COMP1Mod1’, ‘COMP1Mod2’, ‘COMP2Mod1’, ‘COMP2Mod2’, ‘DMS’, ‘DNMS’, ‘DMC’, ‘DMNC’

### Language Production Training

#### Self-Supervised Language Production Network Training

Our language production framework is inspired by classic sequence to sequence modeling using RNNs [78]. We treat each time step in the sequence as a token position in the target sentence that we wish to decode. Our ProductionRNN is a GRU with 256 hidden units using ReLU non-linearities. At each step in the sequence a set of decoder weights Linear_words_ attempts to decode the next token *w*_*τ*+1_ from the hidden state of the recurrent units. The hidden state of the ProductionRNN is initialized by concatenating the time average and maximum sensorimotor activity of a sbertNet (L) and passing that through weights Linear_sm_. Using an instructed model means that training the ProductionRNN can proceed in a completely self-supervised manner. The linguistic instruction used to drive the initializing sensorimotor activity is in turn used as the target set of tokens for the ProductionRNN outputs. The first input to the ProductionRNN is always a special start-of-sentence token (<SOS>) and the decoder runs until an end-of-sentence token (<EOS>) is decoded or until input reaches a length of 30 tokens. Suppose 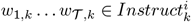is the sequence of tokens in instruction *k* where *k* is in the instruction set for task *i* and *X*^*i*^ is sensory input for a trial of task *i*. For brevity, we denote the process by which language models embed instructions as *Embed*() (see *Pre-trained Transformers* above).

The decoded token at the *τ*^*th*^ position, *ŵ*_*τ,k*_, is then given by

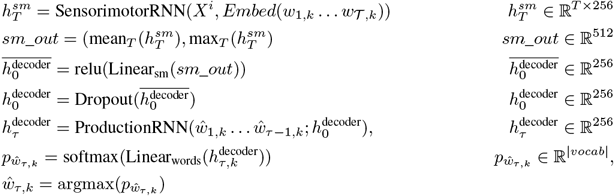

The model parameters Θ^production^ = Linear_sm_, Linear_words_, ProductionRNN are trained using cross-entropy loss between the 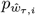and the instruction token *w*_*τ,k*_ provided to the SensorimotorRNN as input. Because there is no ground truth performance criteria here, we train for 80 epochs of 2400 batches with 64 trials per batch and with task type randomly interleaved. We found that using an initial learning rate of 0.001 sometimes caused models to diverge in early phases of training so we opted for a learning rate of *lr* = 1*e*^−4^ which led to stable early training. In order to alleviate similar oscillation problems detected in sensorimotor training we also decay the learning rate by *γ* = 0.99 per epoch. Additionally, the use of a dropout layer with a dropout rate of 0.05 improved performance. We also use a teacher forcing curriculum, where for some ratio of training batches we input the ground truth instruction token *w*_*τ,k*_ at each time step instead of the models decoded word *ŵ*_*τ,k*_. At each epoch, *teacher*_*forcing*_*ratio* 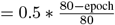.

#### Obtaining Embedding Layer Activity Using Motor Feedback

We wish to determine whether or not networks can communicate task information gained during training on motor feedback, in the absence of linguistic instructions. To accomplish this for a task, *i*, we seek to optimize a set of embedding activity vectors *E*^*i*^ ∈ ℝ^64^ such that when they are input to the model it will perform the task in question. Crucially, we freeze all model weights Θ = {SensorimotorRNN, Linear_out_, Linear_embedding_ } and only update *E*^*i*^ according to the standard supervised loss on the motor output that comes from inputting *E*^*i*^ as task identifying information. For notional clarity GRU dependence on the previous hidden state *h*_*t* − 1_ has been made implicit in the following equations.

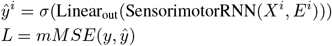

We optimize a set of 25 embedding vectors for each task, again using an Adam optimizer. Here the optimization space has many suboptimal local minimums corresponding to embeddings for related tasks which make learning prone to plateau. Hence, we use a high initial learning rate of *lr* = 0.05 which we decay by *γ* = 0.8 each epoch. This resulted in significantly more robust learning than lower learning rates. An epoch lasts for 800 batches with a batch length of 64 and we train for a minimum of 1 epoch or until we reach a threshold performance of 90% or 85% on ‘DMC’ and ‘DNMC’ tasks.

#### Producing Task Instructions

To produce task instructions we simply use the set *E*^*i*^ as task identifying information in the input of the SensorimotorRNN and use the ProductionRNN to output instructions based on the sensorimotor activity driven by *E*^*i*^. For each task we use the set of embedding vectors to produce 50 instructions per task. We repeat this process for each of the 5 initializations of SensorimotorRNN, resulting in 5 distinct language production networks, and 5 distinct sets of learned embedding vectors. Reported results for each task are averaged over these 5 networks. For the confusion matrix (Fig. 5d), we report the average percentage that decoded instructions are in the training instruction set for a given task or a novel instruction. Partner model performance (Fig. 5e) for each network initialization is computed by testing each of the 4 possible partner networks and averaging over these results.

## Code Availability

All code used to train models and analyze results can be found at https://github.com/ReidarRiveland/Instruct-RNN

## References

[1] Michael W Cole, Jeremy R Reynolds, Jonathan D Power, Grega Repovs, Alan Anticevic, and Todd S Braver. Multi-task connectivity reveals flexible hubs for adaptive task control. Nature neuroscience, 16(9):1348–1355, 2013.

[2] Takuya Ito, Guangyu Robert Yang, Patryk Laurent, Douglas H Schultz, and Michael W Cole. Constructing neural network models from brain data reveals representational transformations linked to adaptive behavior. Nature Communications, 13(1):1–16, 2022.

[3] Earl K. Miller and Jonathan D. Cohen. An integrative theory of prefrontal cortex function. Annual Review of Neuroscience, 24(1):167–202, 2001. PMID: 11283309.

[4] Michael W Cole, Todd S Braver, and Nachshon Meiran. The task novelty paradox: Flexible control of inflexible neural pathways during rapid instructed task learning. Neuroscience & Biobehavioral Reviews, 81:4–15, 2017.

[5] Hannes Ruge, Theo AJ Schäfer, Katharina Zwosta, Holger Mohr, and Uta Wolfensteller. Neural representation of newly instructed rule identities during early implementation trials. Elife, 8:e48293, 2019.

[6] Silvia Bernardi, Marcus K Benna, Mattia Rigotti, Jérôme Munuera, Stefano Fusi, and C Daniel Salzman. The geometry of abstraction in the hippocampus and prefrontal cortex. Cell, 183(4):954–967, 2020.

[7] Juri Minxha, Ralph Adolphs, Stefano Fusi, Adam N. Mamelak, and Ueli Rutishauser. Flexible recruitment of memory-based choice representations by the human medial frontal cortex. Science, 368(6498):eaba3313, 2020.

[8] Ramon Nogueira, Chris C. Rodgers, Randy M. Bruno, and Stefano Fusi. The geometry of cortical representations of touch in rodents. bioRxiv, 2021.

[9] Takuya Ito, Tim Klinger, Douglas H. Schultz, John D. Murray, Michael W. Cole, and Mattia Rigotti. Compositional generalization through abstract representations in human and artificial neural networks, 2022.

[10] Laura Driscoll, Krishna Shenoy, and David Sussillo. Flexible multitask computation in recurrent networks utilizes shared dynamical motifs. bioRxiv, 2022.

[11] Ana F Palenciano, Carlos González-García, Juan E Arco, Luiz Pessoa, and María Ruz. Representational organization of novel task sets during proactive encoding. Journal of Neuroscience, 39(42):8386–8397, 2019.

[12] Alberto Sobrado, Ana F. Palenciano, Carlos González-García, and María Ruz. The effect of task demands on the neural patterns generated by novel instruction encoding. Cortex, 149:59–72, 2022.

[13] Carlos González-García, Juan E Arco, Ana F Palenciano, Javier Ramírez, and María Ruz. Encoding, preparation and implementation of novel complex verbal instructions. Neuroimage, 148:264–273, 2017.

[14] Paul S Muhle-Karbe, John Duncan, Wouter De Baene, Daniel J Mitchell, and Marcel Brass. Neural coding for instruction-based task sets in human frontoparietal and visual cortex. Cerebral Cortex, 27(3):1891–1905, 2017.

[15] Tom B. Brown, Benjamin Mann, Nick Ryder, Melanie Subbiah, Jared Kaplan, Prafulla Dhariwal, Arvind Neelakantan, Pranav Shyam, Girish Sastry, Amanda Askell, Sandhini Agarwal, Ariel Herbert-Voss, Gretchen Krueger, Tom Henighan, Rewon Child, Aditya Ramesh, Daniel M. Ziegler, Jeffrey Wu, Clemens Winter, Christopher Hesse, Mark Chen, Eric Sigler, Mateusz Litwin, Scott Gray, Benjamin Chess, Jack Clark, Christopher Berner, Sam McCandlish, Alec Radford, Ilya Sutskever, and Dario Amodei. Language models are few-shot learners. CoRR, abs/2005.14165, 2020.

[16] Long Ouyang, Jeff Wu, Xu Jiang, Diogo Almeida, Carroll L. Wainwright, Pamela Mishkin, Chong Zhang, Sandhini Agarwal, Katarina Slama, Alex Ray, John Schulman, Jacob Hilton, Fraser Kelton, Luke Miller, Maddie Simens, Amanda Askell, Peter Welinder, Paul Christiano, Jan Leike, and Ryan Lowe. Training language models to follow instructions with human feedback, 2022.

[17] Aditya Ramesh, Mikhail Pavlov, Gabriel Goh, Scott Gray, Chelsea Voss, Alec Radford, Mark Chen, and Ilya Sutskever. Zero-shot text-to-image generation. CoRR, abs/2102.12092, 2021.

[18] Alec Radford, Jeffrey Wu, Rewon Child, David Luan, Dario Amodei, Ilya Sutskever, et al. Language models are unsupervised multitask learners. OpenAI blog, 1(8):9, 2019.

[19] Martin Schrimpf, Idan Asher Blank, Greta Tuckute, Carina Kauf, Eghbal A. Hosseini, Nancy Kanwisher, Joshua B. Tenenbaum, and Evelina Fedorenko. The neural architecture of language: Integrative modeling converges on predictive processing. Proceedings of the National Academy of Sciences, 118(45), 2021.

[20] Ariel Goldstein, Zaid Zada, Eliav Buchnik, Mariano Schain, Amy Price, Bobbi Aubrey, Samuel A Nastase, Amir Feder, Dotan Emanuel, Alon Cohen, et al. Shared computational principles for language processing in humans and deep language models. Nature neuroscience, 25(3):369–380, 2022.

[21] Aakanksha Chowdhery, Sharan Narang, Jacob Devlin, Maarten Bosma, Gaurav Mishra, Adam Roberts, Paul Barham, Hyung Won Chung, Charles Sutton, Sebastian Gehrmann, Parker Schuh, Kensen Shi, Sasha Tsvyashchenko, Joshua Maynez, Abhishek Rao, Parker Barnes, Yi Tay, Noam Shazeer, Vinodkumar Prabhakaran, Emily Reif, Nan Du, Ben Hutchinson, Reiner Pope, James Bradbury, Jacob Austin, Michael Isard, Guy Gur-Ari, Pengcheng Yin, Toju Duke, Anselm Levskaya, Sanjay Ghemawat, Sunipa Dev, Henryk Michalewski, Xavier Garcia, Vedant Misra, Kevin Robinson, Liam Fedus, Denny Zhou, Daphne Ippolito, David Luan, Hyeontaek Lim, Barret Zoph, Alexander Spiridonov, Ryan Sepassi, David Dohan, Shivani Agrawal, Mark Omernick, Andrew M. Dai, Thanumalayan Sankaranarayana Pillai, Marie Pellat, Aitor Lewkowycz, Erica Moreira, Rewon Child, Oleksandr Polozov, Katherine Lee, Zongwei Zhou, Xuezhi Wang, Brennan Saeta, Mark Diaz, Orhan Firat, Michele Catasta, Jason Wei, Kathy Meier-Hellstern, Douglas Eck, Jeff Dean, Slav Petrov, and Noah Fiedel. Palm: Scaling language modeling with pathways, 2022.

[22] Romal Thoppilan, Daniel De Freitas, Jamie Hall, Noam Shazeer, Apoorv Kulshreshtha, Heng-Tze Cheng, Alicia Jin, Taylor Bos, Leslie Baker, Yu Du, YaGuang Li, Hongrae Lee, Huaixiu Steven Zheng, Amin Ghafouri, Marcelo Menegali, Yanping Huang, Maxim Krikun, Dmitry Lepikhin, James Qin, Dehao Chen, Yuanzhong Xu, Zhifeng Chen, Adam Roberts, Maarten Bosma, Vincent Zhao, Yanqi Zhou, Chung-Ching Chang, Igor Krivokon, Will Rusch, Marc Pickett, Pranesh Srinivasan, Laichee Man, Kathleen Meier-Hellstern, Meredith Ringel Morris, Tulsee Doshi, Renelito Delos Santos, Toju Duke, Johnny Soraker, Ben Zevenbergen, Vinodkumar Prabhakaran, Mark Diaz, Ben Hutchinson, Kristen Olson, Alejandra Molina, Erin Hoffman-John, Josh Lee, Lora Aroyo, Ravi Rajakumar, Alena Butryna, Matthew Lamm, Viktoriya Kuzmina, Joe Fenton, Aaron Cohen, Rachel Bernstein, Ray Kurzweil, Blaise Aguera-Arcas, Claire Cui, Marian Croak, Ed Chi, and Quoc Le. Lamda: Language models for dialog applications, 2022.

[23] Robin Rombach, Andreas Blattmann, Dominik Lorenz, Patrick Esser, and Björn Ommer. High-resolution image synthesis with latent diffusion models, 2021.

[24] Anthony Brohan, Noah Brown, Justice Carbajal, Yevgen Chebotar, Xi Chen, Krzysztof Choromanski, Tianli Ding, Danny Driess, Avinava Dubey, Chelsea Finn, Pete Florence, Chuyuan Fu, Montse Gonzalez Arenas, Keerthana Gopalakrishnan, Kehang Han, Karol Hausman, Alex Herzog, Jasmine Hsu, Brian Ichter, Alex Irpan, Nikhil Joshi, Ryan Julian, Dmitry Kalashnikov, Yuheng Kuang, Isabel Leal, Lisa Lee, Tsang-Wei Edward Lee, Sergey Levine, Yao Lu, Henryk Michalewski, Igor Mordatch, Karl Pertsch, Kanishka Rao, Krista Reymann, Michael Ryoo, Grecia Salazar, Pannag Sanketi, Pierre Sermanet, Jaspiar Singh, Anikait Singh, Radu Soricut, Huong Tran, Vincent Vanhoucke, Quan Vuong, Ayzaan Wahid, Stefan Welker, Paul Wohlhart, Jialin Wu, Fei Xia, Ted Xiao, Peng Xu, Sichun Xu, Tianhe Yu, and Brianna Zitkovich. Rt-2: Vision-language-action models transfer web knowledge to robotic control. In arXiv preprint arXiv:2307.15818, 2023.

[25] Josh Abramson, Arun Ahuja, Iain Barr, Arthur Brussee, Federico Carnevale, Mary Cassin, Rachita Chhaparia, Stephen Clark, Bogdan Damoc, Andrew Dudzik, Petko Georgiev, Aurelia Guy, Tim Harley, Felix Hill, Alden Hung, Zachary Kenton, Jessica Landon, Timothy Lillicrap, Kory Mathewson, Soňa Mokrá, Alistair Muldal, Adam Santoro, Nikolay Savinov, Vikrant Varma, Greg Wayne, Duncan Williams, Nathaniel Wong, Chen Yan, and Rui Zhu. Imitating interactive intelligence, 2021.

[26] DeepMind Interactive Agents Team, Josh Abramson, Arun Ahuja, Arthur Brussee, Federico Carnevale, Mary Cassin, Felix Fischer, Petko Georgiev, Alex Goldin, Mansi Gupta, Tim Harley, Felix Hill, Peter C Humphreys, Alden Hung, Jessica Landon, Timothy Lillicrap, Hamza Merzic, Alistair Muldal, Adam Santoro, Guy Scully, Tamara von Glehn, Greg Wayne, Nathaniel Wong, Chen Yan, and Rui Zhu. Creating multimodal interactive agents with imitation and self-supervised learning, 2022.

[27] Joshua I Gold and Michael N Shadlen. The neural basis of decision making. Annu. Rev. Neurosci., 30:535–574, 2007.

[28] Maxime Chevée, Eric A. Finkel, Su-Jeong Kim, Daniel H. O’Connor, and Solange P. Brown. Neural activity in the mouse claustrum in a cross-modal sensory selection task. Neuron, 110(3):486–501.e7, 2022.

[29] Sofia Soares, Bassam V Atallah, and Joseph J Paton. Midbrain dopamine neurons control judgment of time. Science, 354(6317):1273–1277, 2016.

[30] Christopher Summerfield and Floris P De Lange. Expectation in perceptual decision making: neural and computational mechanisms. Nature Reviews Neuroscience, 15(11):745–756, 2014.

[31] Guangyu Robert Yang, Madhura R Joglekar, H Francis Song, William T Newsome, and Xiao-Jing Wang. Task representations in neural networks trained to perform many cognitive tasks. Nature neuroscience, 22(2):297–306, 2019.

[32] Alexis Dubreuil, Adrian Valente, Manuel Beiran, Francesca Mastrogiuseppe, and Srdjan Ostojic. The role of population structure in computations through neural dynamics. Nature Neuroscience, pages 1–12, 2022.

[33] Douglas P Munoz and Stefan Everling. Look away: the anti-saccade task and the voluntary control of eye movement. Nature Reviews Neuroscience, 5(3):218–228, 2004.

[34] Earl K Miller, Cynthia A Erickson, and Robert Desimone. Neural mechanisms of visual working memory in prefrontal cortex of the macaque. Journal of neuroscience, 16(16):5154–5167, 1996.

[35] David J Freedman and John A Assad. Neuronal mechanisms of visual categorization: an abstract view on decision making. Annual review of neuroscience, 39:129–147, 2016.

[36] Nick Yeung and Christopher Summerfield. Metacognition in human decision-making: confidence and error monitoring. Philosophical Transactions of the Royal Society B: Biological Sciences, 367(1594):1310–1321, 2012.

[37] Ashish Vaswani, Noam Shazeer, Niki Parmar, Jakob Uszkoreit, Llion Jones, Aidan N. Gomez, Lukasz Kaiser, and Illia Polosukhin. Attention is all you need. CoRR, abs/1706.03762, 2017.

[38] Alec Radford, Jeffrey Wu, Jack Clark, Daniela Amodei, Drundage Miles, David Luan, Dario Amodei, Ilya Sutskever, et al. Better language models and their implications, 2019.

[39] Jacob Devlin, Ming-Wei Chang, Kenton Lee, and Kristina Toutanova. BERT: pre-training of deep bidirectional transformers for language understanding. CoRR, abs/1810.04805, 2018.

[40] Nils Reimers and Iryna Gurevych. Sentence-bert: Sentence embeddings using siamese bert-networks, 2019.

[41] Samuel R. Bowman, Gabor Angeli, Christopher Potts, and Christopher D. Manning. A large annotated corpus for learning natural language inference. CoRR, abs/1508.05326, 2015.

[42] Alec Radford, Jong Wook Kim, Chris Hallacy, Aditya Ramesh, Gabriel Goh, Sandhini Agarwal, Girish Sastry, Amanda Askell, Pamela Mishkin, Jack Clark, Gretchen Krueger, and Ilya Sutskever. Learning transferable visual models from natural language supervision. CoRR, abs/2103.00020, 2021.

[43] Lawrence W Barsalou. Grounded cognition. Annu. Rev. Psychol., 59:617–645, 2008.

[44] Vinod Goel, Brian Gold, Shitij Kapur, and Sylvain Houle. Neuroanatomical correlates of human reasoning. Journal of cognitive neuroscience, 10(3):293–302, 1998.

[45] Vinod Goel, Christian Buchel, Chris Frith, and Raymond J Dolan. Dissociation of mechanisms underlying syllogistic reasoning. Neuroimage, 12(5):504–514, 2000.

[46] Carlo Reverberi, Paolo Cherubini, Attilio Rapisarda, Elisa Rigamonti, Carlo Caltagirone, Richard SJ Frackowiak, Emiliano Macaluso, and Eraldo Paulesu. Neural basis of generation of conclusions in elementary deduction. Neuroimage, 38(4):752–762, 2007.

[47] Ira A Noveck, Vinod Goel, and Kathleen W Smith. The neural basis of conditional reasoning with arbitrary content. Cortex, 40(4-5):613–622, 2004.

[48] Martin M Monti, Daniel N Osherson, Michael J Martinez, and Lawrence M Parsons. Functional neuroanatomy of deductive inference: a language-independent distributed network. Neuroimage, 37(3):1005–1016, 2007.

[49] Martin M Monti, Lawrence M Parsons, and Daniel N Osherson. The boundaries of language and thought in deductive inference. Proceedings of the National Academy of Sciences, 106(30):12554–12559, 2009.

[50] John P Coetzee and Martin M Monti. At the core of reasoning: Dissociating deductive and non-deductive load. Human Brain Mapping, 39(4):1850–1861, 2018.

[51] Martin M Monti and Daniel N Osherson. Logic, language and the brain. Brain research, 1428:33–42, 2012.

[52] Jérôme Prado. The relationship between deductive reasoning and the syntax of language in broca’s area: A review of the neuroimaging literature. L’année Psychologique, 118(3):289–315, 2018.

[53] Michael N Shadlen and William T Newsome. Neural basis of a perceptual decision in the parietal cortex (area lip) of the rhesus monkey. Journal of neurophysiology, 86(4):1916–1936, 2001.

[54] Alexander C Huk and Michael N Shadlen. Neural activity in macaque parietal cortex reflects temporal integration of visual motion signals during perceptual decision making. Journal of Neuroscience, 25(45):10420–10436, 2005.

[55] Matthew F Panichello and Timothy J Buschman. Shared mechanisms underlie the control of working memory and attention. Nature, 592(7855):601–605, 2021.

[56] Edward H Nieh, Manuel Schottdorf, Nicolas W Freeman, Ryan J Low, Sam Lewallen, Sue Ann Koay, Lucas Pinto, Jeffrey L Gauthier, Carlos D Brody, and David W Tank. Geometry of abstract learned knowledge in the hippocampus. Nature, 595(7865):80–84, 2021.

[57] Evelina Fedorenko and Idan A Blank. Broca’s area is not a natural kind. Trends in cognitive sciences, 24(4):270–284, 2020.

[58] Evelina Fedorenko, John Duncan, and Nancy Kanwisher. Language-selective and domain-general regions lie side by side within broca’s area. Current Biology, 22(21):2059–2062, 2012.

[59] Zhiyao Gao, Li Zheng, Rocco Chiou, André Gouws, Katya Krieger-Redwood, Xiuyi Wang, Dominika Varga, Matthew A Lambon Ralph, Jonathan Smallwood, and Elizabeth Jefferies. Distinct and common neural coding of semantic and non-semantic control demands. NeuroImage, 236:118230, 2021.

[60] John Duncan. The multiple-demand (md) system of the primate brain: mental programs for intelligent behaviour. Trends in cognitive sciences, 14(4):172–179, 2010.

[61] Giovanni Buccino, Ivan Colagè, Nicola Gobbi, and Giorgio Bonaccorso. Grounding meaning in experience: A broad perspective on embodied language. Neuroscience Biobehavioral Reviews, 69:69–78, 2016.

[62] Farshad Alizadeh Mansouri, David J Freedman, and Mark J Buckley. Emergence of abstract rules in the primate brain. Nature Reviews Neuroscience, 21(11):595–610, 2020.

[63] David Chen and Raymond Mooney. Learning to interpret natural language navigation instructions from observations. In Proceedings of the AAAI Conference on Artificial Intelligence, volume 25, pages 859–865, 2011.

[64] Junhyuk Oh, Satinder P. Singh, Honglak Lee, and Pushmeet Kohli. Zero-shot task generalization with multi-task deep reinforcement learning. CoRR, abs/1706.05064, 2017.

[65] Mohit Shridhar, Jesse Thomason, Daniel Gordon, Yonatan Bisk, Winson Han, Roozbeh Mottaghi, Luke Zettle-moyer, and Dieter Fox. ALFRED: A benchmark for interpreting grounded instructions for everyday tasks. CoRR, abs/1912.01734, 2019.

[66] Devendra Singh Chaplot, Kanthashree Mysore Sathyendra, Rama Kumar Pasumarthi, Dheeraj Rajagopal, and Ruslan Salakhutdinov. Gated-attention architectures for task-oriented language grounding. CoRR, abs/1706.07230, 2017.

[67] Howard Chen, Alane Suhr, Dipendra Kumar Misra, Noah Snavely, and Yoav Artzi. Touchdown: Natural language navigation and spatial reasoning in visual street environments. CoRR, abs/1811.12354, 2018.

[68] Victor Zhong, Tim Rocktäschel, and Edward Grefenstette. RTFM: generalising to novel environment dynamics via reading. CoRR, abs/1910.08210, 2019.

[69] Pratyusha Sharma, Antonio Torralba, and Jacob Andreas. Skill induction and planning with latent language. CoRR, abs/2110.01517, 2021.

[70] Yiding Jiang, Shixiang Gu, Kevin Murphy, and Chelsea Finn. Language as an abstraction for hierarchical deep reinforcement learning. CoRR, abs/1906.07343, 2019.

[71] Jacob Andreas, Dan Klein, and Sergey Levine. Modular multitask reinforcement learning with policy sketches. CoRR, abs/1611.01796, 2016.

[72] Dzmitry Bahdanau, Felix Hill, Jan Leike, Edward Hughes, Arian Hosseini, Pushmeet Kohli, and Edward Grefenstette. Learning to understand goal specifications by modelling reward. arXiv preprint arXiv:1806.01946, 2018.

[73] Prasoon Goyal, Scott Niekum, and Raymond J. Mooney. Using natural language for reward shaping in reinforcement learning. CoRR, abs/1903.02020, 2019.

[74] Wenlong Huang, Pieter Abbeel, Deepak Pathak, and Igor Mordatch. Language models as zero-shot planners: Extracting actionable knowledge for embodied agents, 2022.

[75] Shuang Li, Xavier Puig, Yilun Du, Clinton Wang, Ekin Akyurek, Antonio Torralba, Jacob Andreas, and Igor Mordatch. Pre-trained language models for interactive decision-making, 2022.

## Methods References

[76] Jane Bromley, James W Bentz, Léon Bottou, Isabelle Guyon, Yann LeCun, Cliff Moore, Eduard Säckinger, and Roopak Shah. Signature verification using a “siamese” time delay neural network. International Journal of Pattern Recognition and Artificial Intelligence, 7(04):669–688, 1993.

[77] Junyoung Chung, Caglar Gulcehre, KyungHyun Cho, and Yoshua Bengio. Empirical evaluation of gated recurrent neural networks on sequence modeling. arXiv preprint arXiv:1412.3555, 2014.

[78] Ilya Sutskever, Oriol Vinyals, and Quoc V. Le. Sequence to sequence learning with neural networks. CoRR, abs/1409.3215, 2014.

[79] Thomas Wolf, Lysandre Debut, Victor Sanh, Julien Chaumond, Clement Delangue, Anthony Moi, Pierric Cistac, Tim Rault, Rémi Louf, Morgan Funtowicz, and Jamie Brew. Huggingface’s transformers: State-of-the-art natural language processing. CoRR, abs/1910.03771, 2019.

