## Supplementary Information for "Generalization in Sensorimotor Networks Configured with Natural Language Instructions"

### 1 Learning Curves Across All Tasks

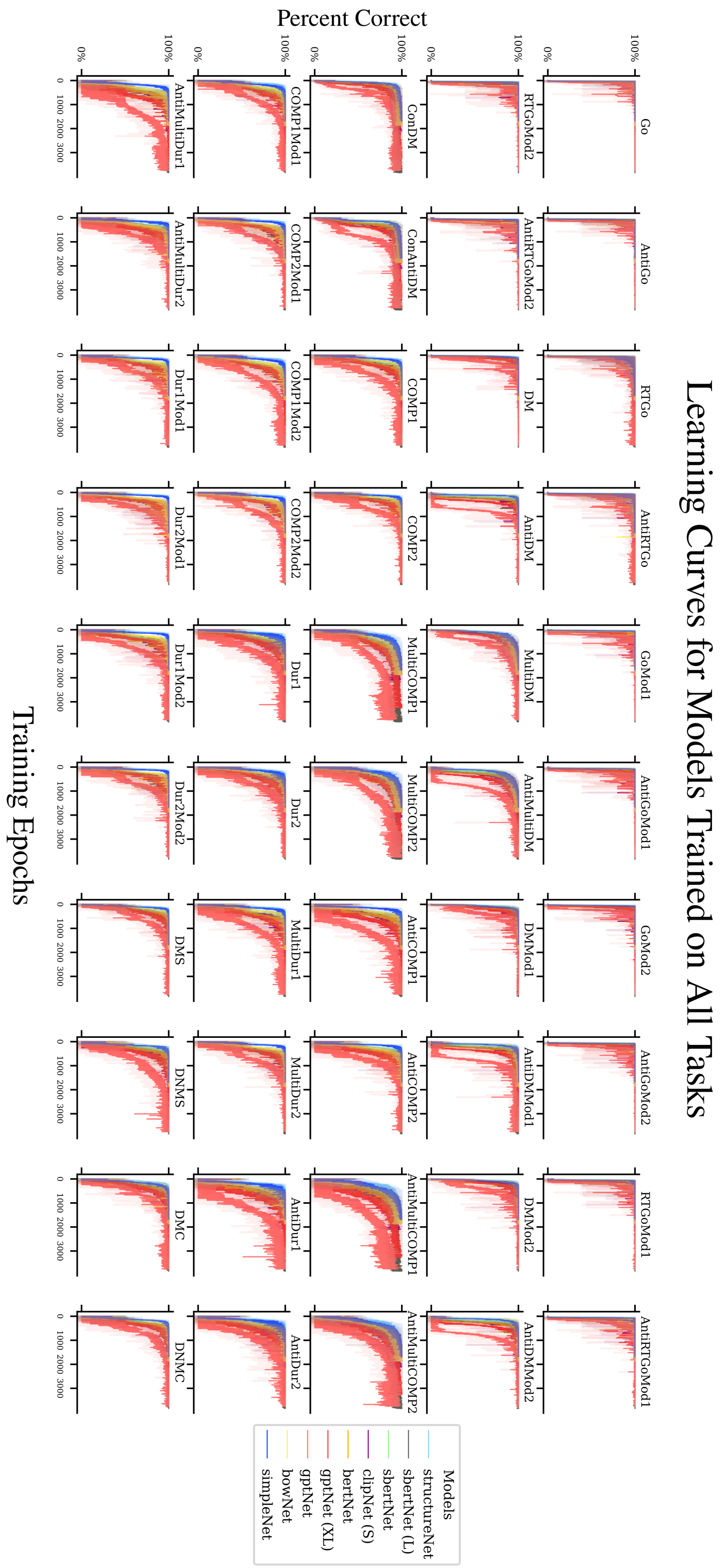


**Supplementary Info. Fig. 1**

#### 2 Validation Instructions

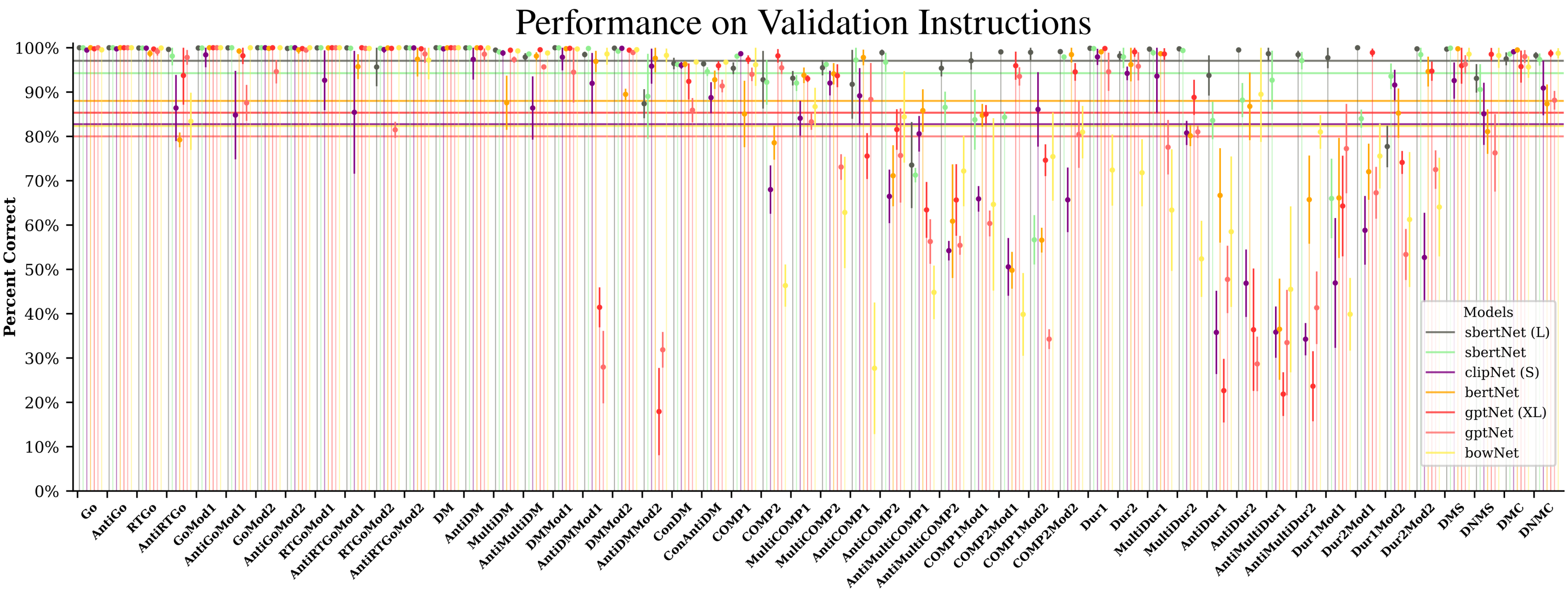


**Supplementary Info. Fig. 2**

Bars indicate plus or minus one standard deviation amongst 5 different random initializations of model parameters

#### 3 Zero-Shot Performance Across Tasks

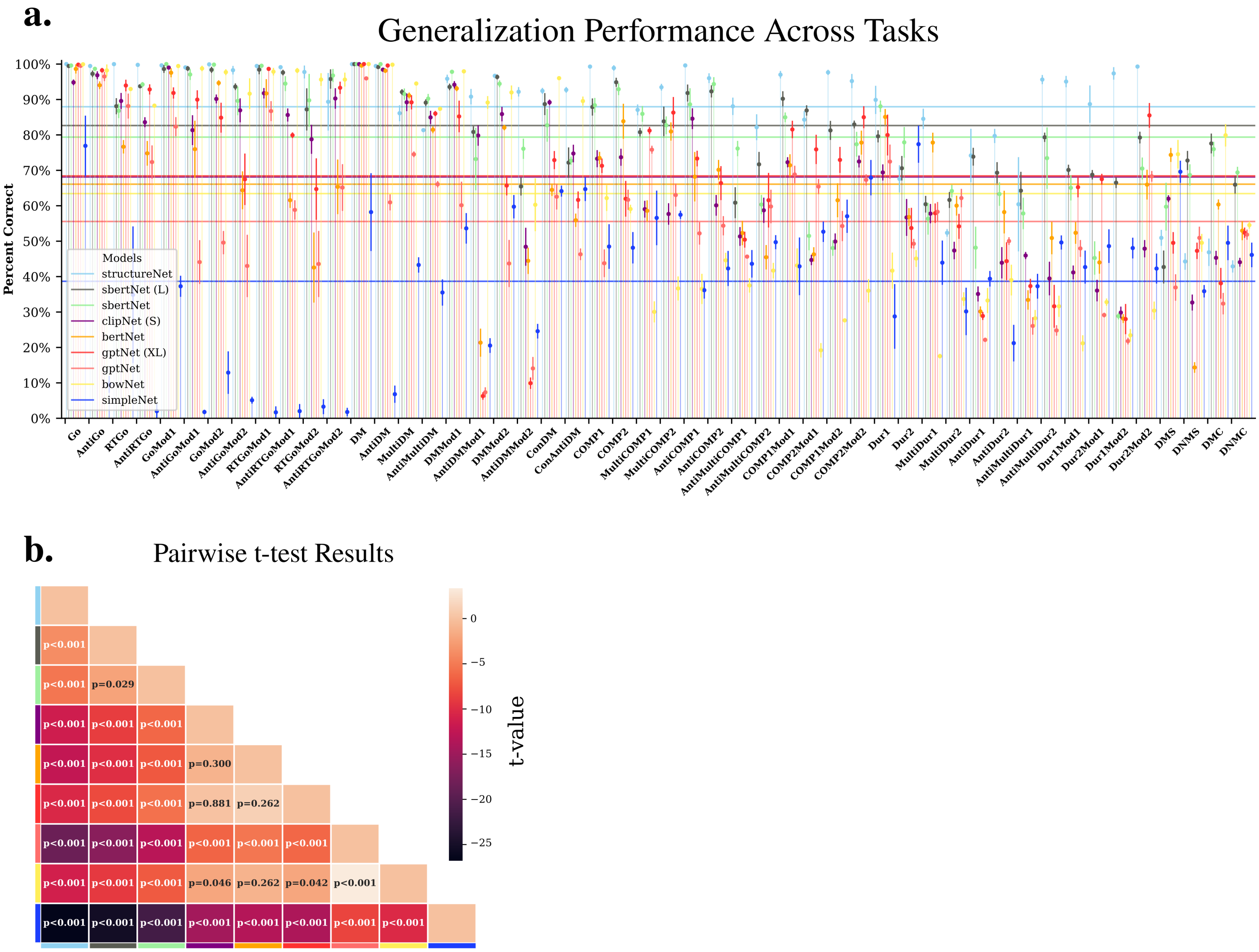


**Supplementary Info. Fig. 3**

**a** Performance across all tasks **b** unequal variance t-test comparison for generalization performance of results in Fig. 2a,b.

### 4 Additional Holdout Test

#### 4.1 Swap Holdouts

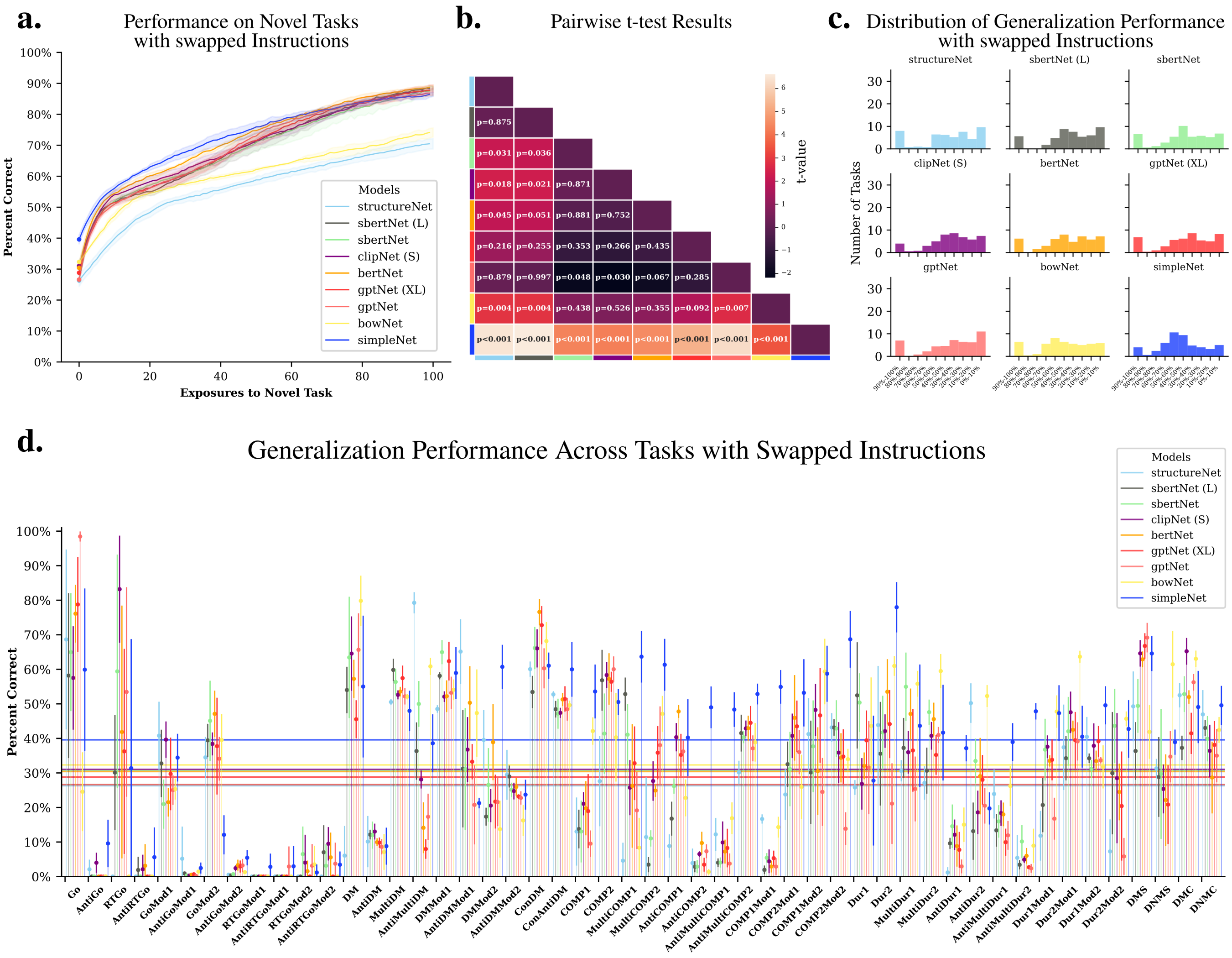


**Supplementary Info. Fig. 4.1**

The following groups of tasks are held out from models in the main results. Tasks were chosen from diverse groups to ensure that when we swapped instructions (results above), it didn't inadvertently lead to a correct result for the swapped task (e.g. swapping 'DM' and 'AntiDM' instructions would lead to equally valid results given both tasks share the same underlying input structure). We swap task instructions in the following manner:

Task Swaps:

*swap0*: AntiDMMod2 $\rightarrow$ RTGo; RTGo $\rightarrow$ DM; DM $\rightarrow$ MultiCOMP2 ; MultiCOMP2 $\rightarrow$ AntiMultiDur1; AntiMultiDur1 $\rightarrow$ AntiDMMod2;

*swap1*: COMP1Mod1 $\rightarrow$ AntiGoMod2; AntiGoMod2 $\rightarrow$ DMS; DMS $\rightarrow$ AntiDur1; AntiDur1 $\rightarrow$ RTGoMod2; RTGoMod2 $\rightarrow$ COMP1Mod1;

*swap2*: RTGoMod1 $\rightarrow$ AntiCOMP2; AntiCOMP2 $\rightarrow$ AntiRTGo; AntiRTGo $\rightarrow$ Dur2; Dur2 $\rightarrow$ MultiCOMP1; MultiCOMP1 $\rightarrow$ RTGoMod1;

*swap3*: GoMod2 $\rightarrow$ AntiMultiCOMP2; AntiMultiCOMP2 $\rightarrow$ DMMod2; DMMod2 $\rightarrow$ AntiRTGoMod1; AntiRTGo-Mod1 $\rightarrow$ AntiDur2; AntiDur2 $\rightarrow$ GoMod2;

*swap4*: MultiDM → COMP2Mod2; COMP2Mod2 → AntiMultiCOMP1; AntiMultiCOMP1 → AntiGoMod1; AntiGoMod1 → Dur1Mod1; Dur1Mod1 → MultiDM;

*swap5*: AntiDM → AntiRTGoMod2; AntiRTGoMod2 → Dur2Mod2; Dur2Mod2 → AntiCOMP1; AntiCOMP1 → DNMS; DNMS → AntiDM;

*swap6*: MultiDur1 → GoMod1; GoMod1 → COMP2; COMP2 → DMC; DMC → Dur2Mod1; Dur2Mod1 → MultiDur1;

*swap7*: COMP2Mod2 → AntiMultiDM; AntiMultiDM → DNMC; DNMC → DMMod1; DMMod1 → Dur1Mod2; Dur1Mod2 → COMP2Mod1;

*swap8*: ConAntiDM → COMP1; COMP1 → MultiDur2; MultiDur2 → COMP1Mod2; COMP1Mod2 → Go; Go → ConAntiDM;

*swap9*: AntiGo → Dur1; Dur1 → ConDM; ConDM → AntiDMMod1; AntiDMMod1 → AntiMultiDur2; AntiMultiDur2 → AntiGo

##### 4.2 Family Holdouts

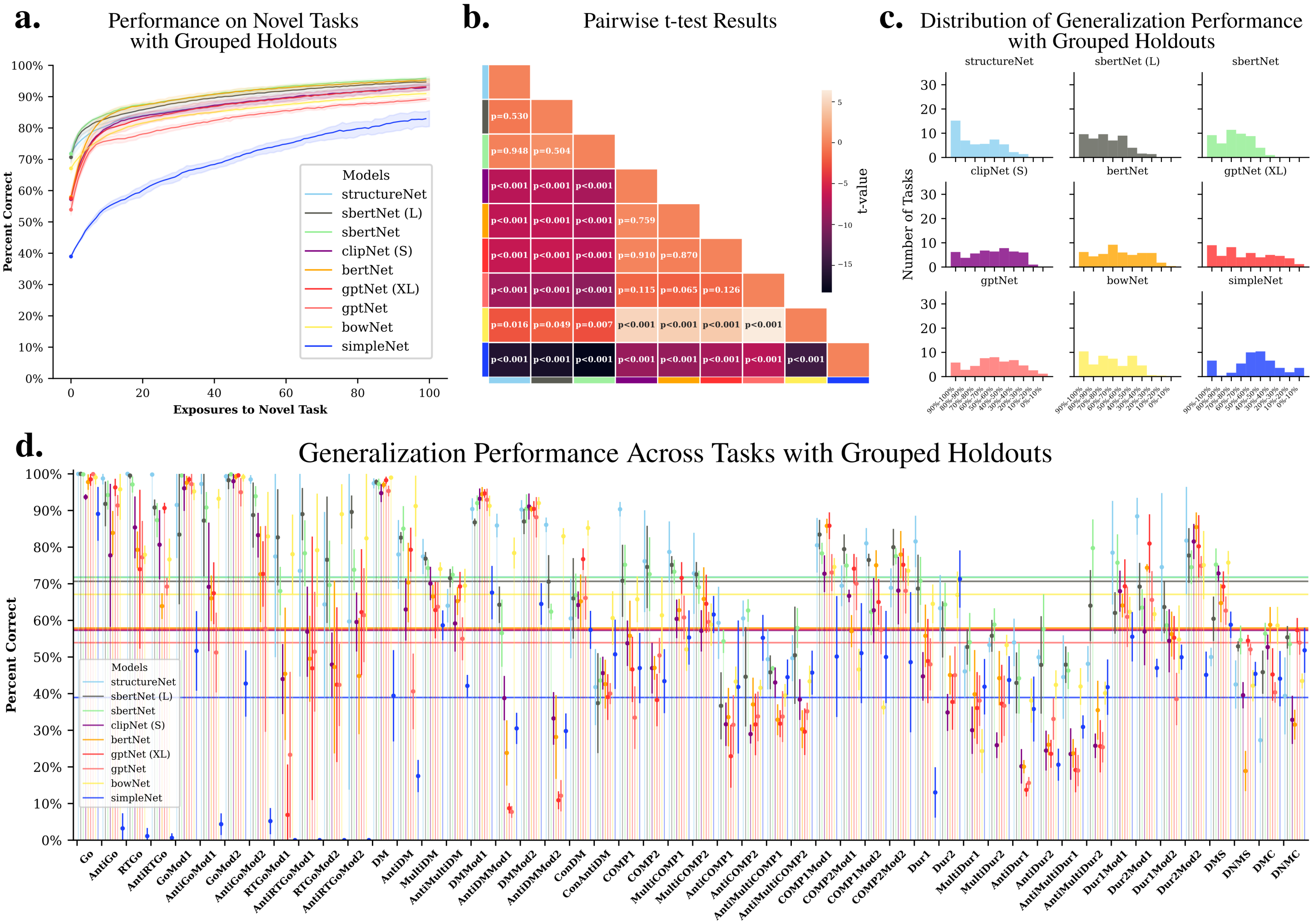


**Supplementary Info. Fig. 4.2**

We also test models in the more difficult setting where we hold out tasks from an family of tasks, forcing the model to infer an entire set of interrelated sensorimotor mappings from comparatively unfamiliar instructions.

*Group 0*: ‘Go’, ‘AntiGo’, ‘RTGo’, ‘AntiRTGo’

*Group 1*: ‘GoMod1’, ‘GoMod2’, ‘AntiGoMod1’, ‘AntiGoMod2’

*Group 2*: ‘RTGoMod1’, ‘RTGoMod2’, ‘AntiRTGoMod1’,‘’AntiRTGoMod2’

*Group 3*: ‘DM’, ‘AntiDM’, ‘MultiDM’, ‘AntiMultiDM’, ‘ConDM’, ‘ConAntiDM’

*Group 4*: ‘DMMod1’, ‘AntiDMMod1’, ‘DMMod2’, ‘AntiDMMod2’

*Group 5*: ‘COMP1’, ‘COMP2’, ‘MultiCOMP1’, ‘MultiCOMP2’

*Group 6*: ‘AntiCOMP1’, ‘AntiCOMP2’, ‘AntiMultiCOMP1’, ‘AntiMultiCOMP2’

*Group 7*: ‘COMP1Mod1’, ‘COMP2Mod1’, ‘COMP1Mod2’, ‘COMP2Mod2’

*Group 8*: ‘Dur1’, ‘Dur2’, ‘MultiDur1’, ‘MultiDur2’

*Group 9*: ‘AntiDur1’, ‘AntiDur2’, ‘AntiMultiDur1’, ‘AntiMultiDur2’

*Group 10*: ‘Dur1Mod1’, ‘Dur2Mod1’, ‘Dur1Mod2’, ‘Dur2Mod2’

*Group 11*: ‘DMS’, ‘DNMS’, ‘DMC’, ‘DNMC’

##### 4.3 Language Model Fine-Tuning

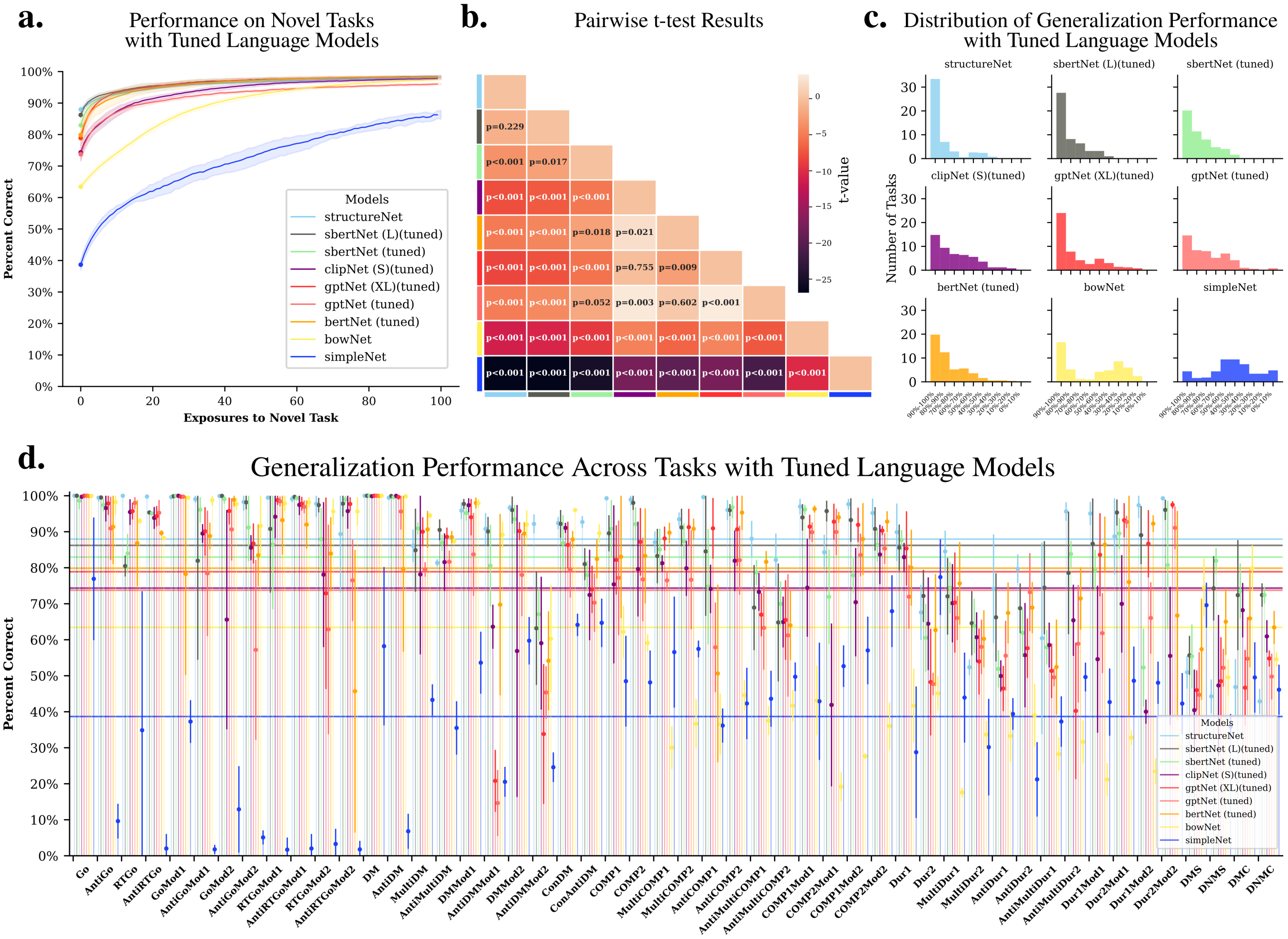


**Supplementary Info. Fig. 4.3**

Full results for models where langauge network is allowed to fine-tune of sensorimotor tasks. We also plot SIMPLENET from main text Fig. 2a. for reference.

##### 4.4 Non-Linear MLP Embedding Maps

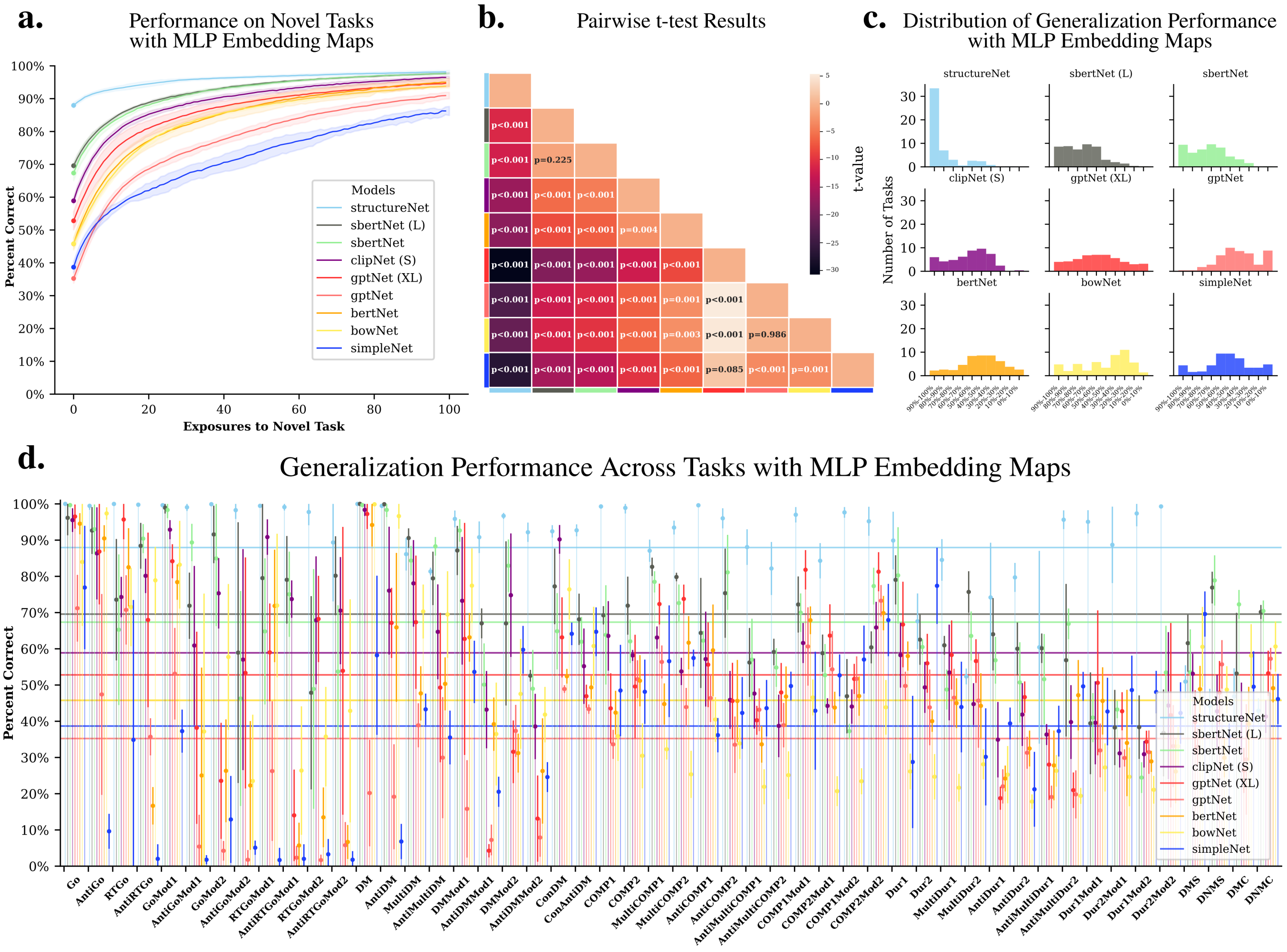


**Supplementary Info. Fig. 4.4**

Here we let a Multi Layer Perception (MLP) with a 256 hidden unit layer and a ReLU nonlinearity map the language model outputs to the final embedding used by the Sensorimotor-RNN in order to determine whether or not this more powerful mapping would improve generalization performance. Performance drops across all our models, suggesting a simple linear mapping is actually advantageous for performance in novel settings. Again, we include SIMPLENET and STRUCTURENET from main text Fig. 2a as comparisons.

##### 4.5 Additional Models

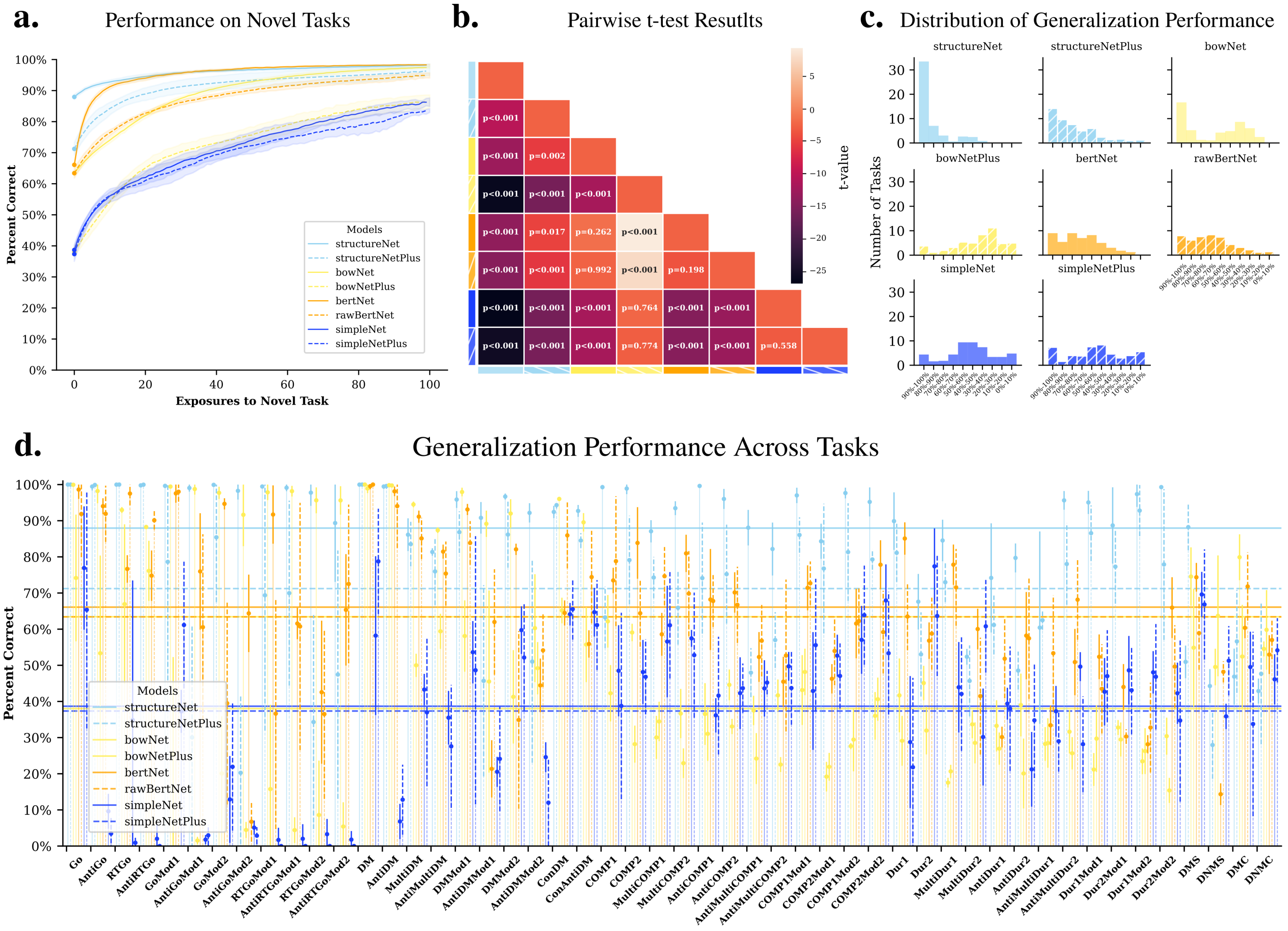


**Supplementary Info. Fig. 4.5**

Since allowing the loss from sensorimotor loss tasks to update the final layers in transformer network improved performance, we tested two additional models that possess some basic very basic linguistic knowledge but otherwise uses a blank slate architectures that are tuned using only the sensorimotor loss. First, we test an extended version of BOWNET called BOWNETPLUS which begins with standard Bag of Words embeddings but processes these embeddings through an additional MLP with 5 layers of 256 hidden units each followed by a ReLU nonlinearity. Our second model, called RAWBERTNET uses BERT word embeddings (which includes information about e.g. the ordering of words in the sentence) and passes these through a single randomly initialized BERT encoder layer. Models are trained using the same procedure as described in the training section of Methods with all weights plastic (versions of RAWBERTNET that used more than one transformer layer failed to reach threshold performance across the task set). Here we also plot results for SIMPLENETPLUS and STRUCTURENETPLUS, again described in the methods, and include standard SIMPLENET, STRUCTURENET, and BERTNET from main text Fig. 2a as comparisons.

#### 5 Compositional Rule Performance

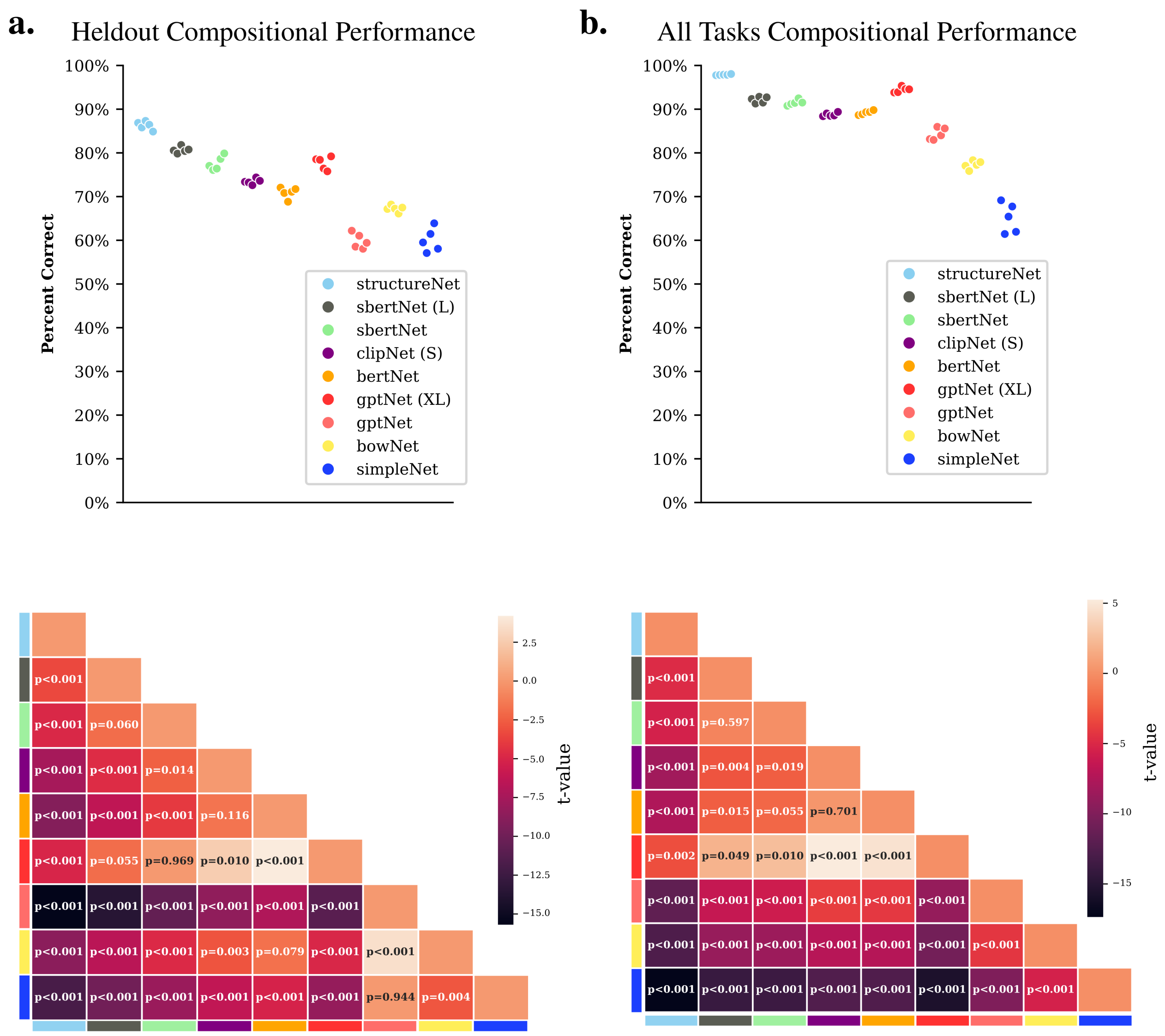


**Supplementary Info. Fig. 5**

We test model performance across tasks where task type information is a simple combination of task information for related tasks. Take 'AntiDMMod1' as an example. For our non-linguistic models the compositional rule for 'AntiDMMod1' is $(rule(\text{AntiDMMod2}) - rule(\text{DMMod2})) + rule(\text{DMMod2})$ where $rule(\text{task})$ denotes the 64-dimensional rule vector corresponding to that task. For our language models, we randomly draw an instruction from the set of each related task, embed the instructions using our language model and then apply the same formula to produce compositional task information. **a.** shows results on tasks that have been held out of training whereas **b.** shows results across tasks using models that have trained on all tasks. The compositional formulas for all tasks are given below. Each point represents average performance across tasks for a given random initialization. Compositional Rules:

'Go' = ('RTGo' - 'AntiRTGo') + 'AntiGo'

'AntiGo' = ('AntiRTGo' - 'RTGo') + 'Go'

'RTGo' = ('Go' - 'AntiGo') + 'AntiRTGo'

'AntiRTGo' = ('AntiGo' - 'Go') + 'RTGo'

'GoMod1' = ('RTGoMod1' - 'AntiRTGoMod1') + 'AntiGoMod1'

'GoMod2' = ('RTGoMod2' - 'AntiRTGoMod2') + 'AntiGoMod2'

'AntiGoMod1' = ('AntiRTGoMod1' - 'RTGoMod1') + 'GoMod1'

'AntiGoMod2' = ('AntiRTGoMod2' - 'RTGoMod2') + 'GoMod2'

'RTGoMod1' = ('GoMod1' - 'AntiGoMod1') + 'AntiRTGoMod1'

'AntiRTGoMod2' = ('AntiGoMod1' - 'GoMod1') + 'RTGoMod1'

'RTGoMod2' = ('GoMod2' - 'AntiGoMod2') + 'AntiRTGoMod2'

'AntiRTGoMod2' = ('AntiGoMod2' - 'GoMod2') + 'RTGoMod2'

'DM' = ('MultiDM' - 'AntiMultiDM') + 'AntiDM'

'AntiDM' = ('AntiMultiDM' - 'MultiDM') + 'DM'

'MultiDM' = ('DM' - 'AntiDM') + 'AntiMultiDM'

'AntiMultiDM' = ('AntiDM' - 'DM') + 'MultiDM'

'ConDM' = ('DM' - 'AntiDM') + 'ConDM'

'ConAntiDM' = ('AntiDM' - 'DM') + 'ConDM'

'DMMod1' = ('DMMod2' - 'AntiDMMod2') + 'AntiDMMod1'

'DMMod2' = ('DMMod1' - 'AntiDMMod1') + 'AntiDMMod2'

'AntiDMMod1' = ('AntiDMMod2' - 'DMMod2') + 'DMMod1'

'AntiDMMod2' = ('AntiDMMod1' - 'DMMod1') + 'DMMod2'

'Dur1' = ('MultiDur1' - 'MultiDur2') + 'Dur2'

'Dur2' = ('MultiDur2' - 'MultiDur1') + 'Dur1'

'MultiDur1' = ('Dur1' - 'Dur2') + 'MultiDur2'

'MultiDur2' = ('Dur2' - 'Dur1') + 'MultiDur1'

'AntiDur1' = ('AntiMultiDur1' - 'AntiMultiDur2') + 'AntiDur2'

'AntiDur2' = ('AntiMultiDur2' - 'AntiMultiDur1') + 'AntiDur1'

'AntiMultiDur1' = ('AntiDur1' - 'AntiDur2') + 'AntiMultiDur2'

'AntiMultiDur2' = ('AntiDur2' - 'AntiDur1') + 'AntiMultiDur1'

'Dur1Mod1' = ('Dur1Mod2' - 'Dur2Mod2') + 'Dur2Mod1'

'Dur1Mod2' = ('Dur1Mod1' - 'Dur2Mod1') + 'Dur2Mod2'

'Dur2Mod1' = ('Dur2Mod2' - 'Dur1Mod2') + 'Dur1Mod1'

'Dur2Mod2' = ('Dur2Mod1' - 'Dur1Mod1') + 'Dur1Mod2'

'COMP1' = ('MultiCOMP1' - 'MultiCOMP2') + 'COMP2'

'COMP2' = ('MultiCOMP2' - 'MultiCOMP1') + 'COMP1'

'MultiCOMP1' = ('COMP1' - 'COMP2') + 'MultiCOMP2'

'MultiCOMP2' = ('COMP2' - 'COMP1') + 'MultiCOMP1'

'AntiCOMP1' = ('AntiMultiCOMP1' - 'AntiMultiCOMP2') + 'AntiCOMP2'

'AntiCOMP2' = ('AntiMultiCOMP2' - 'AntiMultiCOMP1') + 'COMP1'

'AntiMultiCOMP1' = ('AntiCOMP1' - 'AntiCOMP2') + 'AntiMultiCOMP2'

'AntiMultiCOMP2' = ('AntiCOMP2' - 'AntiCOMP1') + 'AntiMultiCOMP1'

'COMP1Mod1' = ('COMP1Mod2' - 'COMP2Mod2') + 'COMP2Mod1'

'COMP1Mod2' = ('COMP1Mod1' - 'COMP2Mod1') + 'COMP2Mod2'

'COMP2Mod1' = ('COMP2Mod2' - 'COMP1Mod2') + 'COMP1Mod1'

'COMP2Mod2' = ('COMP2Mod1' - 'COMP1Mod1') + 'COMP1Mod2'

'DMS' = ('DMC' - 'DNMC') + 'DNMS'

'DNMS' = ('DNMC' - 'DMC') + 'DMS'

'DMC' = ('DMS' - 'DNMS') + 'DNMC'

'DMNC' = ('DNMS' - 'DMS') + 'DMC'

#### 6 Statistical Test Results for Conditional Clause/Deduction Analysis

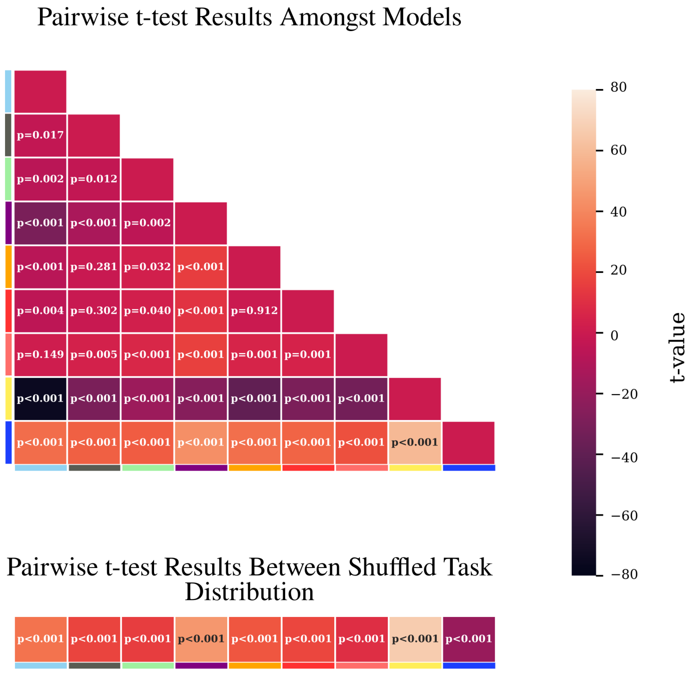


**Supplementary Info. Fig. 6**

Top: Pairwise unequal variance t-test results comparing average differences between tasks with and without conditional clauses/deductive component amongst models compute according to the procedure outlined in Methods of main text. Bottom: t-test results between each model's average difference in performance and a null distribution constructed by randomly shuffling sets of 30 and 20 tasks as outlined in methods.

#### 7 Example Task Variance Plot

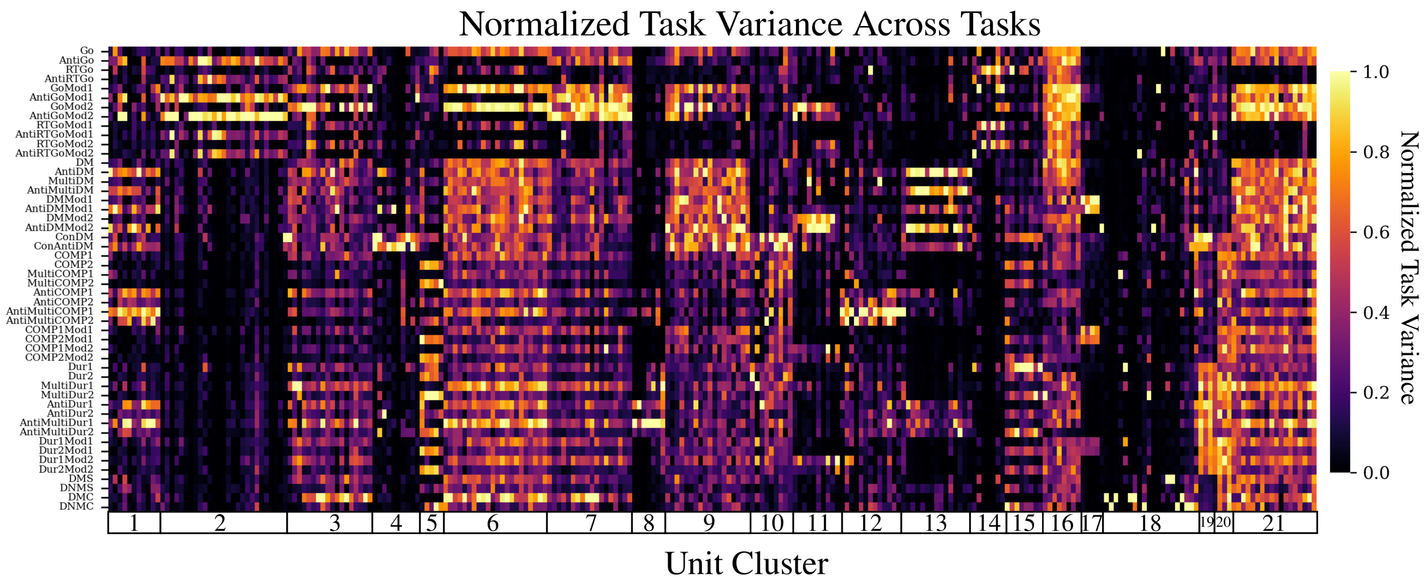


**Supplementary Info. Fig. 7**

The goal of examining possible clustering of individual units across related tasks was to determine if networks did in fact organize themselves according to modularized functional subcomponents. Our procedure follows the one used in Yang et. al. 2019. First, we compute a time averaged variance of unit activity across tasks for 100 different trials for each task. In each case these 100 trials tile the space of possible values for stimulus variables (e.g. 100 even spaced stimulus input directions between 0 and $2\pi$ for 'Go' tasks). For each task, we compute the variance of each model unit at each time point across these trials and then average over all the time steps in the trial after stimulus onset. This results in an overall measure of how much each unit in the network varies in each task. We then normalize each unit variance score across tasks. This results in each unit's task selectivity represented as a 50-dimensional vector where each entry is normalized variance for a given task. We do k-means clustering for each unit's variance score vector. We compute a silhouette score as a measure of clustering quality for k ranging from 2-50. The highest silhouette score determines the number of clusters used to plot the unit clustering. In the above example 'AntiGo', 'Dur1', 'AntiDMMod1', 'ConDM', 'AntiMultiDur2' are held out of training.

#### 8 Training RNN Input Weights Only

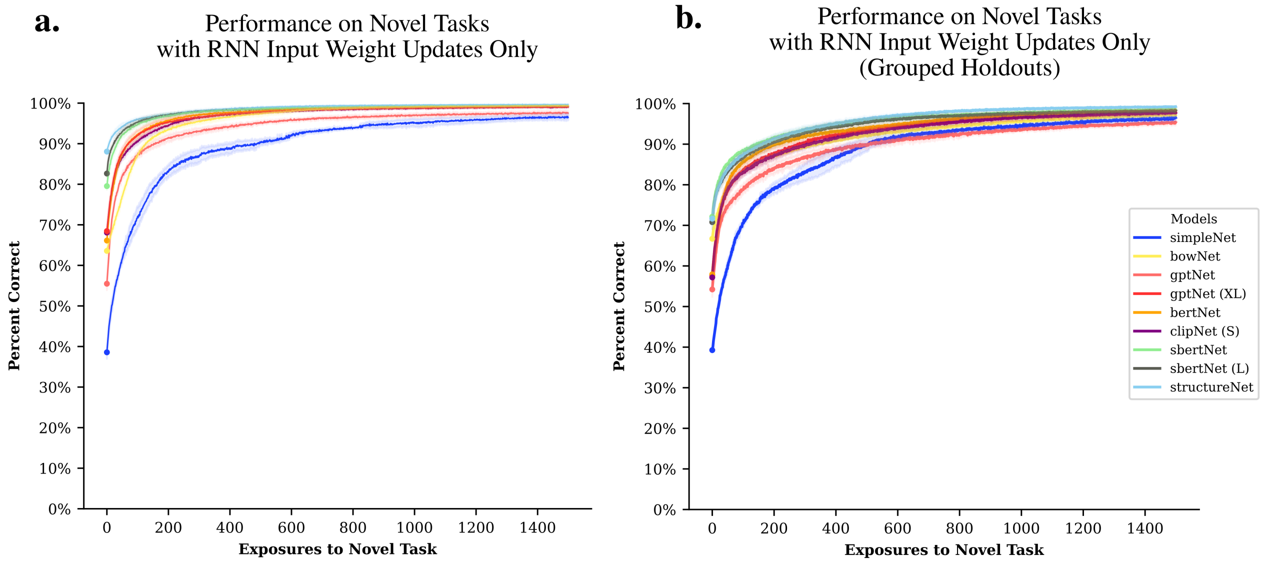


**Supplementary Info. Fig. 8**

Same as Fig. 2a. in the main text except only RNN input weights are plastic. **a.** learning curves for our standard held out setting. **b.** learning curves for models where multiple related tasks are held out during training.

### 9 Additional CCGP Measures

#### 9.1 RNN CCGP Measures

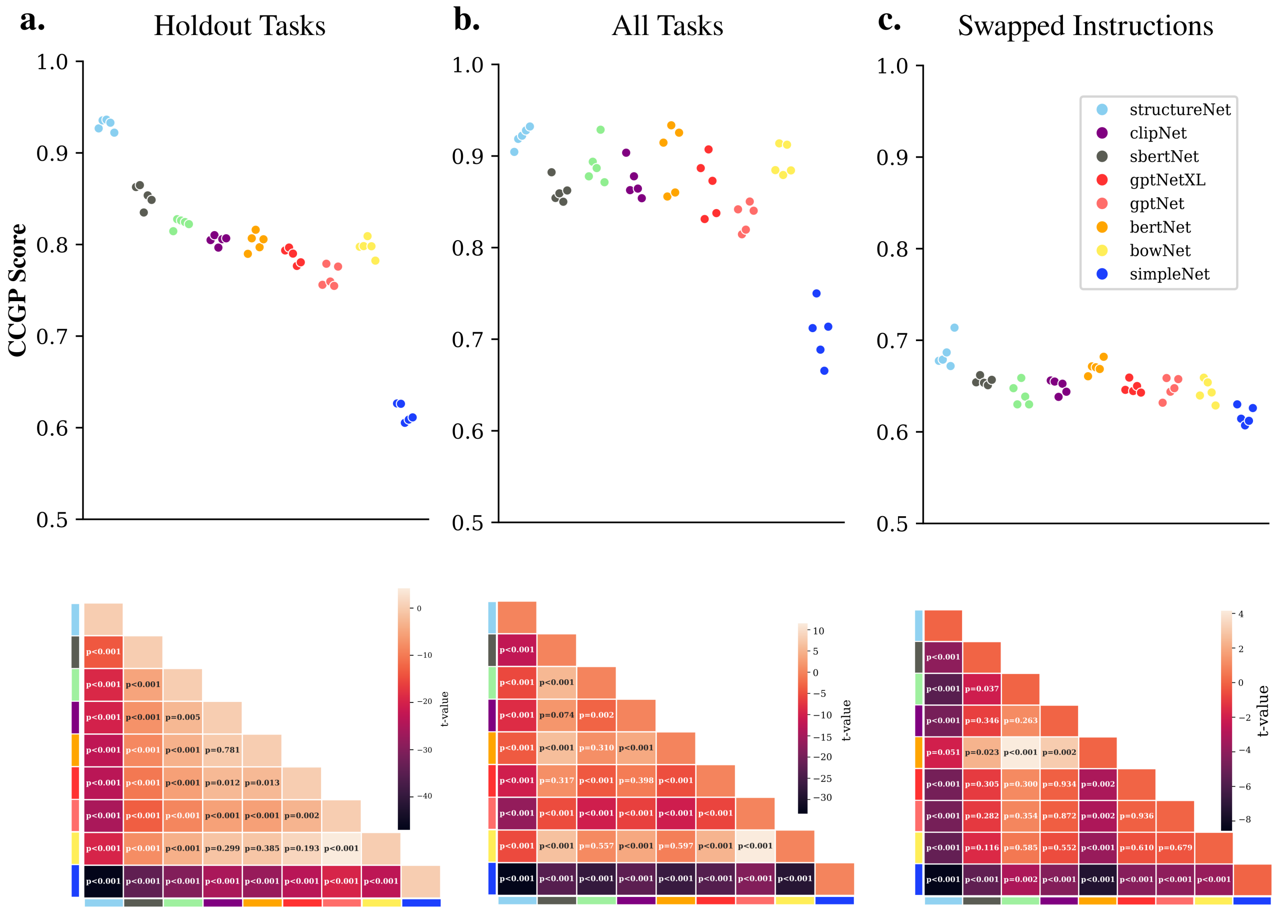


**Supplementary Info. Fig. 9.1**

Additional CCGP measures for different training conditions and inputs for the SensorimotorRNN. For reference we plot Holdout Task which shows the same data as Fig. 3d. "RNN" in the main text (**a.**). All Tasks (**b.**) shows CCGP measures for RNN hidden activity across tasks in networks that have been trained on all tasks (i.e. no held out tasks). Even in this context we see that instructed models tend to have higher CCGP than non-linguistic models. Finally, Swapped Instructions show CCGP measures for RNN hidden activity when instructions are swapped **c.**. Each point represents average CCGP across tasks for a given random initialization.

##### 9.2 Embedding CCGP Measures

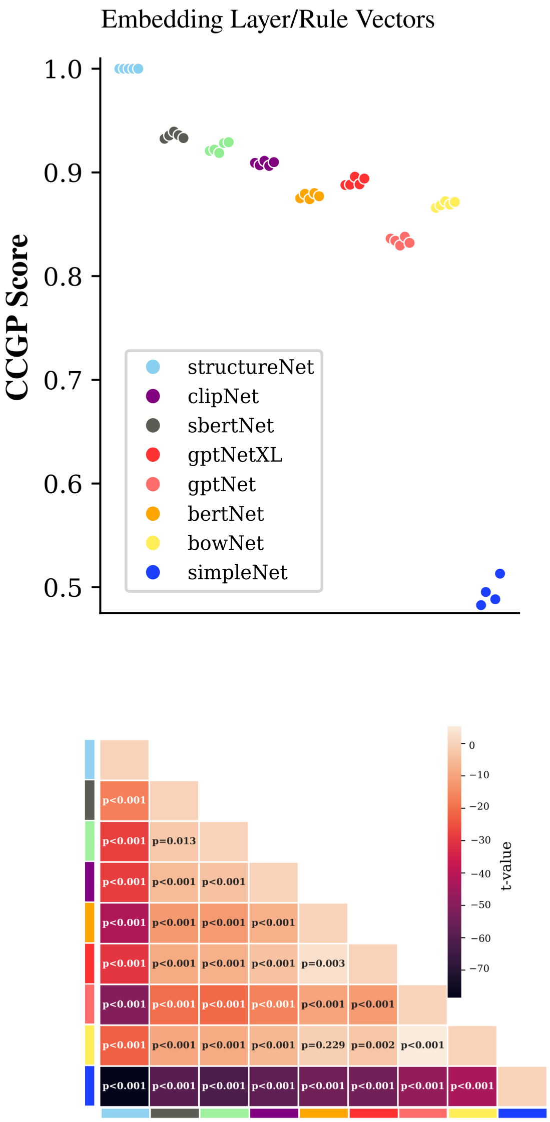


**Supplementary Info. Fig. 9.2**

CCGP for the embedding layer across models.

#### 10 GPT Pooling Comparison

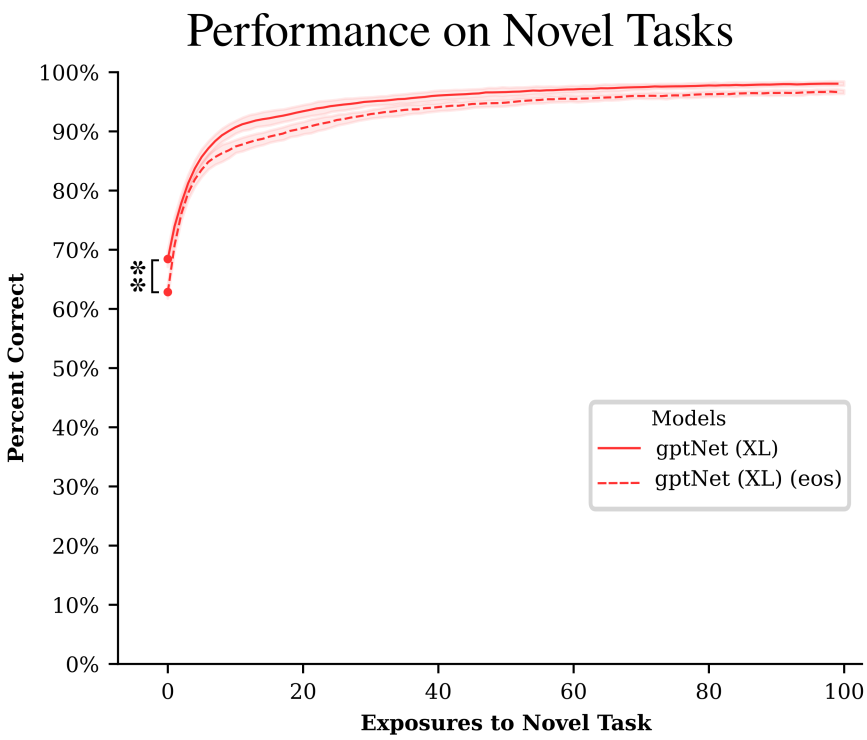


**Supplementary Info. Fig. 10**

Comparison of performance on novel tasks using [eos] embedding method for autoregressive language models (see Method, pre-trained transformers). Using GPT with [eos] sentence embedding failed to achieve even the relaxed performance criteria of 85% across training tasks, hence we only include GPTNETXL (using average pooling, reported in main text) and GPTNETXL (EOS).

### 11 Additional Language Production Tests

#### 11.1 Language Production Performance with Direct from Embeddings

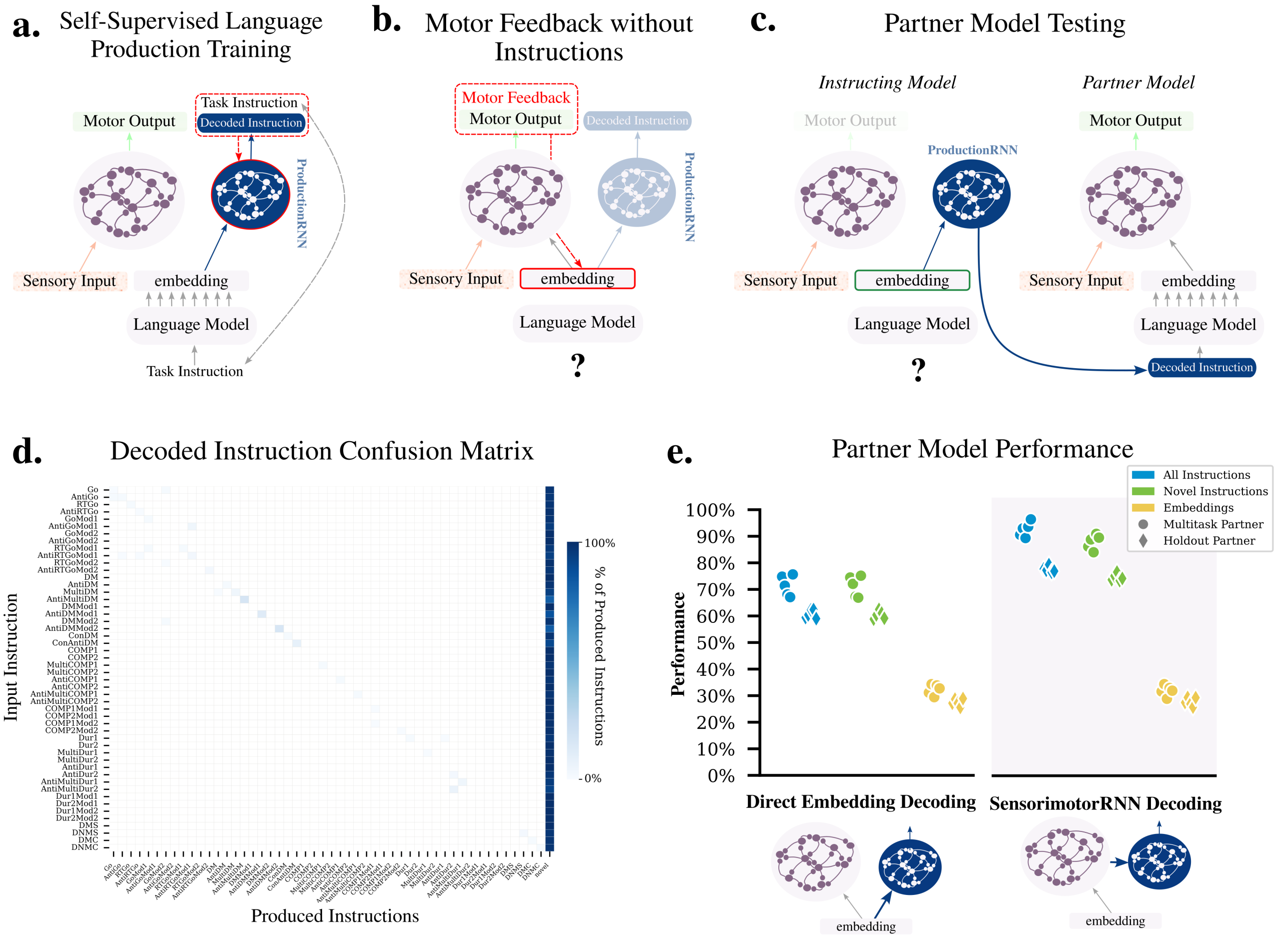


**Supplementary Info. Fig. 11.1**

We also test a version of our language production set up where we map directly from language embeddings to a linguistic output according to the following equations:

$$
\begin{aligned}
h_0^{\text{decoder}} &= Embed(w_{1,k} \dots w_{\mathcal{T},k}) \in \mathbb{R}^{64} \\
h_\tau^{\text{decoder}} &= \text{ProductionRNN}(\hat{w}_{1,k} \dots \hat{w}_{\tau-1,k}; h_0^{\text{decoder}}), && h_\tau^{\text{decoder}} \in \mathbb{R}^{256} \\
p_{\hat{w}_{\tau,k}} &= \text{softmax}(\text{Linear}_{\text{words}}(h_{\tau,k}^{\text{decoder}})) && p_{\hat{w}_{\tau,k}} \in \mathbb{R}^{|vocab|}, \\
\hat{w}_{\tau,k} &= \text{argmax}(p_{\hat{w}_{\tau,k}})
\end{aligned}
$$

Training and evaluation follow the methods outlined in the main text. We find that producing instructions in this manner results in worse performance. This is likely because of a difference in the nature of the training data each production method is exposed to. In the case of a production model that trains on RNN hiddens states, the training set is as diverse as the number of conditions that make up a trial. When decoding directly from the embedding space, the training set is only as large as the number of instructions for a given task. Hence, when this model attempts to process embeddings found by applying gradient descent based on the sensorimotor loss, the resultant points may very well lie out of distribution relative to this small set of training examples.

##### 11.2 Language Production Performance with STRUCTURENET

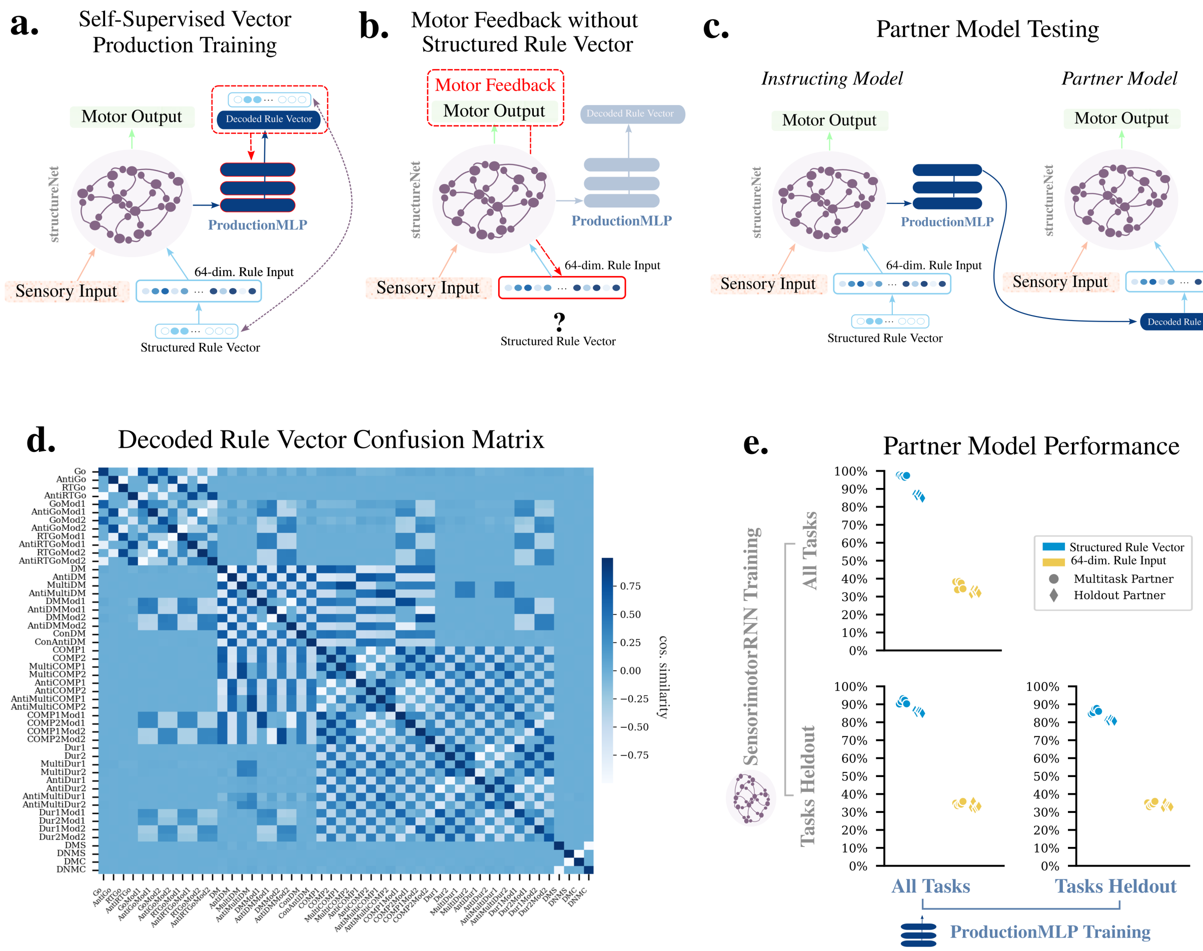


**Supplementary Info. Fig. 11.2**

We also test the ability of STRUCTURENET to perform in the language production setting. Overall, the set up is largely the same as in Fig. 5 of the main text. The key difference is that since STRUCTURENET takes a task vector instead of a sequence of words as its task-type information, we use a ProductionMLP instead of a ProductionRNN. The ProductionMLP consists of a 3 layer MLP with 256 hiddens units per layer and ReLU nonlinearities. Let $X^i$ denote the sensory inputs for a given task $i$ and $RuleVec_i$ denote the simple 10 dimensional rule vector for task $i$ and let $RuleBasis$ denote set of 10 orthogonal 64-dimensional vectors unique to each instance of STRUCTURENET. Then production proceeds according to the following equations.

$$RuleInput = RuleVec_i \cdot RuleBasis \qquad RuleInput \in RuleInput\mathbb{R}^{64}$$

$$h_T^{sm} = \text{SensorimotorRNN}(X^i, RuleInput)) \qquad h_T^{sm} \in \mathbb{R}^{T \times 256}$$

$$sm\_out = (\text{mean}_T(h_T^{sm}), \text{max}_T(h_T^{sm})) \qquad sm\_out \in \mathbb{R}^{512}$$

$$RuleVec_i' = \text{ProductionMLP}(sm\_out) \qquad RuleVec_i' \in \mathbb{R}^{10}$$

Self supervised learning proceeds according to the same set of hyperparameters described in the Methods of the main text, with the use of a MSE loss on between produced $RuleVec_i'$ and ground truth $RuleVec_i$. Training of embedding vectors with motor feedback and evaluation of partner model performance use the same procedures outlined in the main text.

##### 11.3 Language Production Performance Statistical Test

Results for statistic t-tests over different training regimes (e.g. partner all sm all prod. all full instructs labels that partner model, sm network, and prod. network were trained on all tasks and the partner model received all instructions).

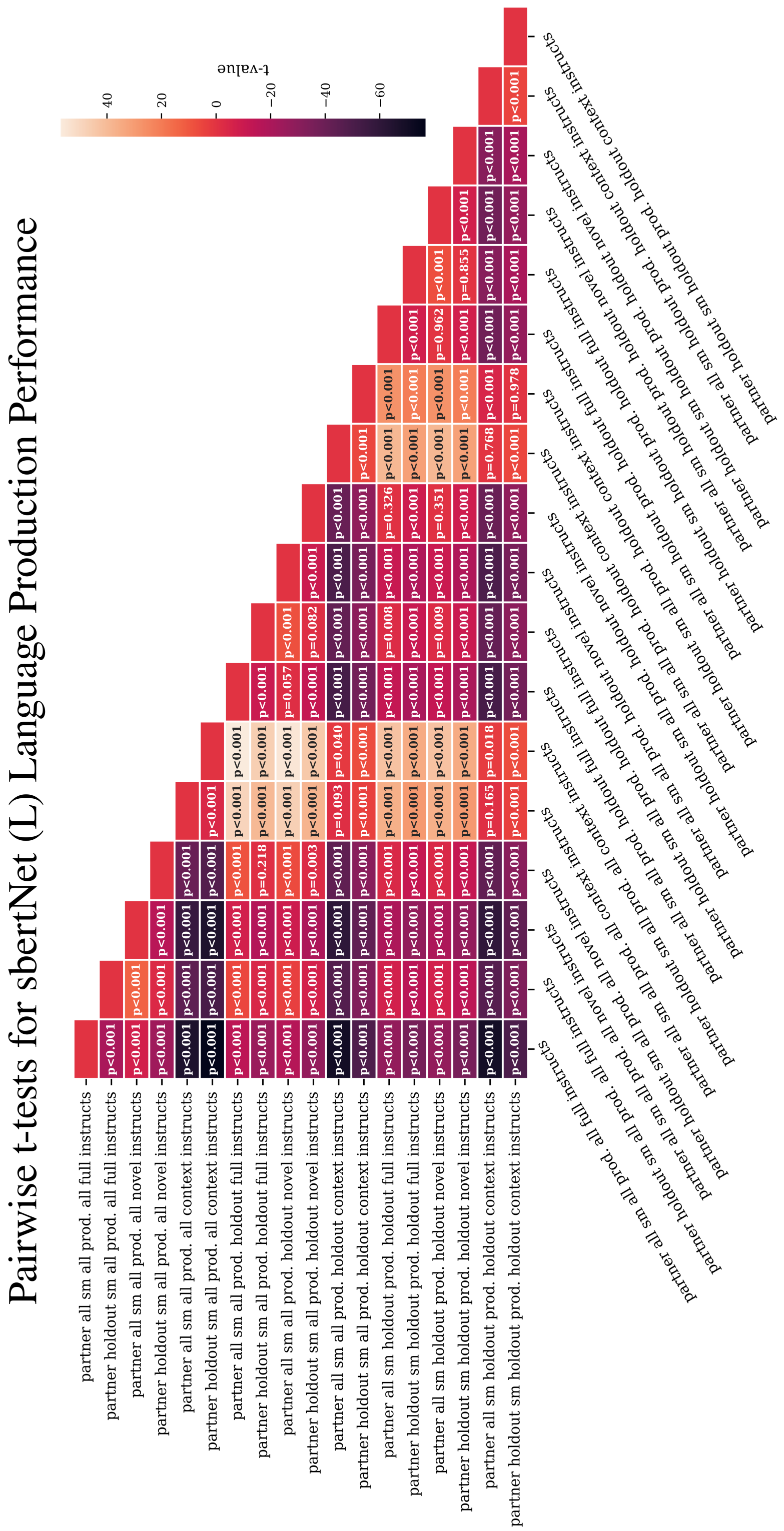


**Supplementary Info. Fig. 11.3.1**

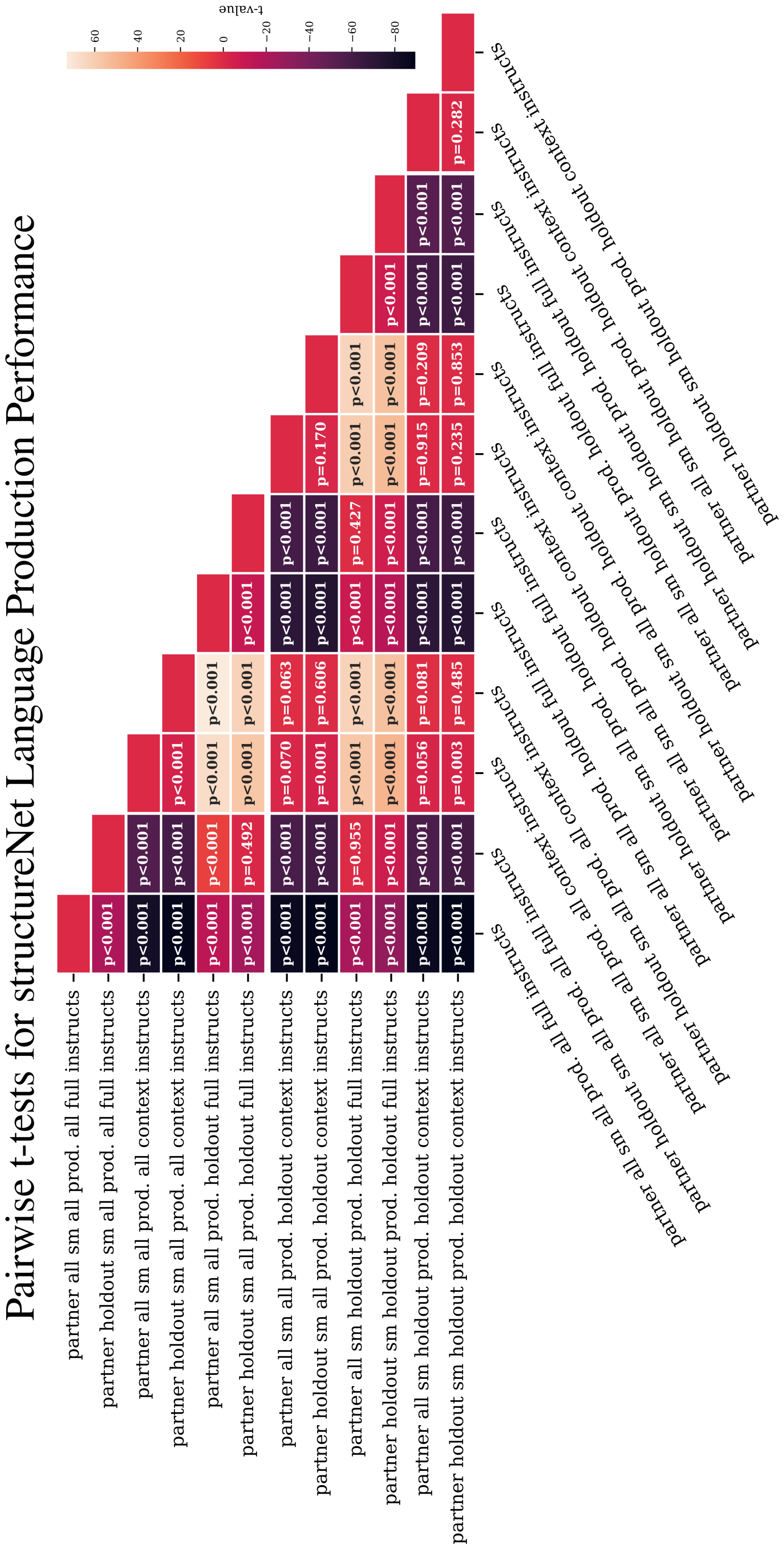


**Supplementary Info. Fig. 11.3.2**

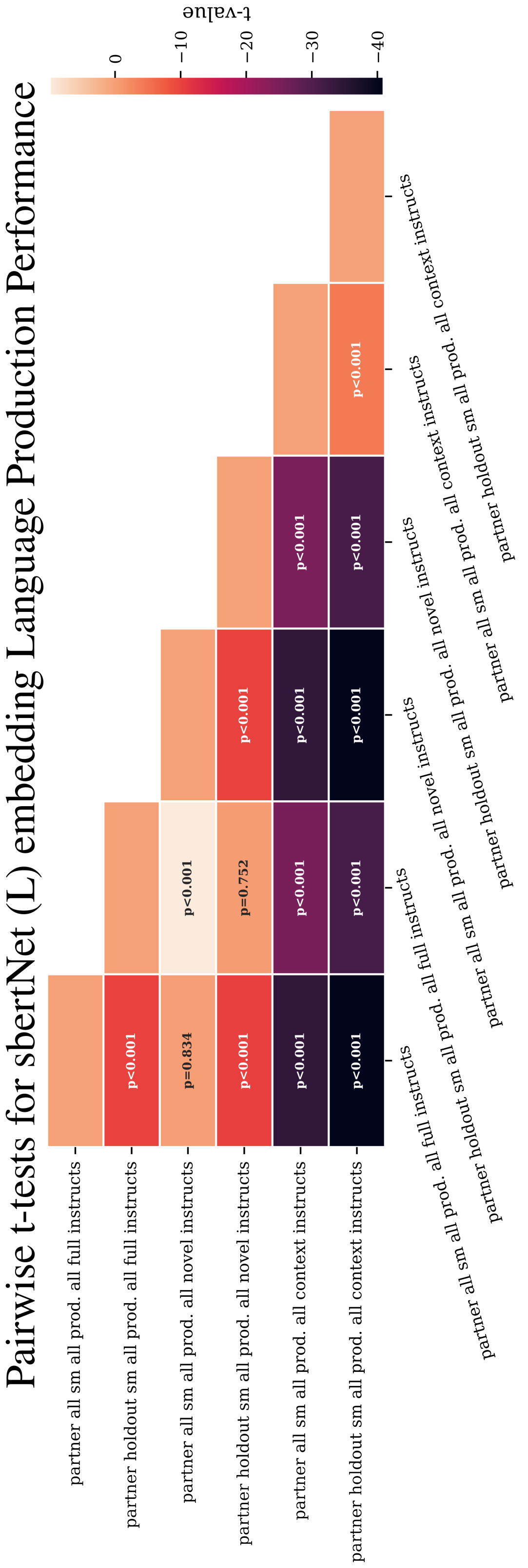


**Supplementary Info. Fig. 11.3.3**

#### 12 CCGP Dichotomies

***Go vs. Anti Dichotomy:***

{(Go, AntiGo), (RTGo, AntiRTGo), (GoMod1, AntiGoMod1), (GoMod2, AntiGoMod2), (RTGoMod1, AntiRTGoMod1), (RTGoMod2, AntiRTGoMod2)}

***Standard vs. RT:***

{(Go, RTGo), (GoMod1, RTGoMod1), (GoMod2, RTGoMod2), (AntiGo, AntiRTGo),(AntiGoMod1, AntiRTGoMod1), (AntiGoMod2, AntiRTGoMod2)}

***Weakest vs. Strongest Dichotomy:***

{(DM, AntiDM), (DMMod1, AntiDMMod1), (DMMod2, AntiDMMod2), (MultiDM, AntiMultiDM), (ConDM, ConAntiDM), (COMP1, AntiCOMP1), (COMP2, AntiCOMP2), (MultiCOMP1, AntiMultiCOMP1), (MultiCOMP2, AntiMultiCOMP2) }

***Longest vs. Shortest:***

{(Dur1, AntiDur1), (Dur2, AntiDur2), (MultiDur1, AntiMultiDur1), (MultiDur2, AntiMultiDur2)}

***First Stim. vs. Second Stim.***

{(Dur1, Dur2), (AntiDur1, AntiDur2), (MultiDur1, MultiDur2), (AntiMultiDur1, AntiMultiDur2), (Dur1Mod1, Dur2Mod1), (Dur1Mod2, Dur2Mod2), (COMP1, COMP2), (MultiCOMP1, MultiCOMP2), (AntiCOMP1, AntiCOMP2), (AntiMultiCOMP1, AntiMultiCOMP2), (COMP1Mod1, COMP2Mod1), (COMP1Mod2, COMP2Mod2)}

***Stim. Match vs. Category Match***

{(DMS, DNMS)}
{(DMC, DNMC)}

***Matching vs. Non-Matching***

{(DMS, DMC)}
{(DNMS, DNMC)}

***Mod1 vs . Mod2***

{(GoMod1, GoMod2), (AntiGoMod1, AntiGoMod2), (RTGoMod1, RTGoMod2), (AntiRTGoMod1, AntiRTGoMod2), (DMMod1, DMMod2), (AntiDMMod1, AntiDMMod2), (COMP1Mod1, COMP1Mod2), (COMP2Mod1, COMP2Mod2), (Dur1Mod1, Dur1Mod2), (Dur2Mod1, Dur2Mod2)}

#### 13 Task Instructions

**Go:**

*Train*

'respond in the direction of the stimulus'
'respond with the same direction as the displayed orientation'
'choose the orientation displayed'
'respond with the same orientation'
'go in the presented direction'
'select the displayed orientation'
'pick the displayed orientation'
'copy the direction displayed'
'select the direction of presented stimulus'
'choose the direction of the stimulus'
'respond with the same orientation as the presented stimulus'
'go in the orientation indicated by the stimulus'
'go in the direction presented on the display'
'choose the displayed orientation'
'opt for the same orientation as indicated by the stimulus'

*Validation*

'select the presented direction'
'respond in the direction shown'
'respond in the direction displayed'
'respond with the displayed orientation'
'choose the presented direction'

**RTGo:**

*Train*

'respond in direction of the stimulus immediately'
'respond with the displayed orientation at stimulus onset'
'choose the orientation displayed immediately'
'respond with the same direction immediately'
'go in the same direction immediately'
'go in the displayed direction at as soon as the stimulus appears'
'respond with the identical orientation immediately'
'select the orientation displayed at stimulus onset'
'choose the orientation displayed as soon as the stimulus is shown'
'choose the same direction as the stimulus immediately'
'pick the identical direction displayed as soon as it appears'
'as soon as the stimulus appears select the same direction'
'when the stimulus is shown respond in the equivalent direction'
'choose the direction that appears on the display immediately'
'select the presented direction when the stimulus is shown'

*Validation*

'choose the direction shown immediately'
'select the direction presented immediately'
'select the displayed orientation as soon as stimulus appears'
'choose the direction presented immediately'
'immediately respond to the presented orientation'

**AntiGo:**

*Train*

'respond with the opposite orientation'
'respond in the reverse direction'
'choose the opposite of the displayed direction'
'choose the opposite direction'
'respond with the reverse orientation'
'go in reverse of the displayed orientation'
'opt for the reverse of the presented stimulus'
'select the reverse of the displayed orientation'
'respond with the reverse of the displayed stimulus'
'opt for the opposite of the presented direction'
'pick the opposite of presented direction'
'select the reverse of presented stimulus'
'pick the reverse of the stimulus displayed'
'select the opposite of the displayed direction'
'go in the opposite direction'

*Validation*

'select the reverse of the displayed orientation'
'choose the opposite of the presented direction'
'respond with the reverse orientation'
'go in the opposite of displayed orientation'
'pick the reverse of the stimulus orientation'

**AntiRTGo:**

*Train*

'respond with the opposite of the stimulus immediately'
'opt for the opposite direction at stimulus onset'
'choose the opposite of the displayed direction at stimulus onset'
'select the reverse orientation immediately'
'respond with the reverse direction immediately'
'go in the reverse of the orientation as soon as the stimulus appears'
'choose the reverse direction immediately'
'opt for the opposite of the orientation immediately'
'go in the reverse of the direction as soon as stimulus appears'
'as soon as stimulus appears respond in the opposite direction'
'select the reverse orientation at stimulus onset'
'choose the reverse orientation of the one displayed at stimulus onset'
'respond in the converse of the displayed direction as soon as the stimulus appears'
'at stimulus onset go in the reverse direction'
'select the reverse of the displayed direction at stimulus onset'

*Validation*

'respond with the reverse of the stimulus direction as quickly as possible'
'select the reverse of the displayed direction as soon as possible'
'respond with the reverse orientation as soon as it is shown'
'choose the opposite of displayed orientation at stimulus onset'
'immediately respond in the opposing direction as is soon as it appears'

**GoMod1:**

*Train*

'respond to the direction displayed in the first modality'
'opt for the orientation of the stimulus that appears in the first modality'
'select the direction corresponding to the stimulus in the first modality'
'attend only to the stimulus in the first modality and respond in that direction'
'focus on the first modality and select the stimulus that appears'
'attend to the first modality and go in the direction of the displayed stimulus'
'select the orientation that appears in the first modality'
'opt for the direction displayed in the first modality'
'focus on the first modality and respond to the orientation displayed'

'choose the stimulus presented in the first modality'
'attend to the first modality and pick the displayed direction'
'select the orientation displayed in the first modality'
'focus only on the first modality and choose the shown orientation'
'opt for the direction of the stimulus that appears in the first modality'
'go in the direction that appears in the first modality'

*Validation*

'choose the orientation in the first modality'
'opt for the stimulus that appears in the first modality'
'only consider the first modality and respond in to the displayed direction'
'go in the direction stimulus in the first modality'
'concentrate only on the first modality and choose the direction displayed there'

**GoMod2:**

*Train*

'pay attention only to the second modality and respond to the direction displayed there'
'focus on the second modality and select the displayed direction'
'select the stimulus displayed in the second modality'
'respond to the orientation that appears in the second modality'
'attend to the second modality and respond to the displayed stimulus'
'opt for the orientation displayed in the second modality'
'go in the direction that appears in the second modality'
'focus only on the second modality and go in the direction of the stimulus displayed there'
'pick the direction that appears in the second modality'
'respond in the direction displayed in the second modality'
'attend only to the second modality and select the displayed orientation'
'opt for the direction of the stimulus displayed in the second modality'
'pick the stimulus orientation that appears in the second modality'
'choose the orientation that appears in the second modality'
'focus on the stimulus displayed in the second modality and respond there'

*Validation*

'respond to the stimulus in the second modality'
'select the direction that is displayed in the second modality'
'only consider the second modality and choose the displayed direction'
'opt for the orientation in the second modality'
'concentrate only on the second modality and select the direction that appears there'

**AntiGoMod1:**

*Train*

'focus only on the first modality and respond to the opposite of the displayed direction'
'select the reverse of the orientation presented in the first modality'
'attend to the first modality and respond in the reverse of the displayed direction'
'choose the opposite of the orientation that appears in the first modality'
'pay attention only to the first modality and respond in the opposite of the displayed direction'
'opt for the opposite of the direction displayed in the first modality'
'attend to the stimulus in the first modality and go in the opposite direction'
'select the opposite of the direction that appears in the first modality'
'choose the reverse of the orientation that is shown in the first modality'
'focus on the first modality and respond in the opposite direction of the stimulus displayed there'
'focus on the first modality and select the opposite of the displayed stimulus'
'opt for the reverse of the stimulus orientation displayed in the first stimulus'
'choose the opposite of the stimulus orientation that appears in the first modality'
'attend to the first modality and pick the reverse of the direction displayed there'
'pay attention only to the first modality and choose the reverse of the direction displayed'

*Validation*

'go in the opposite direction of the stimulus presented in the first modality'
'concentrate only on the first modality and respond with the opposite of the displayed direction'
'select the opposite of the direction displayed in the first modality'
'pick the direction opposite of the one displayed in the first modality'
'only consider stimulus in the first modality and respond in the opposite direction'

**AntiGoMod2:**

*Train*

'pay attention only to the second modality and respond in the reverse direction'
'respond to the opposite of the orientation displayed in the second modality'
'select the opposite of the direction displayed in the second modality'
'attend to the stimulus in the second modality and respond in the opposite direction'
'focus on the second modality and respond in the opposite of the direction that appears there'
'choose the opposite of the stimulus presented in the second modality'
'opt for the opposite of the orientation presented in the second modality'
'respond in the reverse of the stimulus orientation from the second modality'
'pick the reverse of the stimulus displayed in the second modality'
'attend to the second modality and select the opposite of the direction displayed'
'focus on the second modality and choose the reverse of the displayed direction'
'select the opposite of the orientation displayed in the second modality'
'choose the opposite of the orientation displayed in the second modality'
'attend only to the second modality and pick the reverse of the presented orientation'
'choose the opposite of the direction of the stimulus presented in the second modality'

*Validation*

'concentrate exclusively on the second modality and go in the reverse of the displayed orientation'
'opt for the opposite of the direction in the second modality'
'go in the reverse of the direction displayed in the second modality'
'only consider the second modality and respond in the reverse direction'
'pick the opposite of the orientation in the second modality'

**RTGoMod1:**

*Train*

'focus on the first modality and select the stimulus immediately'
'attend only to the first modality and choose the direction at stimulus onset'
'at stimulus onset select the direction that appears in the first modality'
'pay attention only to the first modality and select the orientation immediately'
'focus only on the first modality and choose the displayed orientation at stimulus onset'
'choose the direction displayed in the first modality at stimulus onset'
'select the orientation in the first modality immediately'
'attend to the first modality and go in the displayed direction immediately'
'focus on the first modality and opt for the orientation at stimulus onset'
'pay attention to the first modality and select the direction immediately'
'immediately respond to the direction in the first modality'
'select the orientation that appears in the first modality at stimulus onset'
'attend to the first modality and select the displayed direction immediately'
'go in the direction displayed in the first modality at stimulus onset'
'attend to the first modality and pick the direction immediately'

*Validation*

'concentrate only on the first modality and select the displayed stimulus immediately'
'choose the direction in the first modality as soon as it appears'
'immediately choose the orientation in the first modality'
'focus exclusively on the first modality and respond to the stimulus immediately'
'select the direction in the first modality at stimulus onset'

**RTGoMod2:**

*Train*

'focus only on the second modality and choose the stimulus immediately'
'attend to the second modality and select the orientation at stimulus onset'
'pay attention to the second modality and go in the displayed direction immediately'
'select the stimulus in the second modality at stimulus onset'
'focus on the second modality and select the displayed orientation immediately'
'attend to the second modality and respond to the displayed direction immediately'
'choose the stimulus in the second modality immediately'
'at stimulus onset choose the orientation in the second modality'
'focus only on the second modality and choose the orientation at stimulus onset'
'pay attention to the second modality and pick the stimulus immediately'
'attend to the second modality and respond to the displayed direction at stimulus onset'
'pay attention to the second modality and opt for the displayed orientation at stimulus onset'
'respond to the direction in the second modality at stimulus onset'
'go in the direction displayed in the second modality immediately'
'opt for the stimulus in the second modality immediately'

*Validation*

'attend exclusively to the second modality and choose the stimulus as soon as it appears'
'as soon as the stimulus appears select the direction in the second modality'
'respond to the stimulus that is shown in the second modality immediately'
'only consider the second modality and select the displayed direction as soon as it appears'
'select the direction shown in the second modality as soon as it appears'

**AntiRTGoMod1:**

*Train*

'focus on the first modality and select the opposite of the stimulus immediately'
'opt for the reverse of the orientation in the first modality at stimulus onset'
'choose the opposite of the direction in the first modality immediately'
'attend only to the first modality and select the reverse of the stimulus immediately'
'pay attention only to the first modality and opt for the opposite of the orientation at stimulus onset'
'attend only to the first modality and respond with the opposite direction at stimulus onset'
'go in the opposite of the orientation displayed in the first modality at stimulus onset'
'focus on the first modality and respond in the reverse of the stimulus immediately'
'pay attention to the first modality and go in the reverse of the stimulus immediately'
'focus on the first modality and choose the reverse of the orientation at stimulus onset'
'attend only to the first modality and opt for the reverse direction at stimulus onset'
'pay attention to the first modality and respond to the reverse of the stimulus immediately'
'go in the reverse of the direction in the first modality at stimulus onset'
'respond with the opposite of the stimulus in the first modality immediately'
'attend to the first modality and select the reverse of the orientation at stimulus onset'

*Validation*

'attend exclusively to the first modality and respond to the opposite direction of the stimulus as soon as it appears', 'respond to the opposite of the stimulus in the first modality immediately', 'only consider stimuli in the first modality and choose the opposite of the displayed direction at stimulus onset', 'go in the opposite of the orientation displayed in the first modality as soon as it appears', 'select the reverse of the direction displayed in the first modality immediately'

**AntiRTGoMod2:**

*Train*

'focus only on the second modality and choose the opposite of the stimulus immediately'
'attend only to the second modality and go in the reverse of the direction at stimulus onset'
'pay attention to the second modality and respond with the reverse of the stimulus immediately'
'focus only on the second modality and select the opposite of the orientation at stimulus onset'
'attend to the second modality and opt for the reverse of the displayed direction immediately'
'pay attention to the second modality and pick the opposite orientation at stimulus onset'
'go in the reverse of the direction in the second modality at stimulus onset'
'opt for the opposite of the stimulus in the second modality immediately'
'choose the reverse of the orientation in the second modality at stimulus onset'
'pay attention to the second modality and choose the opposite of the stimulus immediately'
'attend to the second modality and opt for the reverse direction at stimulus onset'
'respond in the opposite of the direction in the second modality at stimulus onset'
'focus on the second modality and select the reverse of the displayed stimulus immediately'
'attend to the second modality and go in the opposite direction at stimulus onset'
'go in the opposite of the stimulus in the second modality immediately'

*Validation*

'attend exclusively to the second modality and go in the opposite of the direction displayed there at once'
'concentrate only on the second modality and respond to the reverse of the stimulus immediately'
'choose the opposite of the orientation in the second modality at stimulus onset'
'opt for the reverse of direction in the second modality immediately'
'select the opposite of the stimulus that is displayed in the second modality as soon as it is shown'

**DM:**

*Train*

'respond in the direction of highest intensity'
'choose the strongest stimulus'
'go in the direction of the stimulus with maximal strength'
'respond in the direction of the strongest stimulus'
'choose the most intense stimulus'
'respond in direction of greatest stimulus strength'
'choose the stimulus presented with highest intensity'
'go in direction of greatest intensity'
'choose the orientation with largest strength'
'select the direction presented with highest intensity'
'respond to the stimulus with maximal intensity'
'select the stimulus orientation with greatest strength'
'select the stimulus with greatest intensity'
'respond with the orientation presented with greatest strength'
'choose the most intensely displayed orientation'

*Validation*

'select the stimulus of greatest strength'
'choose the direction with maximal stimulus intensity'
'pick the direction with stimulus of greatest strength'
'select the orientation displayed with greatest intensity'
'go in the direction of the most intense stimulus'

**AntiDM:**

*Train*

'respond in the direction of minimal strength'
'choose the weakest stimulus'
'go in the direction of stimulus with least intensity'
'select the stimulus presented with the lowest strength'
'go in the direction presented with lowest intensity'
'choose the stimulus with minimal strength'
'respond to the direction presented with the weakest strength'
'pick the stimulus with least strength'
'select the orientation presented with minimal strength'
'choose the stimulus with lowest intensity'
'go in the direction presented with weakest intensity'
'pick the weakest direction'
'select the orientation with lowest intensity'
'pick the least intense direction'
'respond in the direction of minimal intensity'

*Validation*

'choose the direction with lowest value'
'respond in the direction that is presented with least intensity'
'pick the orientation with the lowest strength'
'go in the direction with weakest presentation strength'
'select the direction with minimal intensity'

**MultiDM:**

*Train*

'respond to the combined greatest value between two stimuli'
'choose the direction with highest average intensity between two modalities'
'select the direction with highest overall strength between two stimuli'
'respond in the direction of highest combined stimulus strength'
'go in the direction with largest joint intensity between stimuli'
'select the orientation with highest average strength over both modalities'
'go in the direction of highest combined stimulus value'
'choose the direction representing the highest integrated stimulus strength'
'respond in the direction with greatest sum over two displayed stimulus values'
'select the direction with greatest average value across modalities'
'choose the orientation with highest intensity integrated over modalities'
'respond with the orientation displayed with maximal strength averaged across modalities'
'go in the direction of stimulus with maximal intensity over both modalities'
'pick the orientation with greatest value combined over modalities'
'pick the orientation presented with highest average intensity'

*Validation*

'choose the stimuli with highest intensity averaged over modalities'
'select the orientation with greatest combined strength over modalities'
'go in direction with highest joint intensity between stimuli'
'pick the direction of stimulus with greatest intensity between two modalities'
'select the stimuli with maximal combined strength over both modalities'

**AntiMultiDM:**

*Train*

'respond in the direction with lowest combined value between two stimuli'
'select the orientation with least average intensity between two modalities'
'select the direction with weakest average value between two stimuli'
'respond in the direction of minimal combined stimulus strength'
'go in the direction of smallest intensity integrated over both two modalities'
'choose the orientation with lowest average value over both modalities'
'choose the orientation which has the weakest joint intensity over modalities'
'respond in the direction which has the lowest combined intensity'
'select the orientation with the minimal strength over modalities'
'go in the direction with weakest average value across modalities'
'pick the direction displayed with least intensity across modalities'
'pick the stimulus displayed with the lowest combined strength between modalities'
'choose the stimulus presented with weakest intensity averaged over modalities'
'respond in the direction with minimal strength across both modalities'
'select the direction which has the least combined strength'

*Validation*

'pick the direction with minimal average value over modalities'
'select the orientation with lowest combined strength'
'go in the direction of the stimuli with weakest overall value between modalities'
'choose the direction with lowest presentation strength over modalities'
'select the orientation with weakest intensity over both modalities'

**ConDM:**

*Train*

'respond to the the strongest stimulus only if you are confident otherwise do not respond'
'if you are sure of the correct answer respond to the strongest stimulus otherwise do not respond'
'select the orientation displayed with greatest strength if you have high confidence otherwise do not respond'
'choose the direction with highest intensity if you are confident in your decision otherwise do not respond'
'if you are confident about the answer respond to the orientation with maximal intensity otherwise do not respond'
'go in the direction of greatest stimulus strength if you have high confidence in your answer otherwise do not respond'
'respond to the stimulus presented with highest intensity if you are sure of your decision otherwise do not respond'
'if you are sure of the result select the stronger direction otherwise do not respond'
'if you are confident about the result pick the most intense stimulus otherwise do not respond'
'pick the orientation that appears with greatest strength if you have high confidence otherwise do not respond'
'opt for the direction displayed with highest intensity if you are sure of your answer otherwise do not respond'
'if you are confident in your answer select the stimulus with highest intensity otherwise do not respond'
'opt for the most intensely displayed direction if you are sure about your decision otherwise do not respond'
'respond in the direction of greatest strength if you are confident otherwise do not respond'
'choose the strongest orientation if you have high confidence otherwise do not respond'

*Validation*

'choose the strongest stimulus if you are convinced of your answer otherwise do not respond'
'select the stimulus with greatest strength if there is no doubt in your mind otherwise do not respond'
'if you are positive then choose the direction presented with greatest strength otherwise do not respond'
'opt for the strongest direction displayed if you are certain about your answer otherwise do not respond'
'respond to the orientation with greatest strength if you are certain of the answer otherwise do not respond'

**ConAntiDM:**

*Train*

'opt for the stimulus presented with least strength if you are confident in your answer otherwise do not respond'
'if you have high confidence select the stimulus displayed with minimal intensity otherwise do not respond'
'if you are confident in your answer pick the weakest stimulus otherwise do not respond'
'choose the orientation that appears with lowest intensity if you are confident otherwise do not respond'
'pick the weakest direction if you are confident in your answer otherwise do not respond'
'respond to the stimulus with minimal strength if you are sure about your answer otherwise do not respond'
'go in the direction of the stimulus presented with least strength if you are sure about your decision otherwise do not respond'
'if you have high confidence choose the orientation with least intensity otherwise do not respond'
'if you are sure about your answer select the direction with least strength otherwise do not respond'
'pick the stimulus which appears with minimal strength if you are confident in your answer otherwise do not respond'
'select the least intense direction if you have high confidence otherwise do not respond'
'choose the stimulus presented with least strength if you are confident in your decision otherwise do not respond'
'respond to the orientation with lowest strength if you are sure of your answer otherwise do not respond'
'if you have high confidence choose the orientation with least strength otherwise do not respond'
'opt for the weakest direction if you have high confidence otherwise do not respond'

*Validation*

'select the weakest direction if you are convinced of the answer otherwise do not respond'
'respond to the stimulus with weakest strength if you are positive otherwise do not respond'
'if you are certain of your answer select the weaker of the two directions otherwise do not respond'
'if there is no doubt in your mind choose the stimulus with least strength otherwise do not respond'
'opt for direction with lowest strength if you are certain about your answer otherwise do not respond'

**DMMod1:**

*Train*

'select the orientation in the first modality that is strongest'
'choose the most intense stimulus that appears in the first modality'
'attend to the first modality and select the orientation displayed with highest intensity'
'focus only on the first modality and choose the direction with greatest strength'
'opt for the direction with largest strength in the first modality'
'select the stimulus with maximal intensity in the first modality'
'attend only to the first modality and choose the strongest direction'
'respond to the stimulus displayed with greatest strength in the first modality'
'attend only to the first modality and pick the orientation displayed with maximal strength'
'choose the stimulus which appears with highest intensity in the first modality'
'select the most intensely displayed direction in the first modality'
'respond to the direction in the first modality that is strongest'
'attend only to the first modality and opt for the direction with highest intensity'
'focus on the first modality and choose the orientation which has greatest strength'
'select the direction in the first modality displayed most intensely'

*Validation*

'concentrate only on the first modality and choose the strongest direction'
'focus only on the first modality and opt for the direction displayed with the greatest strength'
'attend exclusively to stimuli in the first modality and choose the strongest one'
'only consider the first modality and choose the strongest direction'
'consider only the stimuli in the first modality and opt for the stimulus presented with highest strength'

**DMMod2:**

*Train*

'choose the direction in the second modality that appears strongest'
'attend only to the second modality and respond to the most intense orientation'
'focus on the second modality and select the direction displayed with greatest strength'
'pick the stimulus in the second modality that is displayed with maximal intensity'
'opt for the direction displayed in the second modality with most intensity'
'select the stimulus that has maximal strength in the second modality'
'focus only on the second modality and choose the strongest direction that appears there'
'respond to the direction which appears with highest intensity in the second modality'
'attend only to the second modality and pick the orientation that appears strongest'
'attend only to the second modality and opt for the stimulus displayed there with greatest strength'
'select the stimulus in the second modality which has maximal strength'
'pick the direction that is displayed most intensely in the second modality'
'focus on the second modality and respond to the orientation displayed there with highest intensity'
'choose the stimulus in the second modality which has maximal strength'
'pick the direction that is most intense in the second modality'

*Validation*

'concentrate on the stimuli from the second modality only and select the strongest direction'
'focus exclusively on the second modality and respond to the orientation presented with greatest strength'
'only consider stimuli presented in the second modality and choose the strongest among them'
'attend to the second modality and opt for the direction displayed with the most strength'
'consider only the stimuli that are displayed in the second modality and select the one with the highest strength'

**AntiDMMod1:**

*Train*

'attend to the first modality and choose the weakest direction'
'attend to the first modality and respond in the direction of minimal intensity'
'select the orientation in the first modality which has lowest strength'
'pick the stimulus in the first modality that is presented with least intensity'
'focus only on the first modality and choose the direction that appears weakest'
'respond to the stimulus in the first modality that has lowest strength'
'choose the direction in the first modality which is weakest'
'select the stimulus in the first modality that has minimal strength'
'attend to the first modality and select the direction that is displayed with least strength'
'opt for the direction in the first modality that appears weakest'
'focus only on the first modality and choose the orientation displayed with minimal intensity'
'select the stimulus in the first modality that has lowest intensity'
'pick the direction which appears with minimal intensity in the first modality'
'opt for the stimulus that appears weakest in the first modality'
'focus on the first modality and choose the stimulus minimal strength'

*Validation*

'concentrate on stimuli from the first modality only and opt for the weakest direction'
'concentrate on only stimuli in the first modality and select the weakest of the displayed directions'

'consider the stimuli in the first modality only and choose the direction with least strength'
'focus entirely on the stimuli appearing in the first modality and respond to the weakest direction presented there'
'concentrate on only the stimuli in the first modality and respond to the weakest direction'

**AntiDMMod2:**

*Train*

'attend to the second modality and choose the orientation that appears weakest'
'focus on the second modality and respond to the least intense direction'
'choose the direction with least intensity in the second modality'
'attend to the second modality and select the direction with least strength'
'choose the least intense orientation in the second modality'
'go in the direction of the weakest stimulus in the second modality'
'respond to the orientation which has lowest strength in the second modality'
'focus only on the second modality and pick the stimulus with lowest intensity'
'attend to the second modality and opt for the weakest stimulus presented there'
'pick the direction in the second modality presented with least strength'
'focus on the second modality and select the weakest direction'
'select the direction with lowest strength in the second modality'
'choose the direction with weakest intensity in the second modality'
'attend to the second modality and choose the orientation with lowest intensity'
'attend only to the second modality and select the direction with lowest strength'

*Validation*

'concentrate on stimuli in the second modality only and opt for the direction with least strength'
'concentrate on the second modality exclusively select the orientation that appears weakest'
'only consider the stimuli presented in the second modality and respond to the one presented with least strength'
'consider only the stimuli presented in the second modality and choose the weakest one that appears there'
'focus entirely on the stimuli displayed in the second modality and opt for the weakest direction'

**Dur1:**

*Train*

'respond to the first direction if it lasts for longer than the final direction otherwise do not respond'
'if the first stimulus is presented for a greater period of time than the latter stimulus then respond to the first stimulus otherwise do not respond'
'select the initial orientation if it has a duration which is greater than the second orientation otherwise do not respond'
'opt for the initial stimulus if it is displayed for more time than the final stimulus otherwise do not respond'
'choose the first orientation if it has a greater duration than the latter orientation otherwise do not respond'
'if the first stimulus is presented for longer than the second stimulus respond to the first stimulus otherwise do not respond'
'if the initial orientation is displayed for more time than the final orientation then choose the first orientation otherwise do not respond'
'go in the first direction if it appears for a greater amount of time than the latter stimulus otherwise do not respond'
'select the initial direction if it appears for longer than the second direction otherwise do not respond'
'if the first direction is displayed for more time than the final stimulus respond to the first direction otherwise do not respond'
'choose the initial orientation if it has a greater duration than the latter orientation otherwise do not respond'
'if the duration of the first orientation is greater than the second orientation respond to the first orientation otherwise do not respond'
'if the duration of the initial stimulus is greater than the duration of the final stimulus respond to the first stimulus otherwise do not respond'
'pick the first direction if it lasts for more time than the latter direction otherwise do not respond'
'pick the initial stimulus if it lasts for a greater period of time than the second stimulus otherwise do not respond'

*Validation*

'select the initial stimulus if it is presented for a greater span of time than the second direction otherwise do not respond'
'if the first orientation is displayed for a greater length of time than the second direction respond to the first orientation otherwise do not respond'
'choose the initial direction if is displayed for a longer span of time than the second direction otherwise do not respond'
'opt for the first stimulus if it lasts for a greater length of time than the second stimulus otherwise do not respond'
'if the initial orientation lasts for a larger span of time than the second orientation then select the initial orientation otherwise do not respond'

**Dur2:**

*Train*

'select the second direction if it lasts for longer than the first direction otherwise do not respond'
'if the duration of the final orientation is greater than the duration of the first orientation respond to the first orientation otherwise do not respond'
'if the duration of the later stimulus is greater than the duration of the first stimulus respond to the first stimulus otherwise do not respond'
'pick the second direction if it has a duration that is greater than the first direction otherwise do not respond'
'choose the final direction if it is displayed for more time than the first direction otherwise do not respond'
'opt for the latter orientation if it lasts for longer than the first orientation otherwise do not respond'
'if the final stimulus is presented for a longer time period than the first stimulus then respond to the final stimulus otherwise do not respond'
'if the second direction is displayed for more time than the first direction respond to the second direction otherwise do not respond'
'respond to the final orientation if is displayed for more time than the first orientation otherwise do not respond'
'select the second stimulus if it is longer than the first stimulus otherwise do not respond'
'if the duration of the latter direction is greater than the duration of the first direction respond to the latter direction otherwise do not respond'

'choose the second direction if it is presented for longer than the first direction otherwise do not respond'
'if the latter stimulus appears for more time than the first stimulus than select the second stimulus otherwise do not respond'
'if the duration of the latter stimulus is greater than the duration of the first stimulus than respond to the latter stimulus otherwise do not respond'
'choose the second direction if it lasts for a greater period of time than the first direction otherwise do not respond'

*Validation*

'respond to the final direction if the stimulus is presented for a greater span of time than the first stimulus otherwise do not respond'
'if the second direction lasts for a greater length of time than the first direction then select the second direction otherwise do not respond'
'select the second orientation if it appears for a longer span of time than the first direction otherwise do not respond'
'opt for the final stimulus if it is displayed for a span of time which is greater than the initial stimulus otherwise do not respond'
'if the final stimulus appears for a length of time which is greater than the first stimulus then choose the final stimulus otherwise do not respond'

**MultiDur1:**

*Train*

'respond to the first direction if it has longer duration averaged over both modalities than the second direction otherwise do not respond'
'if the first direction lasts for longer when combined over both modalities than the final direction then go in that direction otherwise do not respond'
'opt for the initial stimulus if it has a greater duration averaged both modalities than the latter stimulus otherwise do not respond'
'if the initial orientation appears for longer than the second orientation when considered across both modalities than select that orientation otherwise do not respond'
'pick the initial direction if it last for a longer period of time than the final direction in both modalities otherwise do not respond'
'if the first direction appears for a longer period of time then the latter direction over both modalities then select that direction otherwise do not respond'
'choose the initial stimulus if it has a greater duration than the final stimulus when considered over both modalities otherwise do not respond'
'respond to the initial direction if it is displayed for a greater period of time than the second direction averaged over both modalities otherwise do not respond'
'if the initial orientation lasts for longer than the second orientation when considered over both modalities then select that direction otherwise do not respond'
'choose the first displayed direction if it has a duration which is longer than the final direction combined over both modalities otherwise do not respond'
'if the initial stimulus appears for a longer period of time than the second stimulus averaged across modalities then choose that direction otherwise do not respond'
'select the first stimulus if it has a duration which is longer than the latter direction combined across both modalities otherwise do not respond'
'pick the initial orientation if if lasts for longer than the second direction averaged over modalities otherwise do not respond'
'if the first direction lasts for a greater period of time than the final direction in both modalities then respond to the second direction otherwise do not respond'
'select the initial orientation if it has a greater averaged duration over both modalities than the later orientation otherwise do not respond'

*Validation*

'respond to the initial direction if the length of the stimulus integrated over both modalities is greater than the second direction otherwise do not respond'
'choose the first stimulus if it spans a length of time which is greater than the second stimulus when summed over both modalities otherwise do not respond'
'if the first direction is displayed to a longer length of time summed over both modalities than the second direction then select the first direction otherwise do not respond'
'select the initial orientation if it has a longer time span when integrated over both modalities than the final direction otherwise do not respond'
'opt for the initial stimulus it it is displayed for a span of time which is longer than the span of the second stimulus summed over both modalities otherwise do not respond'

**MultiDur2:**

*Train*

'respond to the second direction if it lasts for a longer period of time over both modalities than the first direction otherwise do not respond'
'choose the final direction if it has a greater duration than the first direction averaged across modalities otherwise do not respond'
'if the latter orientation appears for a longer period of time when combined over both modalities than the initial direction choose the latter direction otherwise do not respond'
'if the second stimulus has a greater duration than the first stimulus combined over modalities than select the second stimulus otherwise do not respond'
'choose the latter direction if it is longer than the first direction averaged over both modalities otherwise do not respond'
'if the duration of the final stimulus is longer than the initial stimulus combined across modalities then choose the final stimulus otherwise do not respond'
'select the final orientation if it appears for longer on averaged over modalities than the first orientation otherwise do not respond'
'opt for the second direction if it lasts for a greater period of time than the first direction averaged over both modalities otherwise do not respond'
'if the time period of the final orientation lasts for longer than the first orientation when averaged over both modalities then respond to the final orientation otherwise do not respond'
'pick the second direction if it appears for a greater duration than the first direction when considered over both modalities otherwise do not respond'
'if the duration of the latter direction averaged across both modalities is greater than the first direction then choose the latter direction otherwise do not respond'
'if the final stimulus lasts for longer than the first stimulus when combined over both modalities then select the final stimulus otherwise do not respond'
'respond to the final direction if lasts for a longer period of time than the first stimulus averaged over both modalities otherwise do not respond'
'if the second stimulus has a duration which lasts for longer than the initial stimulus when combined across both modalities then

select the first stimulus otherwise do not respond'
'select the latter direction if it is displayed for a greater period of time than the initial direction when averaged over both modalities otherwise do not respond'

*Validation*

'respond to the final orientation if it spans a greater period of time when summed over both modalities than the initial orientation otherwise do not respond'
'select the last direction if it is presented for a greater length of time than the initial direction when integrated over both modalities otherwise do not respond'
'opt for the final stimulus if it appears for a greater span of time than the initial stimulus when summed over both modalities otherwise do not respond'
'if the second direction is displayed for a length of time which is greater than the first direction when summed over both modalities then respond to the second direction otherwise do not respond'
'pick the final orientation if it appears for a span of time which is larger than the first orientation when integrated over both modalities otherwise do not respond'

**AntiDur1:**

*Train*

'respond to the first direction if it has a shorter duration than the second direction otherwise do not respond'
'choose the initial orientation if it lasts for a shorter period of time than the final orientation otherwise do not respond'
'if the first stimulus has a shorter duration than the latter stimulus then select the first stimulus otherwise do not respond'
'if the initial direction is lasts for less time than the final direction choose the initial direction otherwise do not respond'
'opt for the first orientation if it lasts for less time than the second orientation otherwise do not respond'
'select the initial stimulus if it has a shorter duration than the final stimulus otherwise do not respond'
'pick the first direction if it lasts for less time than the final direction otherwise do not respond'
'go in first direction if it has a shorter duration than the latter direction otherwise do not respond'
'respond to the first orientation if it is shorter than the final orientation otherwise do not respond'
'if the initial direction is displayed for less time than the latter direction select the initial direction otherwise do not respond'
'if the first orientation appears for less time than the second direction choose the first orientation otherwise do not respond'
'if the initial stimulus is shorter than the final stimulus respond to the initial stimulus otherwise do not respond'
'respond to the initial orientation if it is shorter than the second orientation otherwise do not respond'
'select the first direction if it has a shorter duration than the final direction otherwise do not respond'
'pick the initial stimulus if it is displayed for less time than the second stimulus otherwise do not respond'

*Validation*

'select the initial stimulus if it spans a length of time which is shorter than latter stimulus otherwise do not respond'
'opt for the initial stimulus if it appears for a length of time which is less than the latter stimulus otherwise do not respond'
'if the first direction is presented for an amount of time which is less than the latter direction than choose the earlier direction otherwise do not respond'
'choose the first orientation if it appears for a shorter span of time than the second stimulus otherwise do not respond'
'pick the initial orientation if it is displayed for a length of time that is less than the final direction otherwise do not respond'

**AntiDur2:**

*Train*

'respond to the final direction if it has a shorter duration than the first direction otherwise do not respond'
'choose the latter stimulus if it is displayed for less time than the first stimulus otherwise do not respond'
'if the second orientation appears for less time than the initial orientation select the second orientation otherwise do not respond'
'if the latter direction is shorter than the initial direction pick the latter direction otherwise do not respond'
'opt for the final direction if it has a shorter duration than the first direction otherwise do not respond'
'respond to the second stimulus if it is displayed for less time than the first stimulus otherwise do not respond'
'if the final stimulus is shorter than the initial stimulus choose the final stimulus otherwise do not respond'
'if the second direction has a shorter duration than the initial direction select the second direction otherwise do not respond'
'if the second orientation is shorter than the first orientation pick the second orientation otherwise do not respond'
'select the latter stimulus if it has a shorter duration than the initial stimulus otherwise do not respond'
'opt for the final orientation if it is displayed for less time than the initial orientation otherwise do not respond'
'if the latter direction appears for less time than the first direction respond to the latter direction otherwise do not respond'
'choose the final orientation if it is shorter than the initial orientation otherwise do not respond'
'select the second stimulus if it has a shorter duration than the first stimulus otherwise do not respond'
'respond to the latter direction if it is shorter than the first direction otherwise do not respond'

*Validation*

'opt for the latter stimulus if is presented for a span of time which is less than the earlier stimulus otherwise do not respond'
'pick the last direction if it appears for a length of time that is less than the earlier direction otherwise do not respond'
'if the latter orientation is presented for a shorter span of time than the first orientation select the latter orientation otherwise do not respond'
'select the latter stimulus if it was displayed for a span of time which was shorter than the first stimulus otherwise do not respond'
'choose the latter stimulus if it appears for a length of that is was shorter than the first stimulus otherwise do not respond'

**AntiMultiDur1:**

*Train*

'respond to the initial direction if it is displayed for a shorter amount of time than the second direction averaged over modalities otherwise do not respond'
'if the first stimulus is shorter than the final stimulus combined over both modalities then choose the first stimulus otherwise do not respond'
'if the initial orientation has a shorter duration than the latter orientation considered over both modalities then respond to the initial orientation otherwise do not respond'
'choose the first stimulus if it is shorter than the final stimulus

averaged over both modalities otherwise do not respond'
'select the initial direction if it is displayed for less time than the latter direction combined over both modalities otherwise do not respond'
'if the first direction is displayed for less time than the final direction averaged over both modalities then opt for the first direction otherwise do not respond'
'if the initial stimulus has a shorter duration than the second stimulus considered over both modalities then pick the initial stimulus otherwise do not respond'
'opt for the first orientation if it is shorter than the second orientation averaged over both modalities otherwise do not respond'
'pick the first stimulus if it appears for less time than the final stimulus combined over both modalities otherwise do not respond'
'respond to the initial orientation if it is displayed for less time averaged over both modalities than the second orientation otherwise do not respond'
'if the first direction is shorter on averaged over both modalities than the final direction then respond to the first direction otherwise do not respond'
'if the initial stimulus has a shorter duration when combined over both modalities than the latter stimulus pick the initial stimulus otherwise do not respond'
'pick the first stimulus if it appears for less time considered over both modalities than the second stimulus otherwise do not respond'
'if the initial orientation appears for less time averaged over both modalities than the latter stimulus choose the initial orientation otherwise do not respond'
'go in the first direction if it is shorter combined over both modalities than the second direction otherwise do not respond'

*Validation*

'pick the initial direction if its lasts for a span of time which is less than the latter direction when summed over both modalities otherwise do not respond'
'choose the first orientation if it is presented for a span of time which is less than the second direction when integrated over both modalities otherwise do not respond'
'if the initial stimulus appears for a length of time which is shorter when summed over both modalities than the second stimulus respond to the earlier stimulus otherwise do not respond'
'select the initial direction if it is displayed for a length of time which is shorter than the second direction when summed over both modalities otherwise do not respond'
'select the first stimulus if it is displayed for a length of time which is less than that of the second stimulus integrated over both modalities otherwise do not respond'

**AntiMultiDur2:**

*Train*

'select the final direction if it is shorter than the first direction averaged over both modalities otherwise do not respond'
'respond to the second orientation if is displayed for less time than the initial orientation considered across both modalities otherwise do not respond'
'if the latter stimulus is displayed for less time combined over both modalities than the initial stimulus then select the latter stimulus otherwise do not respond'
'if the final stimulus is shorter than the first stimulus averaged over both modalities than select the final stimulus otherwise do not respond'
'choose the latter orientation if it has a shorter duration averaged over both modalities than the initial orientation otherwise do not respond'
'if the second direction has a shorter duration than the first direction considered across both modalities than select the second direction otherwise do not respond'
'if the final orientation is shorter than the initial orientation combined across both modalities then choose the final orientation otherwise do not respond'
'opt for the latter stimulus it it is presented for less time than the initial stimulus considered across both modalities otherwise do not respond'
'pick the final direction if it has a shorter duration averaged over both modalities than the first direction otherwise do not respond'
'go in the second direction if it is shorter when combined over both modalities than the first direction otherwise do not respond'
'respond to the latter direction if it is displayed for less time than the initial direction combined over both modalities otherwise do not respond'
'if the second stimulus is shorter than the first stimulus combined over both modalities then opt for the second stimulus otherwise do not respond'
'if the final direction is displayed for less time combined over both modalities than the first direction then choose the final direction otherwise do not respond'
'select the second orientation if it has a shorter duration considered across both modalities than the initial orientation otherwise do not respond'
'pick the final direction if it is shorter than the first direction averaged over both modalities otherwise do not respond'

*Validation*

'pick the latter orientation if it appears for an amount of time which is less than the first orientation when summed across modalities otherwise do not respond'
'opt for the latter direction if it is presented for a span of time which is less than the earlier orientation when integrated over both modalities otherwise do not respond'
'respond to the latter stimulus if it appears for a length of time that is shorter than the earlier stimulus integrated across modalities otherwise do not respond'
'if the final direction is presented for a time span that is less than the initial direction when summed over both modalities then choose the final direction otherwise do not respond'
'if the latter stimulus appears for the shorter time span than the earlier stimulus when combined over both modalities then opt for the latter stimulus otherwise do not respond'

**Dur1Mod1:**

*Train*

'attend to the first modality and choose the first direction if has a longer duration than the second direction otherwise do not respond'
'attend only to the first modality and select the initial stimulus if it lasts for a longer period of time than the latter stimulus otherwise do not respond'
'if the first orientation lasts for longer than the final orientation in the first modality then respond to the first orientation otherwise do not respond'
'focus only on the first modality and opt for the initial direction if it has a greater duration than the second direction otherwise do not respond'
'pay attention only to the first modality and if the first stimulus lasts for more time than the final stimulus then respond to the first stimulus otherwise do not respond'
'select the first direction if it is presented for a longer duration

than the latter direction in the first modality otherwise do not respond'
'if the initial orientation is displayed for more time than the second orientation in the first modality then respond to the initial orientation otherwise do not respond'
'focus only on the first modality and opt for the first stimulus if it is presented for longer than the final stimulus otherwise do not respond'
'focus on the first modality and pick the first orientation if it appears for a greater period of time than the latter orientation otherwise do not respond'
'go in the first direction if the is has greater duration than the second stimulus in the first modality otherwise do not respond'
'attend only to the first modality and choose the first direction if is appears for more time than the final direction otherwise do not respond'
'pay attention to the first modality and respond to the initial orientation if it has a duration that is greater than the latter stimulus otherwise do not respond'
'pay attention only to the first modality and select the initial stimulus if is lasts for longer than the second stimulus otherwise do not respond'
'attend only to the first modality and pick the initial orientation if it has a longer duration than the final orientation otherwise do not respond'
'focus on the first modality and opt for the first direction if it lasts for longer than the second direction otherwise do not respond'

*Validation*

'concentrate only on the first modality and select the earlier direction if it lasts for a longer time span than the latter direction otherwise do not respond'
'focus exclusively on the first modality and choose the earlier stimulus if it is presented for a greater length of time than the second stimulus otherwise do not respond'
'consider only stimuli from the first modality and pick the first stimulus if it appears for a greater period of time than the second stimulus otherwise do not respond'
'concentrate exclusively on stimuli in the first modality and choose the first direction if it appears for a greater span of time than the second direction otherwise do not respond'
'only consider the first modality and opt for the earlier stimulus if it appears for a length of time which is greater than the latter stimulus otherwise do not respond'

**Dur1Mod2:**

*Train*

'attend only to the second modality and select the initial direction if it lasts for longer than the final direction otherwise do not respond'
'if the first orientation is displayed for more time than the final direction in the second modality then respond to the first orientation presented there otherwise do not respond'
'attend only to the second modality and pick the first stimulus if it lasts for a greater period of time than the latter direction otherwise do not respond'
'pay attention only to the second modality and opt for the initial direction if it has a longer duration than the final direction otherwise do not respond'
'focus on the second modality and select the first stimulus if it appears for a greater amount of time than the second stimulus otherwise to not respond'
'focus on the second modality and choose the initial orientation if it is presented for longer than the final orientation otherwise do not respond'
'go in the first direction if it has a greater duration than the latter direction in the second modality otherwise do not respond'
'select the initial stimulus if it has a greater duration than the final stimulus in the second modality otherwise do not respond'
'if the duration of the first stimulus is longer than the duration of the latter stimulus in the second modality respond to the first stimulus otherwise do not respond'
'if the first direction lasts for a greater period of time than the final direction in the second modality then select the first direction otherwise do not respond'
'pay attention to the second modality and pick the first orientation if it appears for a greater period of time than the second orientation otherwise do not respond'
'pay attention only to the second modality and select the initial direction if it lasts for longer than the second direction otherwise do not respond'
'attend to the second modality and if the duration of the initial stimulus is greater than that of the final stimulus then respond to the initial direction otherwise do not respond'
'focus on the second modality and choose the first orientation if it lasts for more time than the second orientation otherwise do not respond'
'attend only to the second modality and opt for the initial direction if it lasts for longer than the latter direction otherwise do not respond'

*Validation*

'concentrate only on the second modality and respond to the first direction if spans a greater length of time than the second stimulus otherwise do not respond'
'focus exclusively on the stimuli in the second modality and respond to the initial direction if it is displayed for a greater length of time than the latter stimulus otherwise do not respond'
'only consider stimuli which appear in the second modality and select the first orientation if it appears for a greater span of time than the second orientation otherwise do not respond'
'concentrate only on stimuli which appear in the second modality and choose the initial direction if it lasts for a greater period of time than the second direction otherwise do not respond'
'attend exclusively to the second modality and respond to the first stimulus if it is presented for a greater span of time than the final stimulus otherwise do not respond')

**Dur2Mod1:**

*Train*

'attend only to the first modality and select the second direction if it lasts for a greater period of time than the first direction otherwise do not respond'
'pay attention only to the first modality and if the final orientation has a longer duration than the initial direction respond to the first orientation otherwise do not respond'
'focus only on the first modality and choose the latter direction if it lasts for more time than the first direction otherwise do not respond'
'select the second direction if it is displayed for a greater period of time than the first direction in the first modality otherwise do not respond'
'focus only on the first modality and pick the final orientation if it has a longer duration than the initial direction otherwise do not respond'
'pay attention only to the first modality and if the second stimulus is displayed for more time than the first stimulus respond to the second stimulus otherwise do not respond'

'if the duration of the final stimulus is longer than the first stimulus in the first modality then respond to the final stimulus otherwise do not respond'
'if the latter direction lasts for a greater period of time than the initial direction in the first modality then select the latter direction otherwise do not respond'
'attend only to the first modality and opt for the second stimulus if it appears for a greater period of time than the initial stimulus otherwise do not respond'
'choose the second orientation if it has a greater duration than the initial orientation in the first modality otherwise do not respond'
'focus only on the first modality and choose the second direction if it is longer than the initial direction otherwise do not respond'
'attend to the first modality and opt for the final orientation if it lasts for longer than the first orientation otherwise do not respond'
'pay attention only to the first modality and choose the latter stimulus if it is displayed for a longer period of time than the first stimulus otherwise do not respond'
'respond to the second stimulus if it appears for a longer period of time than the first stimulus in the first modality otherwise do not respond'
'attend to the first modality and respond to the latter orientation if it lasts for a greater period of time than the first orientation otherwise do not respond'

*Validation*

'concentrate only on the first modality and select the latter orientation if it spans a longer length of time than the earlier orientation otherwise do not respond'
'attend exclusively to the first modality and choose the latter stimulus if it last for longer than the initial stimulus otherwise do not respond'
'only consider stimuli in the first modality and opt for the second stimulus if it spans a period of time which is greater than the first stimulus otherwise do not respond'
'focus only on the first modality and select the latter direction if it lasts for a greater length of time than the earlier direction otherwise do not respond'
'concentrate on only the stimuli in the first modality and pick the final stimulus if it spans a length of time which is greater than the first stimulus otherwise do not respond'

**Dur2Mod2:**

*Train*

'attend only to the second modality and respond to the final direction if it has a longer duration than the first direction otherwise do not respond'
'focus only on the second modality and choose the latter stimulus if it appears for a greater period of time than the first direction otherwise do not respond'
'if the second orientation appears for a greater period of time than the initial orientation in the second modality then select the second orientation otherwise do not respond'
'if the duration of the second stimulus is longer than that of the first stimulus in the second modality then respond to the second stimulus otherwise do not respond'
'focus on the second modality and choose the final orientation if it lasts for longer than the first orientation otherwise do not respond'
'attend to the second modality and select the latter stimulus if it is displayed for a greater period of time than the first stimulus otherwise do not respond'
'choose the second direction if it has a greater duration than the first direction in the second modality otherwise do not respond'
'respond to the second orientation if it lasts for longer than the initial orientation in the second modality otherwise do not respond'
'if the duration of the final direction is longer than the duration of the first direction in the second modality then respond to the final direction otherwise do not respond'
'pay attention only to the second modality and if the final stimulus lasts for more time than the first stimulus then respond to the second stimulus otherwise do not respond'
'pay attention to the second modality and pick the latter direction if it is displayed for more time than the first direction otherwise do not respond'
'choose the second orientation if it has a longer duration than the first orientation in the second modality otherwise do not respond'
'attend only to the second modality and opt for the final stimulus if it lasts for a greater period of time than the first stimulus otherwise do not respond'
'attend only to the second modality and select the latter orientation if it has a longer duration than the first orientation otherwise do not respond'
'pick the second direction if it appears for more time than the first direction in the second modality otherwise do not respond'

*Validation*

'focus exclusively on stimuli in the second modality and choose the final direction if it spans a period of time which is longer than the earlier direction otherwise do not respond'
'concentrate only on the second modality and opt for the last direction if it appears for a length of time that is greater than the earlier direction otherwise do not respond'
'only consider stimuli in the second modality and choose the final orientation if it is presented for a span of time which lasts longer than the initial orientation otherwise do not respond'
'concentrate only on stimuli in the second modality and select the latter stimulus if it lasts for a greater length of time than the earlier stimulus otherwise do not respond'
'attend exclusively to the second modality and opt for the final direction if it is presented for a length of time which is longer than that of the initial direction otherwise do not respond'

**COMP1:**

*Train*

'if the first stimulus is greater than the second stimulus respond to the first stimulus otherwise do not respond'
'if the first stimulus has higher intensity than the final go in the first direction otherwise do not respond'
'go in the direction of the first stimulus if it is stronger than the second stimulus otherwise do not respond'
'when the initial stimulus has higher value than the final stimulus respond in the initial direction otherwise do not respond'
'if the initial stimulus has greater strength than the latter stimulus respond in the first direction otherwise do not respond'
'respond to the initial stimulus if it is stronger than the last stimulus otherwise do not respond'
'choose the initial stimulus when it is presented with higher intensity relative to the second stimulus otherwise do not respond'
'if the initial stimulus has higher value than the second respond to the first direction otherwise do not respond'
'when the first stimulus is presented with the higher intensity than the latter select the first direction otherwise do not respond'
'choose the first stimulus if it is presented with the greater intensity than the second otherwise do not respond'
'select the initial stimulus if is stronger than the final stimulus otherwise do not respond'

‘if the first stimulus has larger value than than the latter stimulus select the first direction otherwise do not respond’
‘respond to the first direction if it is more intense than the last stimulus otherwise do not respond’
‘if the first stimulus is more intense than the second stimulus choose the first direction otherwise do not respond’
‘when the initial stimulus is presented with greater strength than the second stimulus respond to the first direction otherwise do not respond’

*Validation*

‘when the first stimulus is stronger pick the first orientation otherwise do not respond’
‘choose the initial stimulus if it is the stronger of the two presented stimuli otherwise do not respond’
‘if the initial orientation has higher value than the subsequent select the first orientation otherwise do not respond’
‘pick the first direction if the stimulus presented has the greater value otherwise do not respond’
‘when the first stimulus has greater intensity than the second then respond to the first stimulus otherwise do not respond’

**COMP2:**

*Train*

‘respond in the direction of the second stimulus if it has greater intensity than the first otherwise do not respond’
‘if the final stimulus is presented with higher value than the initial stimulus go in the final direction otherwise do not respond’
‘when the final stimulus has greater strength than the first select the final orientation otherwise do not respond’
‘respond in the direction of the second stimulus if it has higher intensity than the initial stimulus otherwise do not respond’
‘if the final stimulus has higher value than the first stimulus choose the final orientation otherwise do not respond’
‘select the latter stimulus direction if it is presented with greater strength than the first otherwise do not respond’
‘when the last stimulus is presented with more intensity than the first respond in the second direction otherwise do not respond’
‘choose the final stimulus if it has the greater value than the first otherwise do not respond’
‘if the final stimulus is presented with the higher intensity than the initial stimulus respond to the second stimulus otherwise do not respond’
‘select the final stimulus if it is presented with the higher intensity than the initial stimulus otherwise do not respond’
‘when the second stimulus is more intense than the initial stimulus respond to the second stimulus otherwise do not respond’
‘if the second stimulus is presented with greater strength than the first respond to the second stimulus otherwise do not respond’
‘choose the latter stimulus if it is more intense than the the first stimulus otherwise do not respond’
‘when the final stimulus is more intense than the first respond to the final stimulus otherwise do not respond’
‘pick the second stimulus if it has more intensity than the first otherwise do not respond’

*Validation*

‘if the last stimuli is presented with higher intensity than the second respond in second direction otherwise do not respond’
‘pick the direction of the second stimuli if it has greatest value than the first otherwise do not respond’
‘when the final direction has greater intensity then the first go in that direction otherwise do not respond’
‘choose the subsequent stimulus direction if it has greater strength than the first otherwise do not respond’
‘if the second orientation is presented with more intensity than the first respond with the first orientation otherwise do not respond’

**MultiCOMP1:**

*Train*

‘respond to the first direction when it has greater strength on average than the second stimulus otherwise do not respond’
’respond if the first direction has higher combined intensity over two modalities than the final stimulus otherwise do not respond’
’if the joint intensity of the first directions is higher than the second then respond to the first direction otherwise do not respond’
’choose the initial direction if the integrated strength across modalities is greater than the final direction otherwise do not respond’
’go in the initial direction if it has greater value than the final direction when combined over both modalities otherwise do not respond’
’select the first direction when it has greater integrated strength across modalities than the second direction otherwise do not respond’
’if the initial direction has higher overall value over modalities than the final direction then choose the initial direction otherwise do not respond’
’if the combined intensity of the first direction is higher than the latter direction then respond to the first direction otherwise do not respond’
’choose the initial direction when the joint strength over both modalities is greater than the final direction otherwise do not respond’
’pick the first direction if the average value is larger than the second direction for both modalities otherwise do not respond’
’select the initial direction if it displays a higher overall intensity over modalities than the final direction otherwise do not respond’
’if the intensity of the first stimulus integrated over modalities is greater than the second respond in the first direction otherwise do not respond’
’select the first stimulus if it has higher joint strength over modalities than the final otherwise do not respond’
’if the first direction is presented with higher intensity averaged over modalities than the second direction then respond to the first direction otherwise do not respond’
’pick the initial stimulus if it has greater joint intensity over both modalities than the latter otherwise do not respond’

*Validation*

‘if the initial direction displays higher averaged value than the second direction then respond to initial direction otherwise do not respond’
‘pick the first orientation if it has greater combined intensity than the second orientation otherwise do not respond’
‘when the first direction has higher intensity averaged over modalities than the second direction then select the first direction otherwise do not respond’
‘choose the initial orientation when it has greater joint value over modalities than the final direction otherwise do not respond’
‘if the first orientation has greater joint strength over both modalities than the second then respond with the first orientation otherwise do not respond’

**MultiCOMP2:**

*Train*

'respond in the second direction if it has larger overall strength over modalities than the first direction otherwise do not respond'
'if the final direction has greater combined value than the initial direction then respond in the final direction otherwise do not respond'
'respond in the latter direction when it has higher integrated value over modalities than the first direction otherwise do not respond'
'select the second direction if it has greater strength than the first direction over both modalities otherwise do not respond'
'choose the final direction if it has a joint intensity higher than the initial direction over modalities otherwise do not respond'
'if the final direction is presented with greater overall strength across modalities than the initial direction choose the final direction otherwise do not respond'
'respond to the latter direction when it has larger combined value over modalities than the first direction otherwise do not respond'
'select the final direction if the strength integrated over modalities is greater than the first direction otherwise do not respond'
'if the final direction is represented with higher joint intensity over both modalities than the initial direction then respond to the final direction otherwise do not respond'
'choose the second direction when its combined strength over modalities is higher than the first direction otherwise do not respond'
'if the second stimulus is presented with higher joint intensity over modalities than the first then select the second stimulus otherwise do not respond'
'pick the final direction if it has larger strength combined over modalities than the initial direction otherwise do not respond'
'respond to the second stimulus if it is presented with greater intensity averaged over modalities than the first stimulus otherwise do not respond'
'choose the latter direction if it has stronger presentation averaged over modalities than the first direction otherwise do not respond'
'if the final stimulus has higher overall value for both modalities than the initial direction then select the final stimulus otherwise do not respond'

*Validation*

'if the final orientation displays greater overall value than the initial orientations combined over modalities then select the final orientation otherwise do not respond'
'pick the second direction if it has a bigger average value over modalities than the first direction otherwise do not respond'
'choose the last direction if it has bigger combined intensity over modalities than the initial direction otherwise do not respond'
'select the second direction if it has a larger joint strength over modalities than the first direction otherwise do not respond'
'respond to the final orientation if it has larger combined intensity than the first orientation otherwise do not respond'

**AntiCOMP1:**

*Train*

'respond to the first direction if it is weaker than the final direction otherwise do not respond'
'if the initial stimulus is presented with less intensity than the latter stimulus then select the initial stimulus otherwise do not respond'
'choose the initial orientation if it has lower intensity than the second orientation otherwise do not respond'
'if the first direction is displayed less strength than the final direction then select the first direction otherwise do not respond'
'opt for the initial orientation if is weaker than the latter orientation otherwise do not respond'
'pick the initial stimulus if it is displayed with lower strength than the second stimulus otherwise do not respond'
'if the first direction is less intense than the final direction then choose the first direction otherwise do not respond'
'select the initial orientation if it is presented with lower strength than the latter orientation otherwise do not respond'
'if the first stimulus is less intense than the second stimulus respond to the first stimulus otherwise do not respond'
'go in the direction of the first stimulus if it has less intensity than the final stimulus otherwise do not respond'
'if the initial direction has less strength than the second direction then select the initial direction otherwise do not respond'
'if the first stimulus is presented with less strength than the final direction then choose the first stimulus otherwise do not respond'
'respond to the initial orientation if it is weaker than the second orientation otherwise do not respond'
'select the first stimulus if it is presented with less intensity than the second stimulus otherwise do not respond'
'choose the first direction if is has less strength than the latter direction otherwise do not respond'

*Validation*

'go in the initial direction if it is presented with decreased intensity compared to the final orientation otherwise do not respond'
'if the initial direction has decreased strength compared to the last direction then choose the initial direction otherwise do not respond'
'opt for the initial stimulus if it appears weaker than the latter stimulus otherwise do not respond'
'choose the initial orientation if it is presented with lower intensity compared to the latter stimulus otherwise do not respond'
'opt for the initial direction it appear with decreased intensity compared to the latter direction otherwise do not respond'

**AntiCOMP2:**

*Train*

'respond to the final direction if it is weaker than the initial direction otherwise do not respond'
'if the second orientation is presented with less strength than the first direction then select the second orientation otherwise do not respond'
'if the latter stimulus is less intense than the first stimulus then respond to the latter stimulus otherwise do not respond'
'choose the second orientation if it is displayed with lower intensity than the first orientation otherwise do not respond'
'select the final stimulus if it is weaker than the first direction otherwise do not respond'
'opt for the second direction if it has less strength than the initial direction otherwise do not respond'
'if the final orientation is presented with less intensity than the first orientation then select the final orientation otherwise do not respond'
'if the second stimulus has lower intensity than the first stimulus respond to the second stimulus otherwise do not respond'
'go in the final direction if it has less strength than the initial direction otherwise do not respond'
'if the second stimulus has lower intensity than the initial stimulus respond to the second stimulus otherwise do not respond'
'if the latter direction is less intense than the initial direction then select the latter direction otherwise do not respond'
'pick the second orientation if it is weaker than the initial orientation otherwise do not respond'
'respond to the final stimulus if it is weaker than the first stimulus otherwise do not respond'

'choose the second direction if it is presented with lower intensity than the initial direction otherwise do not respond'
'if the final stimulus is weaker than the initial stimulus respond to the final stimulus otherwise do not respond'

*Validation*

'opt for the latter orientation if it presented with lower strength than the earlier orientation otherwise do not respond'
'if the final stimulus appears with less intensity compared to the earlier stimulus then opt for the second stimulus otherwise do not respond'
'choose the last direction if it appears with less intensity compared to the initial direction otherwise do not respond'
'select the last orientation if it is displayed with less intensity compared to the initial direction otherwise do not respond'
'select the final direction if it is presented with less intensity compared to the initial direction otherwise do not respond'

**AntiMultiCOMP1:**

*Train*

'respond to the first direction if it is weaker averaged over both modalities than the final direction otherwise do not respond'
'if the first orientation has less strength over both modalities than the second orientation then select the first orientation otherwise do not respond'
'if the first stimulus is less intense than the latter stimulus when averaged over both modalities then choose the stimulus stimulus otherwise do not respond'
'pick the first direction if it has lower intensity than the second direction combined over both modalities otherwise do not respond'
'select the initial direction if it is weaker than the final direction averaged over both modalities otherwise do not respond'
'opt for the initial orientation if it has lower overall strength than the latter orientation considered over both modalities otherwise do not respond'
'if the initial stimulus is presented with less intensity than the second stimulus averaged over both modalities then select the initial stimulus otherwise do not respond'
'if the first direction is weaker averaged over both modalities than the final direction select the first direction otherwise do not respond'
'if the initial orientation has less intensity combined over both modalities than the latter orientation respond to the initial orientation otherwise do not respond'
'select the first direction if it has less overall strength over both modalities than the second direction otherwise do not respond'
'choose the initial orientation if it is presented with lower overall intensity over both modalities than the final orientation otherwise do not respond'
'if the first orientation has lower combined intensity over both modalities than the second orientation then select the first orientation otherwise do not respond'
'respond to the initial stimulus if it is presented with less combined strength over both modalities than the latter stimulus otherwise do not respond'
'if the first direction is presented lower intensity averaged over both modalities than the second direction then select the first direction otherwise do not respond'
'pick the initial stimulus if it is weaker averaged over both modalities than the final stimulus otherwise do not respond'

*Validation*

'opt for the first direction if it appears with decreased intensity compared to the last direction when summed over both modalities otherwise do not respond'
'select the initial orientation if it is shown with decreased strength compared to the last orientation integrated across modalities otherwise do not respond'
'if the first stimulus has lower strength when compared to the latter stimulus summed over both modalities then choose the latter stimulus otherwise do not respond'
'pick the initial stimulus if it is presented with decreased strength compared to the latter stimulus when integrated over both modalities otherwise do not respond'
'if the first direction is shown with lower strength compared to the strength of the latter direction summed over both modalities then select the first direction otherwise do not respond'

**AntiMultiCOMP2:**

*Train*

'respond to the second direction if it is weaker averaged over both modalities than the first direction otherwise do not respond'
'if the final orientation is presented with less intensity combined over both modalities than the first direction select the final orientation otherwise do not respond'
'if the second stimulus has lower strength averaged over both modalities than the initial stimulus then select the second stimulus otherwise do not respond'
'opt for the final direction if is less intense combined over both modalities than the initial direction otherwise do not respond'
'pick the second orientation if it has lower overall intensity over both modalities than the first orientation otherwise do not respond'
'if the final stimulus is weaker averaged over both modalities than the initial stimulus then select the final stimulus otherwise do not respond'
'if the latter direction is displayed with less intensity combined over both modalities than the first direction then select the latter direction otherwise do not respond'
'if the second orientation has less strength averaged over both modalities than the initial orientation then select the second orientation otherwise do not respond'
'choose the final stimulus if it has lower overall strength over both modalities than the first stimulus otherwise do not respond'
'select the latter direction if has less intensity when combined over both modalities than the initial direction otherwise do not respond'
'if the second orientation is weaker than the first orientation averaged over both modalities then choose the second orientation otherwise do not respond'
'opt for the final stimulus if it has lower intensity averaged over both modalities than the first stimulus otherwise do not respond'
'if the second direction is presented with weaker intensity combined over both modalities than the first stimulus select the second direction otherwise do not respond'
'respond to the final stimulus if it is weaker when combined over both modalities than the first stimulus otherwise do not respond'
'if the latter orientation is displayed with less intensity averaged over both modalities than the first orientation then choose the latter orientation otherwise do not respond'

*Validation*

'if the last orientation is shown with less strength than the earlier orientation when added across modalities then opt for the latter orientation otherwise do not respond'
'choose the last displayed direction if it is shown with lower intensity compared to the first direction when integrated over modalities otherwise do not respond'

'opt for the latter direction if it appears with less strength compared to the initial direction when stimuli strength are integrated over both modalities otherwise do not respond'
'select the last stimulus if it is displayed with a low strength compared to the first stimulus when intensity is summed over both modalities otherwise do not respond'
'pick the latter direction if it has less intensity compared to earlier direction when strength is integrated over both modalities otherwise do not respond'

**COMP1Mod1:**

*Train*

'attend only to the first modality and respond to the first direction if is stronger than the second stimulus otherwise do not respond'
'if the first stimulus has greater strength than the final stimulus in the first modality select the first direction otherwise do not respond'
'pay attention only to the first modality and select the first orientation if it is presented with greater intensity than the latter orientation otherwise do not respond'
'focus only on the first modality and choose the initial stimulus if it has more intensity than the second stimulus otherwise do not respond'
'if the initial direction is stronger than the latter direction in the first modality then respond to the initial direction otherwise do not respond'
'choose the first direction if it is presented with higher intensity than the final direction in the first modality otherwise do not respond'
'if the first stimulus has more intensity than the latter stimulus in the first modality then choose the first stimulus otherwise do not respond'
'attend only to the first modality and select the initial direction if if has more strength than the final stimulus otherwise do not respond'
'if the first orientation is presented with higher intensity than the second orientation in the first modality then respond to the first orientation otherwise do not respond'
'pick the first direction if it is stronger than the second direction in the first modality otherwise do not respond'
'respond to the initial direction if it is presented with more strength than the final direction in the first modality otherwise do not respond'
'opt for the first orientation if it is more intense than the second direction in the first modality otherwise do not respond'
'focus only on the first modality and select the initial stimulus if it has greater strength than the final direction otherwise do not respond'
'if the first orientation has higher intensity than the latter orientation in the first modality then respond to the first orientation otherwise do not respond'
'pay attention only to the first modality and opt for the first direction if it has more strength than the second direction otherwise do not respond'

*Validation*

'concentrate on stimuli from the first modality only and opt for the first direction if it is presented with larger intensity compared to the latter direction otherwise do not respond'
'only consider the stimuli that are displayed in the first modality and select the initial stimulus if is appears with a greater contrast when compared to the final stimulus otherwise do not respond'
'only concentrate on the orientations presented in the first modality and if the initial direction is presented with higher contrast than the latter direction opt for the earlier direction otherwise do not respond'
'exclusively attend to stimuli from the first modality and respond to the earlier orientation if it is displayed with higher intensity compared to the last orientation otherwise do not respond'
'concentrate only on the first modality and pick the first direction if it is displayed with higher contrast compared to the latter direction otherwise do not respond'

**COMP1Mod2:**

*Train*

'focus only on the second modality and select the first direction if it is stronger than the second direction otherwise do not respond'
'if the initial orientation has higher intensity than the final orientation in the second modality then select the initial orientation otherwise do not respond'
'pay attention only to the second modality and choose the initial stimulus if it has greater strength than the final stimulus otherwise do not respond'
'select the first direction if it has more intensity than the latter direction in the second modality otherwise do not respond'
'if the first stimulus is stronger than the final stimulus in the second modality then select the first stimulus otherwise do not respond'
'attend only to the second modality and if the first stimulus is greater than the second stimulus respond to the first stimulus otherwise do not respond'
'choose the first direction if it is presented with greater strength than the final direction in the second modality otherwise do not respond'
'if the initial direction has greater strength than the latter direction in the second modality then respond to the initial direction otherwise do not respond'
'focus only on the second modality and opt for the first orientation if it is presented with more intensity than the second orientation otherwise do not respond'
'select the initial direction if it is displayed with greater strength than the final direction in the second modality otherwise do not respond'
'pick the first stimulus if it is stronger than the latter stimulus in the second modality otherwise do not respond'
'attend only to the second modality and select the first orientation if it has greater strength than the second orientation otherwise do not respond'
'choose the first direction if it has higher intensity than the final direction in the second modality otherwise do not respond'
'if the initial stimulus is more intense than the last stimulus in the second modality then respond to the initial stimulus otherwise do not respond'
'opt for the first direction if it is stronger than the second direction in the second modality otherwise do not respond'

*Validation*

'only consider stimuli from the second modality and select the initial direction if it appears with more contrast than the latter direction otherwise do not respond'
'concentrate exclusively on the second modality and choose the first orientation if it is displayed with a more intensity compared to last orientation otherwise do not respond'
'focus entirely on the second modality and if the initial stimulus is displayed with higher contrast than the last stimulus respond to the first stimulus otherwise do not respond'
'consider only stimuli from the second modality and if the initial direction has a greater contrast when compared to the last direc-

tion then select the initial direction otherwise do not respond'
'attend exclusively to the second modality and if the first orientation is shown with a higher strength compared to the final direction then pick the first orientation otherwise do not respond'

**COMP2Mod1:**

*Train*

'attend only to the first modality and select the second direction if it is stronger than the first otherwise do not respond'
'opt for the final direction if it is presented with greater strength than the initial stimulus in the first modality otherwise do not respond'
'if the second stimulus is stronger than the first stimulus in the first modality than select the second stimulus otherwise do not respond'
'focus only on the first modality and choose the second orientation if is presented more intensely than the first direction otherwise do not respond'
'pick the final direction if it is displayed with higher intensity than the first direction in the first modality otherwise do not respond'
'pay attention only to the first modality and select the latter stimulus if it is stronger than the first stimulus otherwise do not respond'
'if the second orientation is greater than the first orientation in the first modality then respond to the second orientation otherwise do not respond'
'select the second stimulus if it is presented with higher strength than the first stimulus in the first modality otherwise do not respond'
'if the final orientation is stronger than the first orientation in the first modality than choose the last orientation otherwise do not respond'
'focus only on the first modality and go in the second direction if it is presented with higher intensity than the first direction otherwise do not respond'
'respond to the second stimulus if it has greater strength than the first stimulus in the first modality otherwise do not respond'
'attend only to the first modality and opt for the final direction if it is more intense than the initial direction otherwise do not respond'
'go in the direction of the latter stimulus if it has greater strength than the initial stimulus in the first modality otherwise do not respond'
'focus only on the first modality and choose the the final direction if it is stronger than the first direction otherwise do not respond'
'select the second orientation if it is greater than the first orientation in the first modality otherwise do not respond'

*Validation*

'only consider stimuli from the first modality and opt for the last direction if it appears with a higher contrast than the initial direction otherwise do not respond'
'focus solely on the stimuli that appear in the first modality and choose the latter stimulus if it is presented with more strength compared to that of the earlier stimulus otherwise do not respond'
'concentrate only on the first modality and if the latter orientation appears stronger compared to the initial orientation then opt for the latter orientation otherwise do not respond'
'attend exclusively to the first modality and select the final direction if it is presented with greater contrast compared to that of the earlier direction otherwise do not respond'
'consider only stimuli in the first modality and respond to the latter stimulus if is has a contrast which is higher compared to the initial stimulus otherwise do not respond'

**COMP2Mod2:**

*Train*

'focus only on the second modality and select the latter direction if it is stronger than the first direction otherwise do not respond'
'select the final direction if it is greater than the initial direction in the second modality otherwise do not respond'
'pay attention only to the second modality and pick the final orientation if it has higher intensity than the first orientation otherwise do not respond'
'if the final stimulus is presented with higher intensity than the first stimulus in the second modality then respond to the final stimulus otherwise do not respond'
'select the second orientation if it is displayed more intensely than the first orientation in the second modality otherwise do not respond'
'if the final stimulus is greater than the initial stimulus in the second modality than respond to the first stimulus otherwise do not respond'
'attend only to the second modality and if the final stimulus is stronger than the initial stimulus select the final stimulus otherwise do not respond'
'pick the second stimulus if it is presented with greater strength than the first stimulus in the second modality otherwise do not respond'
'choose the final orientation if it is greater than the initial orientation in the second modality otherwise do not respond'
'focus only on the second modality and pick the latter direction if it has more strength than the initial direction otherwise do not respond'
'go in the direction of the second stimulus if it has more intensity than the initial stimulus in the second modality otherwise do not respond'
'select the second direction if it is stronger than the first direction in the second modality otherwise do not respond'
'choose the final stimulus if it has higher intensity than the initial stimulus in the second modality otherwise do not respond'
'attend only to the second modality and select the final direction if it has higher intensity than the first direction otherwise do not respond'
'if the latter stimulus is stronger than the first stimulus in the second modality respond to the latter stimulus otherwise do not respond'

*Validation*

'concentrate only on stimuli which are displayed in the second modality and choose the last orientation if it has a contrast that is higher than the contrast of the earlier orientation otherwise do not respond'
'attend solely to the second modality and select the latter direction if it appears with a greater strength compared to that of the earlier direction otherwise do not respond'
'concentrate entirely on stimuli from the second modality and if the last stimulus has a larger contrast compared to that of the earlier stimulus then select the last stimulus otherwise do not respond'
'consider exclusively those stimuli displayed in the second modality and if the last direction is displayed with a higher contrast compared to the initial direction respond to the last direction otherwise do not respond'
'only consider stimuli which appear in the the second modality and pick the latter direction if it is displayed with greater intensity compared to the earlier direction otherwise do not respond'

**DMS:**

*Train*

'if the first and the second stimuli match then respond with that orientation otherwise do not respond'
'if the same directions are displayed then respond to the stimuli otherwise do not respond'
'if the stimuli match respond in the direction of the stimuli otherwise do not respond'
'when the two displayed directions are the same respond to the stimulus otherwise do not respond'
'if the stimuli match go in the displayed direction otherwise do not respond'
'select the displayed direction if the directions match otherwise do not respond'
'respond if the stimuli are displayed with the same orientation otherwise do not respond'
'if the first and second stimuli has the same orientation then respond otherwise do not respond'
'when the two stimuli match respond in the displayed direction otherwise do not respond'
'if the stimuli match then respond otherwise do not respond'
'if the displayed directions are identical then select the displayed direction otherwise do not respond'
'select the displayed orientation if the first and second stimuli are identical otherwise do not respond'
'pick the stimuli direction if the first and second direction are identical otherwise do not respond'
'choose the presented orientation if the first and second stimuli match otherwise do not respond'
'respond in the displayed direction if the first and second stimuli match otherwise do not respond'

*Validation*

'respond if stimuli are presented in the same directions otherwise do not respond'
'go in the displayed direction if the stimuli match otherwise do not respond'
'when displayed directions are the same respond to the stimulus otherwise do not respond'
'go in the direction displayed if the two stimuli match otherwise do not respond'
'select the stimuli direction if both directions are the same otherwise do not respond'

**DNMS:**

*Train*

'if the first and second stimuli are different then respond to the second stimuli otherwise do not respond'
'if the stimuli are mismatched go in direction of the second stimuli otherwise do not respond'
'when the displayed directions are distinct go in final direction otherwise do not respond'
'when stimuli are presented in different directions respond to the latter direction otherwise do not respond'
'respond in the final direction if stimuli are mismatched otherwise do not respond'
'go in the second direction when stimuli orientations are different otherwise do not respond'
'respond in the final displayed direction if stimuli are mismatched otherwise do not respond'
'if stimuli are mismatched respond in the latter direction otherwise do not respond'
'if displayed directions are distinct select the final direction otherwise do not respond'
'go in second displayed direction if stimuli are mismatched otherwise do not respond'
'if the first and second stimuli are different then select the final direction otherwise do not respond'
'select the second orientation if both presented orientations are distinct otherwise do not respond'
'choose the last direction if both displayed directions are mismatched otherwise do not respond'
'when the stimuli are distinct then select the final stimuli otherwise do not respond'
'respond to the second orientation if the presented orientations are different otherwise do not respond'

*Validation*

'if displayed directions are different go in the second direction otherwise do not respond'
'when the stimuli are distinct respond to the final stimulus otherwise do not respond'
'if the orientations are mismatched then go in the last direction otherwise do not respond'
'if the two stimuli mismatched respond to the second stimulus otherwise do not respond'
'select the second direction if both stimuli are presented in different directions otherwise do not respond'

**DMC:**

*Train*

'if the stimuli are on the same half of the display go in first direction otherwise do not respond'
'respond to the initial orientation if stimuli are in the same half of the display otherwise do not respond'
'go in first direction if directions are in same half of display otherwise do not respond'
'if the stimuli are in the same half of display respond in the first direction otherwise do not respond'
'if displayed orientations are in the same half respond in the first direction otherwise do not respond'
'respond in the initial direction if the stimuli occur in the same half of the display otherwise do not respond'
'when the stimuli are presented on the same half of the display go in the first direction otherwise do not respond'
'if the stimuli are on the same half of the display choose the first direction otherwise do not respond'
'when the displayed directions are in the same half select the initial direction otherwise do not respond'
'choose the first stimulus when both stimuli are presented on the same half of the display otherwise do not respond'
'select the first orientation if both orientation are in the same half of the display otherwise do not respond'
'if the initial and second direction are in the same half of the display then respond to the initial direction otherwise do not respond'
'select the initial stimulus if the initial and final stimuli are presented on the same half of the display otherwise do not respond'
'if the displayed directions are on the same half of the display then respond to the first direction otherwise do not respond'
'choose the first direction if both presented stimuli are on the same half of the display otherwise do not respond'

*Validation*

'go in the first orientation if both are displayed on the same side of the display otherwise do not respond'
'if the stimuli appear in the same half of the display respond to the initial direction otherwise do not respond'

'if stimuli are on the same half respond in the initial displayed direction otherwise do not respond'
'pick the first orientation if both stimuli appear on the same side of the display otherwise do not respond'
'select the first stimulus if both are presented on the same side of the display otherwise do not respond'

**DNMC:**

*Train*

'if stimuli are on different halves of display respond in the second direction otherwise do not respond'
'when the stimuli are presented are on opposing sides of the display go in the final direction otherwise do not respond'
'go in the final direction when stimuli are on distinct halves of the display otherwise do not respond'
'if the directions are on different halves select the second direction otherwise do not respond'
'when the stimuli appear in distinct halves choose the latter direction otherwise do not respond'
'select the final direction if stimuli are displayed on opposing sides of display otherwise do not respond'
'choose the latter direction when stimuli appear on different halves otherwise do not respond'
'if the stimuli are displayed on distinct sides then respond in second direction otherwise do not respond'
'select the final orientation if stimuli are on different sides of display otherwise do not respond'
'respond in the latter direction when directions are presented on different halves of the display otherwise do not respond'
'if the stimuli are presented on opposing sides of the display then select the second stimulus otherwise do not respond'
'if the first and second directions are displayed on opposite halves of the display respond in the final direction otherwise do not respond'
'go in the second direction when the stimuli are presented on opposite halves of the display otherwise do not respond'
'respond to the final orientation if both are presented on distinct halves otherwise do not respond'
'choose the final stimulus if both presented stimuli are on opposite halves of the display otherwise do not respond'

*Validation*

'pick the second direction if both stimuli are on different sides of the display otherwise do not respond'
'choose the final displayed orientation if they appear on opposing halves of the display otherwise do not respond'
'if the directions appear on distinct sides of the display respond to the last direction otherwise do not respond'
'if the stimuli are on different sides respond to the second direction otherwise do not respond'
'select the final orientation if both stimuli appear on opposite halves of the display otherwise do not respond'

#### 14 Rule Vectors for STRUCTURENET

The rule vectors used by STRUCTURENET are a combination of 10 basic dimensions which together describe the task set. The dimension are: {pro/anti, standard/RT, mod1/mod2, strongest/weakest, multi-sensory, confidence-based, first/second stimuli, longest/shortest stimuli, matching/non-matching, stimuli/category}. All other models we tested use a task information embedding dimension of 64, so in order to keep this consistent we use a similar method as for SIMPLENET and draw a set of 10 orthogonal vectors for each instantiation of STRUCTURENET. Each of these vectors corresponds to one of the basic dimensions in the task set and the rule vector for a given task is the linear combination of this orthogonal set and the underlying vector describing the subcomponenets of the task. The underlying vector for each task are:

'Go': {1, 1, 0, 0, 0, 0, 0, 0, 0, 0}

'AntiGo': {-1, 1, 0, 0, 0, 0, 0, 0, 0, 0}

'RTGo': {1, -1, 0, 0, 0, 0, 0, 0, 0, 0}

'AntiRTGo': {-1, -1, 0, 0, 0, 0, 0, 0, 0, 0}

'GoMod1': {1, 1, 1, 0, 0, 0, 0, 0, 0, 0}

'GoMod2': {1, 1, -1, 0, 0, 0, 0, 0, 0, 0}

'AntiGoMod1': {-1, 1, 1, 0, 0, 0, 0, 0, 0, 0}

'AntiGoMod2': {-1, 1, -1, 0, 0, 0, 0, 0, 0, 0}

'RTGoMod1': {1, -1, 1, 0, 0, 0, 0, 0, 0, 0}

'AntiRTGoMod1': {-1, -1, 1, 0, 0, 0, 0, 0, 0, 0}

'RTGoMod2': {1, -1, -1, 0, 0, 0, 0, 0, 0, 0}

'AntiRTGoMod2': {-1, -1, -1, 0, 0, 0, 0, 0, 0, 0}

'DM': {0, 0, 0, 1, 0, 0, 0, 0, 0, 0}

'AntiDM': {0, 0, 0, -1, 0, 0, 0, 0, 0, 0}

'MultiDM': {0, 0, 0, 1, 1, 0, 0, 0, 0, 0}

'AntiMultiDM': {0, 0, 0, -1, 1, 0, 0, 0, 0, 0}

'ConDM': {0, 0, 0, 1, 0, 1, 0, 0, 0, 0}

'ConAntiDM': {0, 0, 0, -1, 0, 1, 0, 0, 0, 0}

'DMMod1': {0, 0, 1, 1, 0, 0, 0, 0, 0, 0}

'DMMod2': {0, 0, -1, 1, 0, 0, 0, 0, 0, 0}

'AntiDMMod1': {0, 0, 1, -1, 0, 0, 0, 0, 0, 0}

'AntiDMMod2': {0, 0, -1, -1, 0, 0, 0, 0, 0, 0}

'Dur1': {0, 0, 0, 0, 0, 0, 1, 1, 0, 0}

'Dur2': {0, 0, 0, 0, 0, 0, -1, 1, 0, 0}

'MultiDur1': {0, 0, 0, 0, 1, 0, 1, 1, 0, 0}

'MultiDur2': {0, 0, 0, 0, 1, 0, -1, 1, 0, 0}

'AntiDur1': {0, 0, 0, 0, 0, 0, 1, -1, 0, 0}

'AntiDur2': {0, 0, 0, 0, 0, 0, -1, -1, 0, 0}

‘AntiMultiDur1’: {0, 0, 0, 0, 1, 0, 1, -1, 0, 0}
‘AntiMultiDur2’: {0, 0, 0, 0, 1, 0, -1, -1, 0, 0}
‘Dur1Mod1’: {0, 0, 1, 0, 0, 0, 1, 1, 0, 0}
‘Dur1Mod2’: {0, 0, -1, 0, 0, 0, 1, 1, 0, 0}
‘Dur2Mod1’: {0, 0, 1, 0, 0, 0, -1, 1, 0, 0}
‘Dur2Mod2’: {0, 0, -1, 0, 0, 0, -1, 1, 0, 0}
‘COMP1’: {0, 0, 0, 1, 0, 0, 1, 0, 0, 0}
‘COMP2’: {0, 0, 0, 1, 0, 0, -1, 0, 0, 0}
‘MultiCOMP1’: {0, 0, 0, 1, 1, 0, 1, 0, 0, 0}
‘MultiCOMP2’: {0, 0, 0, 1, 1, 0, -1, 0, 0, 0}
‘AntiCOMP1’: {0, 0, 0, -1, 0, 0, 1, 0, 0, 0}
‘AntiCOMP2’: {0, 0, 0, -1, 0, 0, -1, 0, 0, 0}
‘AntiMultiCOMP1’: {0, 0, 0, -1, 1, 0, 1, 0, 0, 0}
‘AntiMultiCOMP2’: {0, 0, 0, -1, 1, 0, -1, 0, 0, 0}
‘COMP1Mod1’: {0, 0, 1, 1, 0, 0, 1, 0, 0, 0}
‘COMP1Mod2’: {0, 0, -1, 1, 0, 0, 1, 0, 0, 0}
‘COMP2Mod1’: {0, 0, 1, 1, 0, 0, -1, 0, 0, 0}
‘COMP2Mod2’: {0, 0, -1, 1, 0, 0, -1, 0, 0, 0}
‘DMS’: {0, 0, 0, 0, 0, 0, 0, 0, 1, 1}
‘DNMS’: {0, 0, 0, 0, 0, 0, 0, 0, -1, 1}
‘DMC’: {0, 0, 0, 0, 0, 0, 0, 0, 1, -1}
‘DNMC’: {0, 0, 0, 0, 0, 0, 0, 0, -1, -1}

#### 15 Novel Decoded Instructions

Top five most common novel instructions produced from our language production network.

**Sensorimotor-RNN trained on All Tasks and Language Production trained on All Tasks: Go:** 'respond in the direction of the stimulus with maximal strength', 18; 'respond in the displayed direction if the first strength otherwise do not respond', 17; 'respond in the direction of the stimulus if you are confident otherwise do not respond', 17; 'respond in the direction of the stimulus stimulus if you are confident otherwise do not respond', 5; 'respond in the direction of the stimulus with', 4; **AntiGo:** 'pay attention only to the second modality and respond in the opposite direction', 16; 'respond with the opposite direction', 12; 'respond in the opposite direction', 10; 'respond in the reverse of', 9; 'respond in the direction of the stimulus presented with the strongest', 8; **RTGo:** 'go in the same direction at stimulus onset', 44; 'go in the displayed direction at stimulus', 30; 'choose the same direction as the stimulus is shown', 18; 'choose the direction immediately when the stimulus is shown', 8; 'pay attention only to the first modality and respond', 7; **AntiRTGo:** 'pay attention only to the second modality and choose the opposite of the orientation at stimulus onset', 7; 'respond in the converse of the direction in the second modality at stimulus onset', 6; 'go in the opposite of the display direction at stimulus onset', 5; 'go in the opposite of the direction as soon as the stimulus appears', 5; 'go in the reverse of the direction as soon as stimulus stimulus', 5; **GoMod1:** 'focus only on the first modality and choose the shown orientation at stimulus onset', 9; 'focus on the first modality and choose the stimulus that appears', 8; 'go in the direction of the stimulus presented', 7; 'focus on on the first modality and the stimulus that appears', 6; 'go in the direction of the stimulus in the second modality', 5; **AntiGoMod1:** 'attend only to the first modality and select the opposite of the stimulus displayed there', 12; 'focus on the first modality and respond in the opposite of the displayed direction', 10; 'focus on the first modality and respond in the opposite of the direction that appears there', 6; 'pay attention only to the first modality and choose the reverse of the orientation displayed', 5; 'attend only to the first modality and select the opposite of the stimulus displayed', 4; **GoMod2:** 'focus only on the second modality and choose the orientation that appears there', 15; 'attend only to the second modality and pick the displayed direction', 9; 'choose the stimulus presented with minimal intensity', 7; 'attend only to the second modality and pick the displayed orientation', 6; 'attend only to the second modality and select the orientation displayed there', 6; **AntiGoMod2:** 'attend only to the second modality and pick the reverse of the orientation presented', 23; 'attend only to the second modality and pick the opposite direction', 11; 'focus only on the second modality and respond in the reverse of the direction that appears there', 4; 'attend to the stimulus in the second modality and go in the opposite direction', 3; 'attend only to the second modality and pick the opposite of the orientation displayed', 3; **RTGoMod1:** 'attend only to the first modality and select the orientation immediately', 16; 'pay attention to the first modality and select the displayed direction immediately', 11; 'attend only to the first modality and select the direction immediately', 11; 'pay attention to the first modality and select the orientation immediately', 5; 'pay attention to the first modality and select the direction of', 2; **AntiRTGoMod1:** 'pay attention to the first modality and respond in the reverse of the stimulus immediately', 13; 'select the reverse of the orientation in the first modality at stimulus onset', 9; 'go in the reverse of the orientation displayed in the first orientation at stimulus onset', 5; 'pay attention only to the first modality and opt for the opposite at stimulus onset', 3; 'respond with the opposite of the stimulus in the first modality at stimulus onset', 3; **RTGoMod2:** 'attend only to the second modality and opt for the displayed orientation at stimulus onset', 29; 'pay attention to the second modality and pick the stimulus orientation at stimulus onset', 23; 'pay attention to the second modality and go in the second modality immediately', 14; 'select the orientation in the second modality at stimulus onset', 13; 'attend only to the second modality and respond to the orientation at stimulus onset', 11; **AntiRTGoMod2:** 'pay attention to the second modality and opt for the opposite direction at stimulus onset', 24; 'go in the opposite of the stimulus in the second modality at stimulus onset', 15; 'pay attention to the second modality and opt for the opposite of the second modality', 5; 'pay attention to the second modality and respond the reverse of the displayed direction at stimulus onset', 4; 'pay attention to the second modality and respond to the reverse of the stimulus appears', 4; **DM:** 'respond in the direction of the strongest stimulus if you are confident otherwise do not respond', 8; 'choose the orientation with highest intensity integrated over', 8; 'choose the direction with highest intensity', 8; 'choose the orientation presented with highest intensity', 8; 'go in the direction presented with highest intensity', 5; **AntiDM:** 'respond to the orientation presented with minimal strength', 13; 'respond in the direction of minimal strength stimulus strength', 11; 'respond to the direction presented with the strength', 10; 'select the orientation with lowest strength', 6; 'select the direction presented with least strength', 6; **MultiDM:** 'respond in the direction with greatest strength between two stimuli', 9; 'respond in the direction with highest intensity between two stimuli', 8; 'respond in the direction with highest overall strength between', 7; 'go in the direction with highest intensity', 6; 'respond in the direction of highest intensity between two stimuli', 6; **AntiMultiDM:** 'go in the direction of smallest intensity combined over both two', 2; 'respond in the direction with weakest combined value between two stimuli', 2; 'select the direction with weakest average value', 2; 'respond to the orientation with lowest combined value between modalities', 2; 'respond to the orientation with lowest combined value between two modalities', 2; **DMMod1:** 'choose the stimulus presented with highest intensity in the first modality', 28; 'choose the stimulus which with highest intensity in the first modality', 11;

'select the stimulus with maximal strength in the first modality', 9; 'select the stimulus displayed with maximal intensity in the first modality', 8; 'select the stimulus in the first modality select the first modality', 7; **AntiDMMod1:** 'select the stimulus in the first modality that is presented with least intensity', 12; 'select the stimulus in the first modality that has lowest strength', 5; 'focus only on the first modality and choose the orientation with lowest intensity', 4; 'attend to the first modality and select the weakest direction', 4; 'focus only on the first modality and choose the orientation presented with lowest intensity', 2; **DMMod2:** 'respond to the stimulus displayed with maximal intensity in the second modality', 9; 'focus only on the second modality and select the direction displayed with greatest strength', 6; 'focus only to the second modality and respond to the second modality', 5; 'attend only to the second modality and pick the orientation that appears there', 5; 'opt for the direction displayed in the second modality with', 4; **AntiDMMod2:** 'attend to the second modality and choose the weakest stimulus is presented with the stimulus', 12; 'attend to the second modality and choose the orientation with lowest strength', 8; 'attend to the second modality and choose the weakest stimulus is presented with the', 5; 'attend to the second modality and opt for the weakest with weakest intensity', 3; 'focus on the second modality and choose the weakest direction that appears weakest', 2; **ConDM:** 'go in the direction of the stimulus strength if you are sure in your answer otherwise do not respond', 4; 'respond to the stimulus presented with highest intensity if you are sure of your answer otherwise do not respond', 4; 'respond in the direction of highest intensity if you are confident otherwise do not respond', 4; 'choose the strongest orientation if you have high confidence otherwise do not not', 4; 'opt for the direction with highest intensity if you are sure of your answer otherwise do not respond', 3; **ConAntiDM:** 'if you have high confidence select the stimulus displayed with least intensity otherwise do not respond', 6; 'if you are confident in your answer select the weakest stimulus otherwise do not respond', 6; 'if you have high confidence choose the stimulus with least intensity otherwise do not respond', 5; 'respond to the orientation with lowest strength if you are sure about your answer otherwise do not respond', 3; 'if you have high confidence choose the orientation with least strength otherwise otherwise do not respond', 3; **COMP1:** 'choose the first stimulus if it is presented with higher intensity relative to the second stimulus otherwise do not respond', 16; 'when the initial stimulus has higher value than the final stimulus respond in the first direction otherwise do not respond', 10; 'when the first stimulus is presented with greater strength than the second stimulus respond to the first direction otherwise do not respond', 7; 'when the first stimulus is presented with the higher intensity than the latter select the first stimulus otherwise do not respond', 5; 'if the first stimulus has higher intensity than the final go in the first stimulus otherwise do not respond', 5; **COMP2:** 'when the final stimulus is more intense than the first respond to the second stimulus otherwise do not respond', 14; 'if the final stimulus has higher value than the first stimulus select the final orientation otherwise do not respond', 13; 'when the presented with', 9; 'when the final stimulus has higher respond to the initial stimulus otherwise do not respond', 9; 'when the final stimulus has higher value than the first stimulus otherwise do not respond', 8; **MultiCOMP1:** 'if the intensity of the first stimulus integrated over modalities is greater than the second respond to the first direction otherwise do not respond', 7; 'select the first direction when it has higher joint strength over modalities than the final otherwise do not respond', 7; 'if the intensity of the first direction when combined over modalities is modalities than the second orientation otherwise do not respond', 6; 'if the first direction is presented with higher intensity than the second respond to the first direction otherwise do not respond', 6; 'if the joint intensity of the first intensity is combined over both modalities than the second direction otherwise do not respond', 6; **MultiCOMP2:** 'if the final direction is presented with greater overall strength across modalities than the initial direction then choose the final direction otherwise do not respond', 19; 'if the final stimulus has higher overall value both modalities than the initial direction then select the final stimulus otherwise do not respond', 12; 'if the final direction is presented with greater overall strength across modalities than the initial direction then select the final direction otherwise do not respond', 6; 'pick the final direction if it has less strength than the initial direction otherwise do not respond', 5; 'if the final stimulus has higher overall value strength both modalities than the initial direction then select the final stimulus otherwise do not respond', 4; **AntiCOMP1:** 'if the first stimulus is presented with less strength than the final direction then select the first stimulus otherwise do not respond', 25; 'if the initial stimulus is presented with less intensity than the second stimulus then select the initial stimulus otherwise do not respond', 19; 'if the initial stimulus is presented with less intensity than the second direction then select the initial stimulus otherwise do not respond', 10; 'if the first direction is less intense than the final direction then select the first direction otherwise do not respond', 8; 'choose the first direction if it has less strength than the second direction if do not respond', 5; **AntiCOMP2:** 'if the second direction is presented with less intensity than the first direction than the second direction otherwise do not respond', 31; 'if the latter direction is presented with less intensity than the first direction then select the second stimulus otherwise do not respond', 13; 'if the latter direction is presented with less intensity than the first direction otherwise do not respond', 7; 'if the second orientation is presented with less strength than the first direction then select the final orientation otherwise do not respond', 6; 'if the latter direction is less intense than the initial direction then respond to the latter direction otherwise do not respond', 4; **AntiMultiCOMP1:** 'select the initial direction if it is displayed for less considered than the final direction averaged over both modalities otherwise do not respond', 4; 'if the first orientation has less intensity combined over both modalities than the second orientation then select the first orientation otherwise do not respond', 3; 'if the first direction is presented with less intensity than the final direction averaged over both modalities then choose the first direction otherwise do not respond', 3; 'opt for the initial direction if it lasts for a of time period of time than the second stimulus otherwise do not respond', 2; 'select the initial orientation if it has a duration than the final orientation otherwise do not respond', 2; **AntiMultiCOMP2:** 'if the final orientation is presented with less intensity combined over both modalities than the first orientation select the final orientation otherwise do not respond', 8; 'opt for the second stimulus if it has lower intensity averaged over both modalities than the first stimulus otherwise do not respond', 3; 'select the second orientation if it has a shorter duration averaged over both modalities than the initial orientation otherwise do not respond', 3; 'if the final orientation is presented with less intensity combined over both modalities than the initial orientation then select the latter orientation otherwise do not respond', 3; 'pick the final direction if it is weaker than the first direction averaged over both modalities otherwise do not respond', 3; **COMP1Mod1:** 'attend only to the first modality and choose the first direction if the more intensity than the first direction otherwise do not respond', 11; 'attend only to the first modality and choose the first direction if it has more strength than the final orientation otherwise do not respond', 5; 'attend only to the first modality and choose the first direction if is stronger than the second stimulus otherwise do not respond', 5; 'if the first orientation is stronger than the second orientation in the first modality then respond to the first orientation otherwise do not respond', 4; 'attend only to the first modality and select the initial direction if it has more strength than the final stimulus otherwise do not respond', 4; **COMP2Mod1:** 'focus only on the first modality and choose the second direction if it is stronger than the first direction otherwise do not respond', 14; 'pick the final direction if it is weaker than the first direction in the first modality otherwise do not respond', 9; 'focus only on the first modality and choose the final orientation if it is presented for more intensity than the first direction otherwise do not respond', 8; 'focus only on the first modality and choose the final orientation if it is presented than the first direction otherwise do not respond', 6; 'select the second stimulus if it is greater than the first stimulus in the first modality otherwise do not respond', 5; **COMP1Mod2:** 'if the initial stimulus is more intense than the last stimulus in the second modality then respond to the second stimulus otherwise do not respond', 10; 'if the first stimulus is stronger than the final stimulus in the second modality then select the initial stimulus otherwise do not respond', 8; 'attend only to the second modality and select the first direction if it has greater strength than the second direction otherwise do not respond', 5; 'attend only to the second modality and if the first orientation is greater than the second stimulus respond to the first direction otherwise do not respond', 5; 'if the initial stimulus is more intense than the last direction in the second modality then respond to the initial stimulus otherwise do not respond', 3; **COMP2Mod2:** 'pick the second direction if it appears more than the first direction in the second modality otherwise do not respond', 13; 'pick the second direction if it has more intensity than the first direction in the second modality otherwise do not respond', 9; 'if the final stimulus is greater than the first stimulus in the second modality than respond to the first stimulus otherwise do not respond', 5; 'focus only on the second modality and pick the latter orientation if it has more strength than the first direction otherwise do not respond', 4; 'pick the second stimulus if it is more intense than the first stimulus in the second modality otherwise do not respond', 3; **Dur1:** 'if the first stimulus is presented for a greater period of time than the latter stimulus then respond to the first modality otherwise do not respond', 10; 'pick the first direction if it appears for less time than the latter direction otherwise do not respond', 10; 'opt for the initial stimulus if it has a shorter duration than the final stimulus otherwise do not respond', 9; 'pick the first direction if it lasts for a greater period of time than the latter direction otherwise do not respond', 7; 'pick the first direction if it appears for less time than the final direction otherwise do not respond', 6; **Dur2:** 'if the latter stimulus appears for more time than the first stimulus respond to the second stimulus otherwise do not respond', 35; 'if the final stimulus is presented for more time than the first stimulus respond to the final stimulus otherwise do not respond', 11; 'if the second direction is displayed for more time than the first direction respond to the final direction otherwise do not respond', 6; 'if the latter stimulus appears for more time than the first stimulus than select the latter stimulus otherwise do not respond', 5; 'if the second direction is displayed for a greater period of time than the first direction otherwise do not respond', 5; **MultiDur1:** 'if the first direction appears for a longer period of time then the latter direction over both modalities then select that direction otherwise do not respond', 12; 'pick the initial direction if it last for a longer period of time than the final direction otherwise do not respond', 9; 'if the initial orientation appears for longer than second second orientation when considered across both modalities than the second orientation otherwise do not respond', 7; 'pick the first direction if it appears for a of time than the final direction combined over both modalities otherwise do not respond', 7; 'pick the initial direction if it appears for a longer period of time than the final direction otherwise do not respond', 7; **MultiDur2:** 'if the second stimulus has a greater duration than the first stimulus combined over both modalities than select the second stimulus otherwise do not respond', 14; 'if the second stimulus has a duration which lasts for longer than the first stimulus when combined over both modalities then select the first stimulus otherwise do not respond', 11; 'if the duration of the final stimulus is longer than the initial stimulus combined across modalities then choose the first stimulus otherwise do not respond', 9; 'if the second stimulus has a greater duration than the first stimulus combined over both modalities then select the second stimulus otherwise do not respond', 7; 'if the second stimulus has a duration which is longer than the initial stimulus combined across across modalities then select the first stimulus otherwise do not respond', 7; **AntiDur1:** 'if the initial direction is displayed for less time than the initial direction select the initial direction otherwise do not respond', 13; 'if the initial direction is shorter than the final direction then choose the initial direction otherwise do not respond', 9; 'pick the first direction if it lasts for a shorter period of time than the final otherwise otherwise do not respond', 5; 'pick the first stimulus if it is shorter than the final direction otherwise do not respond', 3; 'if the first stimulus has a shorter duration than the latter stimulus otherwise do not respond', 3; **AntiDur2:** 'if the latter direction appears for a shorter duration than the first direction select the latter direction otherwise do not respond', 19; 'if the final stimulus is shorter than the initial direction choose the final direction otherwise do not respond', 18; 'respond to the final direction if it is shorter than the first direction otherwise do not respond', 12; 'if the latter direction is shorter than the initial direction pick the latter initial do not respond', 10; 'if the second direction is displayed for less time than the initial direction combined over both modalities otherwise do not respond', 9; **AntiMultiDur1:** 'if the initial stimulus is shorter than the latter stimulus when combined over both modalities than the stimulus stimulus otherwise do not respond', 18; 'if the initial stimulus has a shorter duration when combined over both modalities than the latter stimulus then select the initial stimulus otherwise do not respond', 12; 'if the initial stimulus has a shorter duration over both modalities than the initial stimulus then select the final stimulus otherwise do not respond', 6; 'if the first orientation appears for less time averaged over both modalities than the second orientation respond to the initial orientation otherwise do not respond', 4; 'if the first stimulus is shorter than the latter stimulus combined over both modalities then choose the first stimulus otherwise do not respond', 3; **AntiMultiDur2:** 'if the final stimulus is shorter than the first stimulus averaged over both modalities then select the final stimulus otherwise do not respond', 11; 'if the final orientation is shorter than the initial orientation combined over both modalities then choose the final orientation otherwise do not respond', 8; 'if the second stimulus is shorter than the first stimulus combined over both modalities then select the second stimulus otherwise do not respond', 6; 'respond to the final direction if it appears for less time averaged over both modalities than the first direction otherwise do not respond', 5; 'respond to the second direction if it is displayed for less time than the initial stimulus considered over both modalities otherwise do not respond', 5; **Dur1Mod1:** 'focus only on the first modality and select the initial stimulus if it has a duration than the final stimulus otherwise do not respond', 14; 'pay attention only to the first modality and select the initial stimulus if it lasts for a longer period of time than the latter stimulus otherwise do not respond', 7; 'if the first direction which has greater strength than the final stimulus in the first modality otherwise do not respond', 6; 'pay attention only to the first modality and select the initial stimulus if it is longer for longer than the final stimulus otherwise do not respond', 6; 'pay attention only to the first modality and opt for the first stimulus if it is presented for longer than the final stimulus otherwise do not respond', 6; **Dur2Mod1:** 'select the reverse of the displayed direction', 6; 'select the reverse of the orientation displayed', 6; 'respond to the second stimulus if it is displayed for a greater period of time than the first stimulus otherwise to not respond', 4; 'focus only on the second modality and pick the direction with lowest strength', 4; 'focus only on the first modality and choose the final orientation if it is a than initial initial direction otherwise do not respond', 4; **Dur1Mod2:** 'attend only to the second modality and select the initial stimulus if it lasts for a longer period of time than the latter stimulus otherwise do not respond', 14; 'attend only to the second modality and pick the first stimulus if it lasts for a greater period of time than the latter stimulus otherwise do not respond', 7; 'if the first direction if is lasts for longer than the second direction in the first modality then respond to the first direction otherwise do not respond', 6; 'if the first direction if is lasts for longer than the second direction in the second modality then respond to the first direction otherwise do not respond', 6; 'if the first direction if is appears for more time than the second direction in the first modality then respond to the first direction otherwise do not respond', 5; **Dur2Mod2:** 'focus respond to the second modality and select the latter direction if it is displayed for more time than the first direction otherwise do not respond', 8; 'pay attention only to the second modality and pick the latter stimulus if it is displayed for a greater period of time than the first stimulus otherwise do not respond', 5; 'pay attention only to the second modality and pick the latter stimulus if it is presented for a greater period of time than the first direction otherwise do not respond', 4; 'attend only to the second modality and select the latter stimulus if it is displayed for a greater period of time than the first stimulus otherwise do not respond', 4; 'respond to the second stimulus if it is displayed for more time than the first stimulus in the first modality otherwise do not respond', 4; **DMS:** 'if the stimuli match then respond in the same direction otherwise do not respond', 11; 'if the first and the second direction match respond', 7; 'if the stimuli match respond in the displayed direction otherwise do not respond', 6; 'when the two displayed stimulus respond to the stimulus with the intensity otherwise do not respond', 6; 'when the two displayed directions are the same respond to the first orientation otherwise do not respond', 6; **DNMS:** 'if the stimuli are mismatched go in direction if stimuli otherwise do not respond', 9; 'if displayed directions are distinct select the final direction if stimuli are otherwise do not respond', 9; 'if stimuli are mismatched go in the final direction if stimuli are otherwise do not respond', 6; 'when the stimuli are distinct then respond in the final direction otherwise do not respond', 5; 'if the stimuli are mismatched go in direction direction if stimuli otherwise do not respond', 4; **DMC:** 'if the stimuli are in the same half of the display then respond in the direction direction otherwise do not respond', 30; 'if the stimuli are in the same half of the display choose the first direction otherwise do not respond', 12; 'when the displayed directions are in the same half select the initial stimulus otherwise do not respond', 8; 'when the initial stimulus when the same half are the display otherwise do not respond', 7; 'choose the first direction if both presented stimuli are presented on the same half of the display otherwise do not respond', 5; **DNMC:** 'if the stimuli are presented on opposing sides of the display then the second stimulus otherwise do not respond', 12; 'if the stimuli are presented on opposing sides of the display go in the final direction otherwise do not respond', 6; 'choose the final direction if stimuli are on different halves of the display otherwise do not respond', 5; 'if stimuli are on different halves respond to the latter direction otherwise do not respond', 5; 'when the stimuli appear distinct then respond in the first direction otherwise do not respond', 4

**Sensorimotor-RNN trained with Tasks Held Out and Language Production trained on All Tasks: Go:** 'respond to the stimulus with maximal intensity if you are sure of your decision otherwise do not respond', 10; 'respond in the direction of the stimulus presented stimulus', 7; 'respond to the stimulus presented with highest intensity', 5; 'respond in the direction of the strongest stimulus otherwise do not respond', 5; 'respond to the stimulus with greatest strength', 5; **AntiGo:** 'select the reverse of the displayed direction', 21; 'opt for the reverse of the displayed stimulus', 15; 'opt for the opposite of the displayed direction', 14; 'pay attention only to the second modality and respond in the reverse of the displayed direction', 10; 'select the reverse of the stimulus displayed in', 10; **RTGo:** 'pay attention to the second modality and choose the direction immediately', 39; 'respond in direction the direction of the stimulus immediately', 33; 'choose the orientation that appears with the same', 33; 'choose the direction that appears select the stimulus immediately', 29; 'choose the direction displayed as soon as the stimulus immediately', 13; **AntiRTGo:** 'respond with the opposite of the orientation in the first modality at stimulus onset', 18; 'opt for the opposite direction at stimulus onset as stimulus appears', 13; 'choose the reverse of the displayed direction at stimulus onset', 9; 'respond with the opposite of the orientation immediately', 7; 'respond with the opposite of the orientation', 7; **GoMod1:** 'respond to the direction presented with the lowest strength', 16; 'attend to the first modality and select the displayed direction', 11; 'choose the least intense orientation with lowest intensity', 7; 'respond to the direction presented with the minimal strength', 7; 'attend to the first modality and respond in the direction of the displayed stimulus', 5; **AntiGoMod1:** 'attend to the stimulus in the first modality and respond in the opposite direction', 18; 'choose the opposite of the stimulus orientation in the first modality', 8; 'focus on the first modality and respond in the opposite direction of the stimulus displayed', 7; 'focus on the first modality and respond in the opposite direction of the stimulus that appears', 7; 'respond to the stimulus of the opposite', 5; **GoMod2:** 'respond in the direction of the stimulus displayed in the second modality', 13; 'respond in the direction that appears in the second modality', 12; 'pick the stimulus orientation with the lowest strength', 6; 'opt for the orientation of the stimulus displayed in the second modality', 6; 'respond to the direction which appears with highest strength in the second modality', 5; **AntiGoMod2:** 'respond in the opposite of the stimulus displayed in the second modality', 8; 'respond to the opposite of the orientation in the second modality', 6; 'respond in the opposite of the stimulus orientation from the second modality', 6; 'respond to the stimulus in the second modality and respond in the opposite of', 6; 'pick the opposite of the stimulus displayed in the second modality', 5; **RTGoMod1:** 'select the orientation that appears in the first direction at stimulus onset', 36; 'immediately respond to the direction displayed in the first direction immediately', 22; 'select the orientation that appears displayed in the first modality', 21; 'immediately respond to the orientation in the first modality immediately', 18; 'select the orientation that appears displayed in the first modality at stimulus onset', 11; **AntiRTGoMod1:** 'attend to the first modality and opt for the reverse direction at stimulus onset', 7; 'focus on the first modality and opt for the reverse of the stimulus immediately', 6; 'pay attention to the first modality and respond in the reverse of the stimulus immediately', 4; 'pay attention to the first modality and choose the reverse of the orientation at stimulus onset', 3; 'focus on the first modality and select the reverse of the stimulus immediately', 3; **RTGoMod2:** 'pay attention to the second modality and opt for the displayed stimulus onset', 23; 'attend to the second modality and opt for the displayed direction at stimulus onset', 22; 'attend to the second modality and opt for the displayed orientation at stimulus onset', 17; 'pay to the second modality and go in the displayed direction immediately', 5; 'attend to the second modality and opt for the orientation at stimulus onset', 4; **AntiRTGoMod2:** 'attend to the second modality and go in the opposite of the displayed direction immediately', 14; 'go in the opposite of the stimulus in the second modality at stimulus onset', 12; 'go in the reverse of the direction in the second modality immediately', 8; 'pay attention to the second modality and pick the reverse of the stimulus immediately', 6; 'attend to the second modality and go in the opposite of the stimulus immediately', 6; **DM:** 'respond to the direction of highest intensity', 18; 'choose the direction presented with highest intensity', 9; 'respond to the stimulus presented with highest intensity', 5; 'respond to the direction of highest intensity the stimulus', 5; 'select the direction with greatest average value over modalities', 5; **AntiDM:** 'respond to the stimulus presented with minimal strength intensity', 12; 'respond to the direction presented with the lowest strength', 11; 'respond to the stimulus presented with minimal strength', 9; 'respond to the stimulus with minimal strength between the stimulus with minimal intensity', 7; 'respond to the stimulus presented with minimal strength if you are sure', 6; **MultiDM:** 'go in the direction with', 8; 'choose the orientation with highest average intensity', 8; 'choose the direction with highest intensity between two stimuli', 8; 'respond in the direction with', 7; 'select the orientation with highest average strength', 6; **AntiMultiDM:** 'choose the direction which has the weakest joint intensity', 10; 'respond in the direction with minimal combined across both modalities', 6; 'select the orientation with least average intensity both modalities', 6; 'respond to the direction presented with the stimulus strength', 5; 'choose the direction with weakest weakest value between two stimuli', 5; **DMMod1:** 'respond to the orientation in the first modality that is strongest', 9; 'select the orientation in the that that intensity', 9; 'respond to the stimulus in the first modality that is strongest', 8; 'focus only on the first modality and choose the shown that appears strongest', 4; 'focus only on the first modality and choose the strongest that appears strongest', 4; **AntiDMMod1:** 'attend to the first modality and select the direction that appears weakest', 15; 'attend to the first modality and select the direction that is displayed with minimal strength', 15; 'focus on the first modality and choose the orientation with lowest intensity', 14; 'attend to the first modality and select the direction with least strength', 6; 'focus on the first modality and respond in the orientation orientation', 4; **DMMod2:** 'respond to the direction which appears with intensity in the second modality', 27; 'focus only on the second modality and pick the stimulus that appears there', 15; 'choose the direction in the second modality that is displayed with with intensity', 10; 'pick the stimulus in the second modality that is presented with maximal intensity', 5; 'focus on the second modality and respond to the orientation displayed with highest intensity', 5; **AntiDMMod2:** 'pick the stimulus in the second modality that appears minimal', 15; 'pick the direction in the second modality that appears weakest', 11; 'focus on the stimulus modality select the weakest stimulus that appears', 10; 'attend to the second direction in the first modality that', 10; 'focus on the second modality and choose the weakest orientation', 10; **ConDM:** 'respond to the stimulus presented with highest intensity if you are sure of your answer otherwise do not respond', 10; 'respond in the direction of greatest strength', 4; 'respond to the stimulus presented with highest intensity if you are sure otherwise do not respond', 4; 'respond to the stimulus displayed with highest intensity if you are sure of your decision otherwise do not respond', 4; 'respond in the direction with greatest strength if you are confident otherwise do not respond', 3; **ConAntiDM:** 'choose the orientation with lowest intensity', 13; 'choose the stimulus with lowest strength if you are confident in your decision otherwise do not respond', 8; 'respond to the stimulus presented with minimal strength', 7; 'respond in the direction with minimal strength', 6; 'choose the stimulus presented with the lowest strength', 5; **COMP1:** 'when the two stimuli are presented with higher intensity than the latter direction otherwise do not respond', 12; 'if the first stimulus is presented with the higher intensity than the second stimulus then select the first stimulus otherwise do not respond', 9; 'if the first stimulus is presented with the higher intensity than the second stimulus select the first stimulus otherwise do not respond', 9; 'choose the initial stimulus when it is presented with greater intensity than the final stimulus otherwise do not respond', 7; 'when the initial stimulus is presented with greater strength than the second stimulus respond to the initial direction otherwise do not respond', 7; **COMP2:** 'when the final stimulus is presented with the higher intensity than the first orientation otherwise do not respond', 9; 'when the final stimulus is presented with the presented intensity than the first respond in the final orientation otherwise do not respond', 6; 'when the final stimulus has greater value than the first than the final stimulus otherwise do not respond', 5; 'when the final stimulus is presented with the presented intensity than the first orientation otherwise do not respond', 5; 'respond to the final stimulus if it has greater strength than the first otherwise do not respond', 5; **MultiCOMP1:** 'if the first direction is presented with higher intensity averaged over modalities than the second direction then the first direction otherwise do not respond', 25; 'choose the initial direction when the joint strength over both modalities than the final direction otherwise do not respond', 15; 'choose the initial direction when combined over strength over modalities is greater than the final direction otherwise do not respond', 11; 'choose the initial direction when the joint strength over the higher modalities than the final direction otherwise do not respond', 10; 'if the joint intensity of the first directions is higher than the display then respond to the first direction otherwise do not respond', 9; **MultiCOMP2:** 'select the final direction if it has greater joint intensity over modalities than the initial direction otherwise do not respond', 13; 'select the final direction if it has greater joint intensity than the initial direction over both modalities otherwise do not respond', 9; 'if the final direction is represented has higher joint intensity over both modalities than the initial direction then respond to the final stimulus otherwise do not respond', 7; 'if the final direction is presented with greater overall strength across modalities than the initial direction select the final direction otherwise do not respond', 7; 'if the final direction is represented has higher joint intensity over both modalities than the initial direction then select the final stimulus otherwise do not respond', 5; **AntiCOMP1:** 'if the first stimulus is shorter than the second stimulus respond to the first stimulus otherwise do not respond', 13; 'choose the initial orientation if it is presented with lower intensity than the latter orientation otherwise do not respond', 10; 'if the initial stimulus is presented with less strength than the latter direction then select the initial stimulus otherwise do not respond', 7; 'if the first stimulus is presented with less strength than the latter direction then select the first stimulus otherwise do not respond', 7; 'respond to the initial orientation if stimuli are intensity in the same half otherwise do not respond', 7; **AntiCOMP2:** 'if the second stimulus has lower intensity than the first stimulus then select the second stimulus otherwise do not respond', 14; 'if the final stimulus is weaker than the initial stimulus respond to the initial stimulus otherwise do not respond', 10; 'if the final direction is displayed with less intensity than the first direction then select the final direction otherwise do not respond', 9; 'if the latter direction is displayed with less intensity than the first direction then select the latter direction otherwise do not respond', 5; 'respond to the final direction if it is displayed with less intensity combined over both modalities than the first direction otherwise do not respond', 4; **AntiMultiCOMP1:** 'if the initial direction is presented with less intensity than the second direction then select the initial direction otherwise do not respond', 9; 'if the initial direction has less strength than the final direction then select the initial direction otherwise do not respond', 7; 'opt for the initial orientation if it has lower intensity averaged over both modalities otherwise do not respond', 7; 'if the initial orientation has lower intensity averaged over both modalities than the second orientation otherwise do not respond', 7; 'if the initial direction is less intense than the final direction then select the initial direction otherwise do not respond', 6; **AntiMultiCOMP2:** 'respond to the second direction if it is weaker than the initial direction otherwise do not respond', 6; 'go in the second direction when it has less combined intensity over both modalities than the initial direction otherwise do not respond', 5; 'respond to the latter direction if it is displayed for less time than the initial direction otherwise do not respond', 4; 'respond to the latter direction if it is displayed for less time than the first direction otherwise do not respond', 3; 'respond to the latter direction if it is weaker than the initial direction otherwise do not respond', 3; **COMP1Mod1:** 'if the first stimulus has more intensity than the latter stimulus in the first modality then respond to the first stimulus otherwise do not respond', 20; 'if the first orientation is modality than the final orientation in the first modality then respond to the first orientation otherwise do not respond', 12; 'if the first stimulus is more intense than the last stimulus in the first modality then respond to the first stimulus otherwise do not respond', 11; 'if the first orientation has higher intensity than the first orientation in the first modality then respond to the first orientation otherwise do not respond', 8; 'focus only the first modality and opt for the first direction if it has greater strength than the second orientation otherwise do not respond', 7; **COMP2Mod1:** 'select the final stimulus if it is presented with higher intensity than the first stimulus in the first modality otherwise do not respond', 8; 'respond to the final stimulus if it is presented with greater strength than the first in the first modality otherwise do not respond', 7; 'focus only on the first modality and choose the the final orientation if it is stronger than the first direction otherwise do not respond', 4; 'opt for the final orientation if it has a greater period of time than the first orientation otherwise do not respond', 4; 'focus only on the first modality and choose the second direction if it is stronger than the first direction otherwise do not respond', 4; **COMP1Mod2:** 'pick the first direction if it has more intensity than the latter stimulus in the second modality otherwise do not respond', 25; 'select the initial stimulus if it is displayed with greater strength than the second stimulus otherwise do not respond', 20; 'opt for the initial stimulus if it is presented with greater strength than the final stimulus in the second modality otherwise do not respond', 7; 'if the initial stimulus is more intense than the final stimulus in the second modality then respond to the initial stimulus otherwise do not respond', 7; 'select the initial stimulus if it is stronger than the latter stimulus in the second modality otherwise do not respond', 6; **COMP2Mod2:** 'focus only on the second modality and choose the latter direction if it more intensely than the first direction otherwise do not respond', 34; 'focus on the final modality and choose the latter stimulus if it is stronger than the initial direction otherwise do not respond', 10; 'attend only to the second modality and select the final direction if it has more intensity than the first direction otherwise do not respond', 9; 'focus only on the second modality and pick the latter direction if it has more strength than the first direction otherwise do not respond', 7; 'attend only to the second modality and select the latter direction if it is stronger than the first direction otherwise do not respond', 6; **Dur1:** 'if the first stimulus is shorter than the final stimulus respond to the initial stimulus otherwise do not respond', 5; 'if the latter stimulus appears for more time than the first stimulus than the second stimulus otherwise do not respond', 5; 'go in the direction of the stimulus choose the stimulus stimulus', 4; 'if the latter stimulus appears for less time than the final stimulus respond to the initial stimulus otherwise do not respond', 4; 'if the duration of the first stimulus is greater than the duration of the first stimulus respond to the first stimulus otherwise do not respond', 4; **Dur2:** 'if the stimuli match respond in the direction of the stimulus otherwise do not respond', 11; 'choose the final direction if it lasts for more time than the first stimulus otherwise do not respond', 8; 'if the final direction is displayed for more time than the first direction respond to the final orientation otherwise do not respond', 7; 'choose the final direction choose the a', 6; 'choose the stimulus direction', 5; **MultiDur1:** 'pick the first direction if it has shorter shorter', 6; 'choose the first direction if it lasts for a longer a longer than the final direction otherwise do not respond', 4; 'if the initial stimulus has a duration which lasts than the final stimulus when combined across both modalities then select the first stimulus otherwise do not respond', 4; 'if the initial stimulus appears for a longer period of time than the second stimulus averaged across both modalities then choose that direction otherwise do not respond', 3; 'pick the first direction if it lasts for more time than the final direction otherwise do not respond', 3; **MultiDur2:** 'pick the final direction if it has a shorter duration than the initial direction combined over both modalities otherwise do not respond', 6; 'respond to the final direction if it lasts for a longer period of time than the first stimulus otherwise do not respond', 6; 'select the second orientation if it appears for longer than the first orientation in the second modality otherwise do not respond', 5; 'select the second orientation if it has a longer duration than the first orientation in the second modality otherwise do not respond', 4; 'select the second orientation if it lasts for longer than the first orientation in the second modality otherwise do not respond', 4; **AntiDur1:** 'respond to the initial stimulus if it is displayed for less time than the second stimulus otherwise do not respond', 5; 'respond to the initial orientation if it is displayed for less time than the second orientation otherwise do not respond', 5; 'respond to the first orientation if it lasts for longer of time than the final direction otherwise do not respond', 5; 'select the first stimulus if it has a shorter duration than the final stimulus otherwise do not respond', 5; 'respond to the first stimulus if it is shorter than the final stimulus otherwise do not respond', 4; **AntiDur2:** 'opt for the final direction if it is shorter than the final direction otherwise do not respond', 11; 'go in the second direction when it has presented intensity', 6; 'respond with the orientation orientation', 5; 'choose the latter direction if it is shorter than the final direction otherwise do not respond', 4; 'if the initial direction has less strength than the second direction then respond to the initial direction otherwise do not respond', 4; **AntiMultiDur1:** 'if the latter direction is shorter than the initial stimulus respond to the latter stimulus otherwise do not respond', 9; 'choose the initial stimulus if is more intense than the last stimulus otherwise do not respond', 6; 'if the first stimulus is less than the second stimulus respond than the first stimulus otherwise do not respond', 5; 'if the initial orientation is shorter than the second direction averaged over both modalities then respond to the initial direction otherwise do not respond', 5; 'choose the initial stimulus if it is presented than the second stimulus otherwise do not respond', 5; **AntiMultiDur2:** 'respond with least stimulus strength', 21; 'respond in the reverse direction if the stimuli are otherwise do not respond', 16; 'opt for the latter direction if it is presented with the initial direction otherwise do not respond', 9; 'pick the weakest direction if', 8; 'respond with least intensity', 6; **Dur1Mod1:** 'focus only on the first modality and opt for the initial direction if it has a greater duration than the final direction otherwise do not respond', 7; 'attend only to the first modality and pick the initial orientation if it has a duration duration than the second orientation otherwise do not respond', 7; 'focus on the first modality and opt for the initial stimulus if it is presented for longer than the final stimulus otherwise do not respond', 5; 'focus only on the first modality and opt for the first direction if it has a greater duration than the second direction otherwise do not respond', 5; 'attend only to the first modality and respond to the first stimulus has a greater duration than the second direction otherwise do not respond', 5; **Dur2Mod1:** 'choose the final stimulus if it has the first intensity than the initial direction otherwise do not respond', 9; 'select the second orientation if it is presented for longer than the first stimulus in the first modality otherwise do not respond', 9; 'attend to the first modality and respond to the final orientation if it lasts for a greater period of time than the first orientation otherwise do not respond', 7; 'select the second orientation if it is presented for longer than the first direction in the first modality otherwise do not respond', 6; 'choose the orientation with lowest in the second modality', 6; **Dur1Mod2:** 'go in the first direction if it has a greater duration than the latter stimulus in the second modality otherwise do not respond', 16; 'select the initial stimulus if it has a greater duration than the second stimulus in the second modality otherwise do not respond', 11; 'select the initial direction if it is displayed for more time than the final direction in the final modality otherwise do not respond', 11; 'select the initial orientation if it has a greater than the latter direction in the second modality otherwise do not respond', 10; 'select the initial direction if it has a greater duration than the latter direction in the second modality otherwise do not respond', 7; **Dur2Mod2:** 'attend to the first modality and respond to the second direction if it is displayed with time than the first direction otherwise do not respond', 9; 'pay attention only to the second modality and pick the final direction if it has a longer duration than the first direction otherwise do not respond', 7; 'attend only to the second modality and pick the final stimulus if it appears for a greater period of time than the initial stimulus otherwise do not respond', 6; 'select the second direction if it is displayed in greater strength than the first direction in the second modality otherwise do not respond', 6; 'attend to the first modality and respond to the second direction if it is displayed with less intensity than the first orientation otherwise do not respond', 4; **DMS:** 'if the stimuli match respond', 14; 'if the stimuli match respond in the displayed direction otherwise do not respond', 8; 'choose the most intense displayed', 8; 'if the duration of the same half of the display then respond to the first direction otherwise do not respond', 5; 'if the most intensely second displayed then are in the first orientation otherwise do not respond', 5; **DNMS:** 'if displayed directions are distinct select the same direction', 8; 'if the stimuli are mismatched go in direction of the second direction otherwise do not respond', 8; 'if the stimuli are mismatched go in the second direction otherwise do not respond', 5; 'choose the latter direction when it is presented for', 5; 'choose the latter direction when stimuli appear on different otherwise do not respond', 5; **DMC:** 'go in the orientation indicated by', 8; 'go in the orientation indicated', 6; 'if the first displayed directions are the same orientation otherwise do not respond', 4; 'pick the first direction if the first and stimuli are the same direction otherwise do not respond', 4; 'pick the first direction if the first and stimuli are the same half of the display otherwise do not respond', 4; **DNMC:** 'if the directions are go in the final direction otherwise do not respond', 8; 'go in the strongest direction if you are confident in your answer otherwise do not respond', 8; 'go in the direction of the orientation otherwise do not respond', 6; 'go in the strongest direction if you have high confidence otherwise do not respond', 6; 'go in the orientation displayed', 4

**Sensorimotor-RNN trained with Tasks Held Out and Language Production trained with Tasks Held Out: Go:** 'respond in the direction of the strongest stimulus strength', 12; 'respond in the direction of the stimulus strength', 12; 'respond in the direction of the stimulus displayed', 12; 'respond in the direction of the stimulus displayed the first modality', 10; 'respond in the direction of greatest displayed with greatest strength', 7; **AntiGo:** 'opt for the reverse of the orientation displayed', 17; 'opt for the opposite of the presented with', 11; 'pay attention to the first modality and respond to the opposite of the displayed direction', 7; 'opt for the opposite of the orientation displayed in the first modality', 6; 'opt for the reverse of the orientation presented in the first modality', 6; **RTGo:** 'respond to the direction direction at the modality at stimulus onset', 33; 'pay attention to the first modality and respond to the direction immediately', 23; 'immediately in the direction displayed in the first modality', 14; 'focus only on the first modality and choose the displayed direction at stimulus', 14; 'pay attention only to the second modality and choose the stimulus at stimulus onset', 10; **AntiRTGo:** 'respond with the opposite of the orientation at stimulus onset', 24; 'respond with the opposite of the stimulus in the first modality at stimulus onset', 19; 'respond with the opposite orientation at stimulus onset', 16; 'opt for the opposite of the stimulus immediately', 13; 'respond with the opposite of the stimulus in the second modality at stimulus onset', 13; **GoMod1:** 'select the orientation presented with lowest intensity', 17; 'respond in the direction presented with the strength', 17; 'select the direction in the first modality that is strongest', 11; 'pick the stimulus in the second modality with least intensity intensity', 9; 'select the stimulus stimulus select the first modality', 8; **AntiGoMod1:** 'choose the opposite of the stimulus in the first modality', 34; 'attend only to the stimulus in the first modality and respond in the opposite direction', 17; 'attend only to the first modality and pick the reverse direction', 13; 'opt for the opposite of the stimulus in the first modality', 9; 'opt for the opposite of the direction in the second modality', 9; **GoMod2:** 'attend only to the second modality and go in the reverse of the second modality', 24; 'choose the direction with largest strength in the second modality', 14; 'focus on the second modality and opt with the stimulus strength', 9; 'focus on the second modality and opt with the weakest stimulus strength', 8; 'choose the stimulus that appears with lowest intensity', 7; **AntiGoMod2:** 'respond in the opposite of the direction in the second modality', 17; 'pay attention to the second modality and select the opposite direction', 15; 'respond to the opposite of the stimulus displayed', 9; 'pay attention to the second modality and the opposite direction of the opposite stimulus', 8; 'pay attention to the second modality and the opposite direction', 8; **RTGoMod1:** 'select the direction in the direction in the first modality', 27; 'select the direction that appears in the first modality', 10; 'select the orientation that in the first modality at stimulus onset', 9; 'choose the first direction in the first modality', 9; 'select the orientation displayed in the first orientation at stimulus onset', 8; **AntiRTGoMod1:** 'attend only to the first modality and choose the reverse of the direction displayed at stimulus onset', 24; 'focus on the first modality and choose the displayed direction at stimulus onset', 12; 'attend only on the first modality and go in the opposite of the displayed direction', 6; 'attend only to the first modality and pick the reverse direction at stimulus onset', 3; 'pay attention only to the first modality and choose the reverse direction at stimulus onset', 3; **RTGoMod2:** 'focus only on the second modality and go in the direction at stimulus onset', 36; 'choose the direction that appears in the second modality at stimulus onset', 23; 'choose the direction that appears in the second modality', 19; 'choose the direction in the second modality that appears on stimulus modality', 19; 'select the stimulus that appears in the second modality at stimulus onset', 17; **AntiRTGoMod2:** 'attend to the second modality and choose the reverse of the orientation at stimulus onset', 29; 'opt for the reverse of the orientation in the second modality at stimulus onset', 17; 'attend to the second modality and respond to the reverse direction immediately', 16; 'attend to the second modality and respond in the reverse of the stimulus immediately', 14; 'opt for the reverse of the orientation in the second modality immediately respond onset', 8; **DM:** 'respond to the the presented stimulus with the same strength', 28; 'respond in the direction of highest stimulus', 21; 'choose the direction with highest intensity', 12; 'respond to the stimulus presented with highest intensity', 8; 'choose the direction representing the highest stimulus strength', 5; **AntiDM:** 'respond in the direction of the presented with least strength if you are confident otherwise do not respond', 11; 'respond in the direction of the presented with least strength otherwise do not respond', 6; 'respond to the stimulus with lowest strength', 4; 'respond to the stimulus with lowest strength if you are sure about your answer otherwise do not respond', 4; 'respond in the direction of the presented with least strength if you are sure about your decision otherwise do not respond', 4; **MultiDM:** 'select the direction of the stimulus', 19; 'select the direction of the stimulus presented over both modalities is', 7; 'select the direction of the stimulus presented over both modalities', 6; 'select the direction of the stimulus with', 5; 'select the direction of the stimulus with maximal strength', 5; **AntiMultiDM:** 'select the orientation presented with minimal strength over both modalities', 10; 'pick the stimulus with the stimuli strength', 8; 'pick the stimulus in the first stimulus', 6; 'select the orientation with minimal intensity', 5; 'select the orientation with lowest intensity over both modalities', 4; **DMMod1:** 'pick the direction that appears in the first modality', 19; 'attend only to the stimulus in the first modality which has maximal strength', 9; 'select the orientation that appears with maximal intensity', 7; 'attend to the stimulus in the first modality and respond in the stimulus', 5; 'attend to the stimulus in the first modality and respond in the direction', 5; **AntiDMMod1:** 'go in the direction of the stimulus presented with least strength', 25; 'select the orientation with lowest in the first modality', 25; 'focus on the first modality and select the weakest direction', 17; 'attend only to the first modality and choose the weakest direction', 14; 'focus on the first modality and select the weakest stimulus', 12; **DMMod2:** 'choose the direction with highest intensity in the second modality', 31; 'focus only on the second modality and pick the stimulus with highest intensity', 15; 'attend to the second modality and select the direction with highest intensity', 14; 'attend only to the second modality and pick the reverse with lowest strength', 14; 'focus on the second modality and pick the weakest direction', 9; **AntiDMMod2:** 'focus the stimulus in the second modality that has the strength', 13; 'pick the stimulus in the second modality that has minimal strength', 10; 'pick the stimulus in the second that that has lowest strength', 6; 'pick the stimulus in the second modality that has lowest strength', 5; 'pick the stimulus in the second modality that is presented with least strength', 5; **ConDM:** 'if the first stimulus is more intense than the second stimulus respond to the first stimulus otherwise do not respond', 9; 'go in the direction with weakest average intensity between two stimuli', 7; 'respond to the initial stimulus if it is displayed with less intensity than the second stimulus otherwise do not respond', 4; 'go in the direction with largest with highest intensity', 3; 'go in the direction with largest average intensity between two stimuli', 2; **ConAntiDM:** 'respond to the stimulus presented with lowest intensity', 16; 'choose the stimulus with weakest strength', 12; 'respond to the stimulus with lowest strength', 6; 'choose the stimulus with has strength', 5; 'respond to the stimulus presented with greatest strength', 4; **COMP1:** 'if the stimuli are in the second of the second direction that intensity otherwise do not respond', 15; 'pick the first direction if it has more intensity than the final stimulus otherwise do not respond', 13; 'pick the first direction if it has greatest strength than the final direction otherwise do not respond', 7; 'pick the first stimulus is more intense than the final stimulus otherwise do not respond', 6; 'respond if the stimuli are presented with greater intensity in the second direction otherwise do not respond', 5; **COMP2:** 'respond to the second stimulus if it is presented with greater intensity than the first stimulus', 22; 'respond in the second direction if it has greater strength than the first direction otherwise do not respond', 8; 'respond in the second direction if it has greatest strength than the first direction otherwise do not respond', 7; 'when the final stimulus is presented with higher intensity than the initial direction in the first direction otherwise do not respond', 6; 'when the latter direction when combined the higher value than the first direction otherwise do not respond', 5; **MultiCOMP1:** 'when the initial stimulus has higher value than the final direction respond in the initial direction otherwise do not respond', 27; 'if the first stimulus has greater strength than the final stimulus respond in the first direction otherwise do not respond', 16; 'if the initial direction has higher intensity than the second direction when considered over both modalities otherwise do not respond', 15; 'respond to the initial direction if it is presented with greater strength than the final direction otherwise do not respond', 9; 'when the initial stimulus has higher intensity than the final stimulus in the initial direction otherwise do not respond', 8; **MultiCOMP2:** 'if the final stimulus is presented with higher intensity than the initial stimulus in the first direction otherwise do not respond', 12; 'if the second stimulus has greater strength than the first stimulus averaged over modalities than the second stimulus otherwise do not respond', 10; 'if the final stimulus is weaker averaged over both modalities than the initial stimulus then respond to the final stimulus otherwise do not respond', 9; 'if the final stimulus is presented with higher intensity than the initial orientation in the first direction otherwise do not respond', 6; 'if the final stimulus is presented with higher intensity than the initial stimulus in the final direction otherwise do not respond', 6; **AntiCOMP1:** 'pick the initial stimulus if it is weaker than the final stimulus otherwise do not respond', 12; 'if the first stimulus has less intensity than the latter stimulus select the first stimulus pick do not respond', 8; 'if the first stimulus is less intense than the latter stimulus then the the first stimulus otherwise do not respond', 8; 'select the first stimulus if it is weaker than the final direction select the second direction otherwise do not respond', 8; 'respond to the initial direction if it is shorter than the second direction otherwise do not respond', 8; **AntiCOMP2:** 'if the final stimulus is weaker than the initial stimulus combined over both modalities then select the final stimulus otherwise do not respond', 17; 'if the latter direction is displayed with less intensity combined over both modalities than the first direction then choose the latter direction otherwise do not respond', 10; 'if the latter direction is displayed less intensity averaged over both modalities than the first direction then choose the latter direction otherwise do not respond', 9; 'select the latter orientation if it has less overall strength over both modalities than the initial orientation otherwise do not respond', 6; 'if the latter direction is displayed with less intensity than the first orientation than the first orientation then do not respond', 5; **AntiMultiCOMP1:** 'if the initial direction has less strength combined over both modalities than the initial direction then respond to the initial direction otherwise do not respond', 11; 'if the initial direction is displayed with less intensity than the latter direction then select the latter stimulus otherwise do not respond', 7; 'if the initial stimulus is presented with intensity intensity than the latter stimulus then select the latter stimulus otherwise do not respond', 7; 'respond to the initial direction if it is weaker than the final direction averaged over modalities otherwise do not respond', 6; 'if the first direction is displayed less strength averaged over both modalities to the final direction then select the first direction otherwise do not respond', 6; **AntiMultiCOMP2:** 'if the final orientation is presented with less intensity than the first direction averaged averaged over modalities than the final orientation otherwise do not respond', 4; 'if the final orientation is presented with less intensity than the first direction then select the final direction otherwise do not respond', 3; 'respond to the second direction if it is weaker than the initial direction otherwise do not respond', 3; 'respond to the second direction if it is weaker than the first stimulus otherwise do not respond', 3; 'if the final direction is weaker averaged over both initial the initial direction then select the final direction otherwise do not respond', 3; **COMP1Mod1:** 'if the first stimulus is greater than the second stimulus in the first modality then respond to the second stimulus otherwise do not respond', 12; 'go in the first direction if it is greater with greater strength than the latter stimulus otherwise do not respond', 9; 'select the first stimulus if it is presented with greater strength than the final stimulus in the first modality otherwise do not respond', 4; 'focus only on the first modality and opt for the first direction if it is more than the second direction otherwise do not respond', 4; 'go in the first direction if it is stronger than the latter stimulus in the first modality otherwise do not respond', 4; **COMP2Mod1:** 'respond to the second stimulus if it is presented with greater strength than the first stimulus in the second modality otherwise do not respond', 15; 'pay attention only to the first modality and select the latter orientation if it has greater intensity than the first orientation otherwise do not respond', 9; 'respond to the second stimulus if it is displayed for a greater period of time than the first stimulus in the first modality otherwise do not respond', 9; 'pay attention only to the first modality and select the latter orientation if it has greater intensity than the the orientation otherwise do not respond', 6; 'if the final stimulus is presented than the first stimulus than the final stimulus otherwise do not respond', 5; **COMP1Mod2:** 'pick the initial stimulus if it is more second than the second direction otherwise do not respond', 19; 'go in the initial direction if it has greater strength than the final direction in the second modality otherwise do not respond', 15; 'respond to the initial direction if it is stronger than the second direction in the first modality otherwise do not respond', 13; 'respond to the initial stimulus if it has higher intensity than the final stimulus respond to the second modality otherwise do not respond', 11; 'attend only to the second modality and pick the initial direction if it has a greater than the final direction otherwise do not respond', 10; **COMP2Mod2:** 'focus only on the second modality and opt for the final stimulus if it than the first stimulus otherwise do not respond', 7; 'focus on the second modality and select the the displayed direction if is presented on the first direction otherwise do not respond', 7; 'focus on the second modality and select the the displayed direction if is presented on more intensely than the first direction otherwise do not respond', 6; 'focus only on the second modality and choose the first orientation if it orientation presented with more intensity than the second direction otherwise do not respond', 6; 'focus only on the second modality and opt for the final stimulus if it is presented than the first stimulus otherwise do not respond', 5; **Dur1:** 'if the first stimulus appears for more time than the second stimulus choose the first stimulus then choose the first stimulus otherwise do not respond', 7; 'choose the first direction if it lasts for a greater period of time than the final direction otherwise do not respond', 6; 'if the second direction has a shorter duration than the initial direction select the initial direction otherwise do not respond', 4; 'if the second direction is shorter than the initial direction pick the initial direction otherwise do not respond', 4; 'if the first stimulus appears for more time than the second stimulus choose the first stimulus choose the first stimulus otherwise do not respond', 4; **Dur2:** 'if the directions match select the displayed direction otherwise do not respond', 32; 'choose the stimulus orientation', 14; 'choose the stimulus orientation in the first modalities', 6; 'when the two displayed directions are go in the direction otherwise do not respond', 6; 'if the directions match select the final displayed otherwise do not respond', 5; **MultiDur1:** 'attend only to the first modality and if the first stimulus lasts for longer than the second stimulus respond to the latter orientation otherwise do not respond', 5; 'pick the stimuli direction if it is shorter than the final direction otherwise do not respond', 5; 'if the initial stimulus lasts for a longer period of time than the latter orientation in the second modality then respond to the first orientation otherwise do not respond', 3; 'go in the direction of the first stimulus if you', 3; 'go in the direction of the first stimulus if is lasts than the second stimulus otherwise do not respond', 3; **MultiDur2:** 'if the first stimulus has a shorter duration than the latter stimulus respond to the first stimulus otherwise do not respond', 7; 'respond to the latter direction if it displayed for a greater period of time than the latter direction otherwise do not respond to the initial direction otherwise do not', 5; 'choose the second orientation if it has a longer duration than the initial orientation considered across both modalities otherwise do not respond', 4; 'attend to the second modality and select the latter stimulus if it is displayed for a greater period of time than the initial stimulus otherwise do not respond', 3; 'if the latter direction is displayed for a time time than the initial direction then respond to the latter direction otherwise do not respond', 3; **AntiDur1:** 'respond to the initial direction if it is displayed for less time than the second direction otherwise do not respond', 21; 'respond to the initial direction if it is displayed for less time than the latter direction otherwise do not respond', 19; 'if the first stimulus is shorter than the final stimulus respond to the first stimulus otherwise do not respond', 15; 'respond to the initial direction if it has a shorter duration than the second direction otherwise do not respond', 10; 'pick the first direction if it has less duration than the final direction otherwise do not respond', 9; **AntiDur2:** 'go in the direction displayed direction', 11; 'go in the displayed direction', 9; 'choose the second orientation if it is displayed for', 5; 'if the initial stimulus is presented with less strength than the latter stimulus than the latter stimulus otherwise do not respond', 5; 'if the initial stimulus is presented with less time than the second stimulus respond to the latter stimulus otherwise do not respond', 4; **AntiMultiDur1:** 'pick the initial stimulus if it is displayed for less time than the final stimulus otherwise do not respond', 21; 'choose the initial orientation if it is presented with greater intensity than the second orientation otherwise do not respond', 8; 'respond to the first stimulus if it has a shorter duration than the second stimulus otherwise do not respond', 7; 'choose the initial orientation if it is greater than the second orientation otherwise do not respond', 7; 'if the first stimulus has less duration than the latter stimulus respond to the first stimulus otherwise do not respond', 4; **AntiMultiDur2:** 'respond in with stimulus with', 10; 'respond in the latter direction when stimuli are mismatched otherwise do not respond', 10; 'respond in the latter direction if stimuli are mismatched otherwise do not respond', 8; 'select the stimulus with', 6; 'pick the weakest direction if you are confident for your answer otherwise do not respond', 5; **Dur1Mod1:** 'pay attention to the first modality and pick the first direction if it has a longer duration than the initial direction otherwise do not respond', 26; 'choose the first orientation if it is orientation which than the final orientation in the first modality then respond to the first stimulus otherwise do not respond', 22; 'pay attention to the first modality and pick the first stimulus if it has a longer duration than the initial direction otherwise do not respond', 15; 'choose the first direction if it is displayed for a greater period of time than the final direction in the first modality otherwise do not respond', 11; 'focus only on the first modality and select the initial orientation if it has longer longer than the first direction otherwise do not respond', 8; **Dur2Mod1:** 'choose the direction of the stimulus go in the second modality', 18; 'focus on the second modality and select the direction displayed', 11; 'if the final orientation is displayed for longer than the initial orientation in the first orientation otherwise do not respond', 10; 'if the final stimulus is shorter than the initial direction then respond do not respond', 6; 'if the final orientation is greater than the first orientation is greater than the first orientation otherwise do not respond', 4; **Dur1Mod2:** 'select the initial orientation if it has a longer duration than the second orientation otherwise do not respond', 21; 'choose the initial orientation if it appears for longer than the second orientation in the first modality otherwise do not respond', 15; 'select the initial orientation if it has a greater duration than the second orientation otherwise do not respond', 13; 'select the initial orientation if it appears for longer than the second orientation in the first modality otherwise do not respond', 11; 'select the initial orientation if is is displayed for more than the second direction in the second modality otherwise do not respond', 10; **Dur2Mod2:** 'select the second direction if it is displayed for more than the initial direction in the first modality otherwise do not respond', 14; 'select the second direction if it is displayed for more than the first direction in the first modality otherwise do not respond', 10; 'select the second direction if it is displayed for a greater of the first direction in the first modality otherwise do not respond', 5; 'pay attention only to the second modality and choose the strongest direction', 4; 'attend only to the first modality and choose the final direction if it is longer than the initial direction otherwise do not respond', 4; **DMS:** 'choose the most direction if the strongest stimulus if', 8; 'choose the most direction if the displayed on do not respond', 7; 'if the displayed directions are distinct halves select the latter direction otherwise do not respond', 5; 'if the stimuli are presented in the same direction of the display otherwise do not respond', 4; 'if the duration of the first orientation is longer than the second direction respond to the second modality otherwise do not respond', 4; **DNMS:** 'if the first direction has the initial direction select the second direction otherwise do not respond', 4; 'opt the the final direction if it has', 4; 'select the orientation when direction when the stimuli match otherwise do not otherwise do not respond', 3; 'if the first direction has the initial direction when select the second direction otherwise do not respond', 3; 'if the first direction has the displayed directions select the second direction when the latter direction otherwise do not respond', 3; **DMC:** 'go in the orientation direction', 12; 'pick the first direction if it appears for the second direction otherwise do not respond', 11; 'pick the first direction if it appears for more time than the second direction otherwise do not respond', 5; 'select the first direction if the first match otherwise do not respond', 4; 'if the first direction is is displayed for less intense than the second direction then choose the first direction otherwise do not respond', 3; **DNMC:** 'choose the displayed direction if both directions are distinct', 8; 'when the displayed directions are distinct select the final direction otherwise do not respond', 8; 'when the with stimulus in the latter stimulus otherwise do not respond', 5; 'if displayed directions are distinct then select the final direction otherwise do not respond', 4; 'choose the last direction if both directions are distinct', 4

#### 16 Example Task Trials

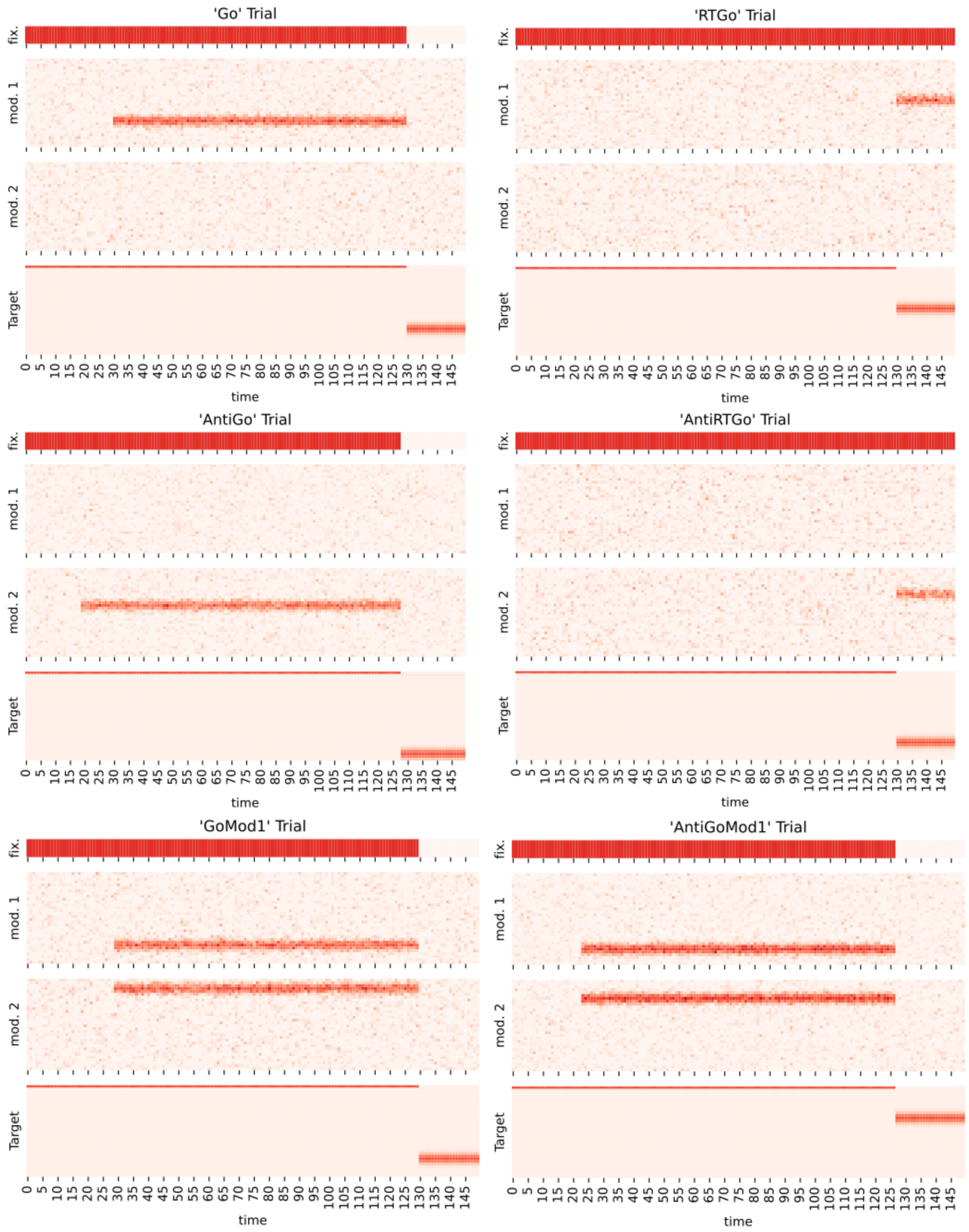

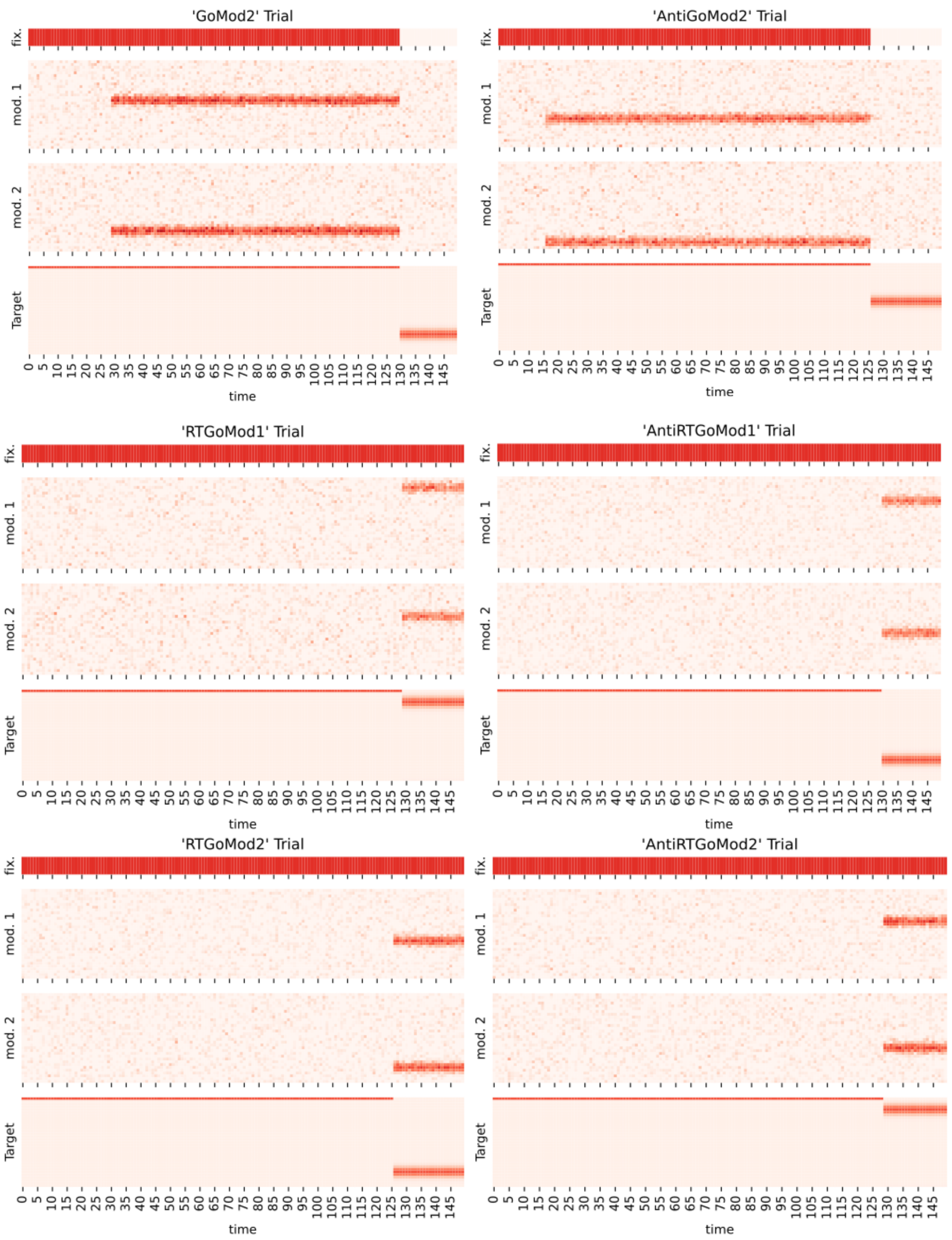
'GoMod2' Trial
'AntiGoMod2' Trial
'RTGoMod1' Trial
'AntiRTGoMod1' Trial
'RTGoMod2' Trial
'AntiRTGoMod2' Trial
fix.
mod. 1
mod. 2
Target
time
0 5 10 15 20 25 30 35 40 45 50 55 60 65 70 75 80 85 90 95 100 105 110 115 120 125 130 135 140 145

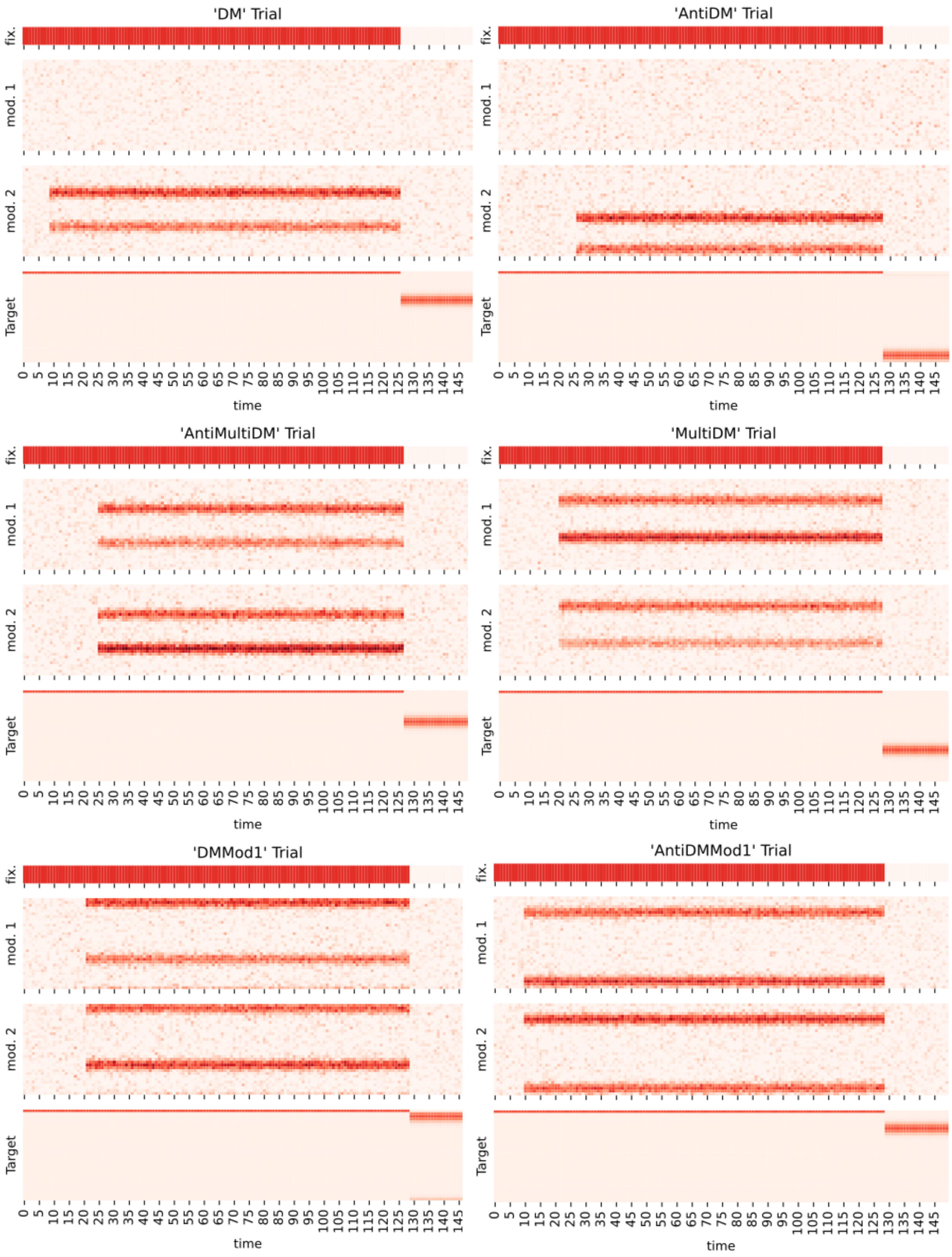
'DM' Trial
'AntiDM' Trial
'AntiMultiDM' Trial
'MultiDM' Trial
'DMMod1' Trial
'AntiDMMod1' Trial
fix.
mod. 1
mod. 2
Target
time
0 5 10 15 20 25 30 35 40 45 50 55 60 65 70 75 80 85 90 95 100 105 110 115 120 125 130 135 140 145

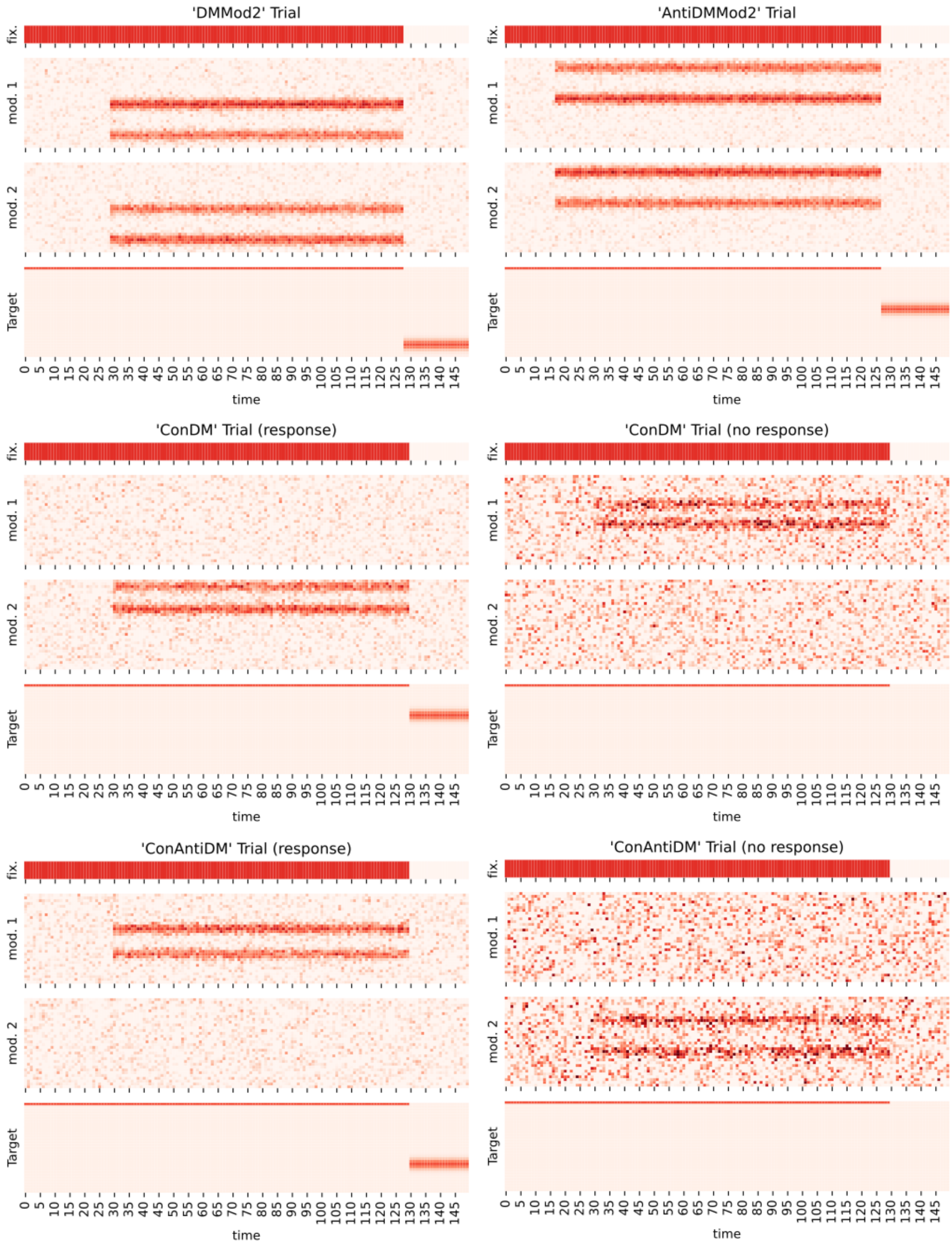
'DMMod2' Trial
'AntiDMMod2' Trial
'ConDM' Trial (response)
'ConDM' Trial (no response)
'ConAntiDM' Trial (response)
'ConAntiDM' Trial (no response)
fix.
mod. 1
mod. 2
Target
0 5 10 15 20 25 30 35 40 45 50 55 60 65 70 75 80 85 90 95 100 105 110 115 120 125 130 135 140 145
time

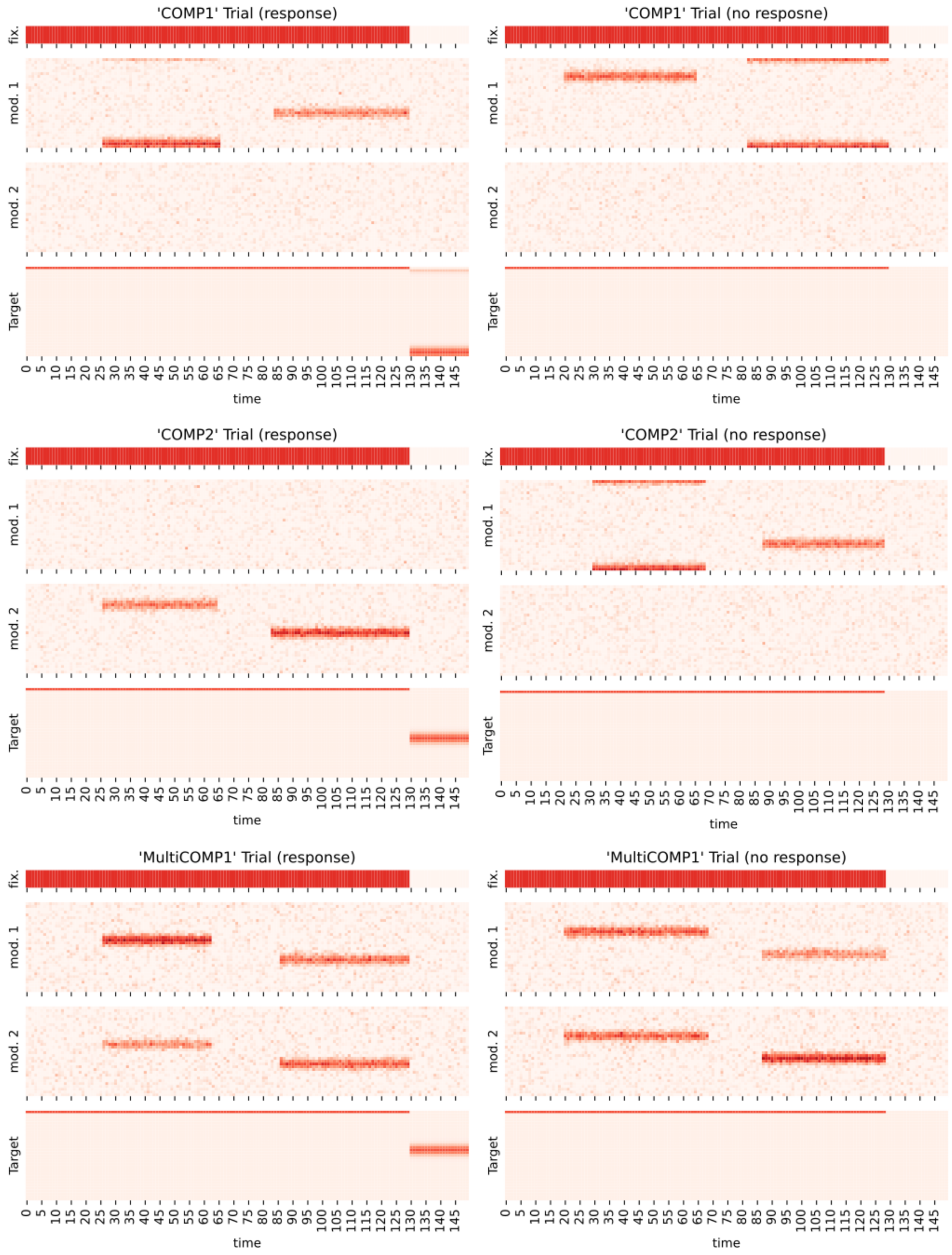
'COMP1' Trial (response)
'COMP1' Trial (no resposne)
'COMP2' Trial (response)
'COMP2' Trial (no response)
'MultiCOMP1' Trial (response)
'MultiCOMP1' Trial (no response)
fix.
mod. 1
mod. 2
Target
0 5 10 15 20 25 30 35 40 45 50 55 60 65 70 75 80 85 90 95 100 105 110 115 120 125 130 135 140 145
time

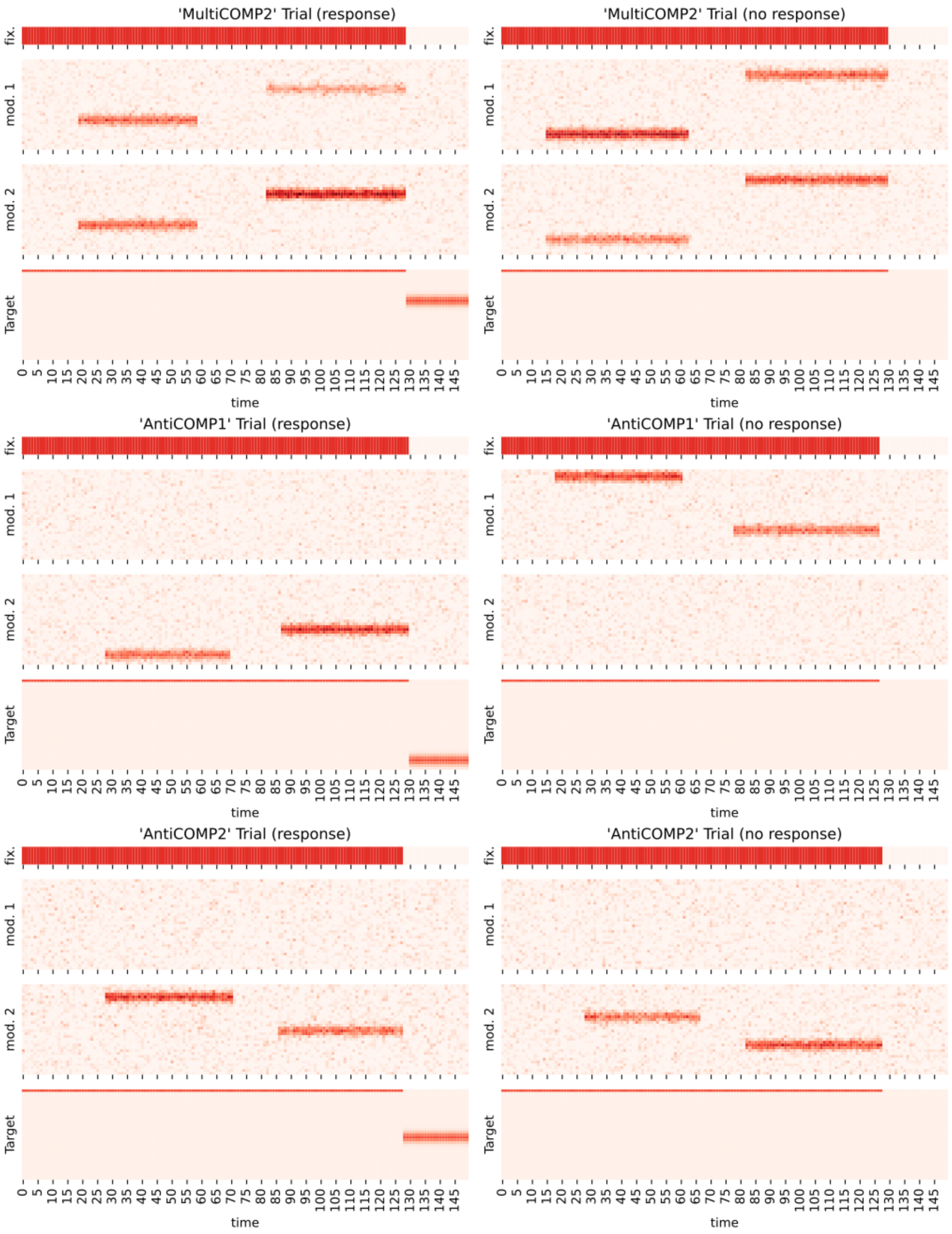
'MultiCOMP2' Trial (response)
'MultiCOMP2' Trial (no response)
'AntiCOMP1' Trial (response)
'AntiCOMP1' Trial (no response)
'AntiCOMP2' Trial (response)
'AntiCOMP2' Trial (no response)
fix.
mod. 1
mod. 2
Target
time
0 5 10 15 20 25 30 35 40 45 50 55 60 65 70 75 80 85 90 95 100 105 110 115 120 125 130 135 140 145

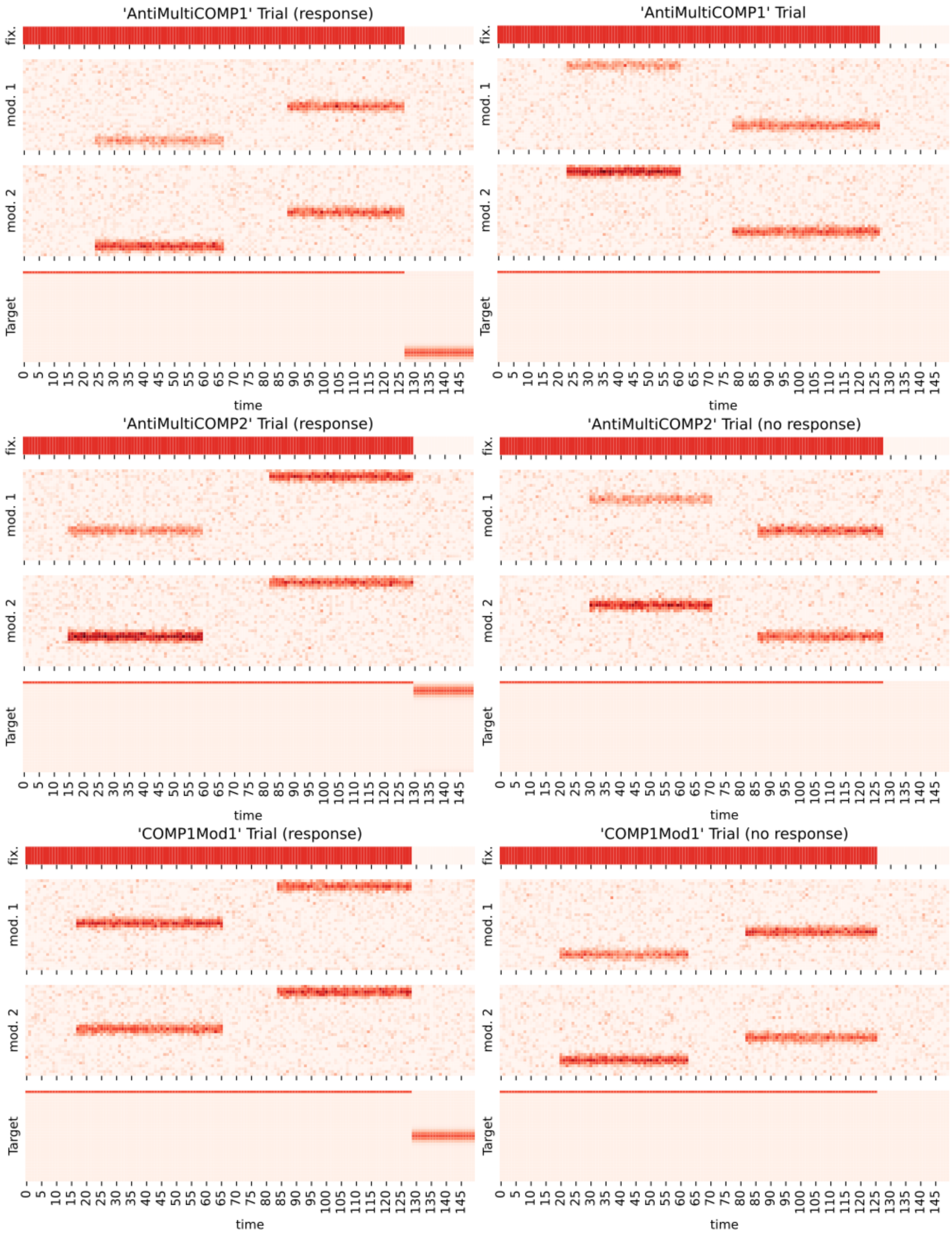
'AntiMultiCOMP1' Trial (response)
fix.
mod. 1
mod. 2
Target
0 5 10 15 20 25 30 35 40 45 50 55 60 65 70 75 80 85 90 95 100 105 110 115 120 125 130 135 140 145
time
'AntiMultiCOMP1' Trial
fix.
mod. 1
mod. 2
Target
0 5 10 15 20 25 30 35 40 45 50 55 60 65 70 75 80 85 90 95 100 105 110 115 120 125 130 135 140 145
time
'AntiMultiCOMP2' Trial (response)
fix.
mod. 1
mod. 2
Target
0 5 10 15 20 25 30 35 40 45 50 55 60 65 70 75 80 85 90 95 100 105 110 115 120 125 130 135 140 145
time
'AntiMultiCOMP2' Trial (no response)
fix.
mod. 1
mod. 2
Target
0 5 10 15 20 25 30 35 40 45 50 55 60 65 70 75 80 85 90 95 100 105 110 115 120 125 130 135 140 145
time
'COMP1Mod1' Trial (response)
fix.
mod. 1
mod. 2
Target
0 5 10 15 20 25 30 35 40 45 50 55 60 65 70 75 80 85 90 95 100 105 110 115 120 125 130 135 140 145
time
'COMP1Mod1' Trial (no response)
fix.
mod. 1
mod. 2
Target
0 5 10 15 20 25 30 35 40 45 50 55 60 65 70 75 80 85 90 95 100 105 110 115 120 125 130 135 140 145
time

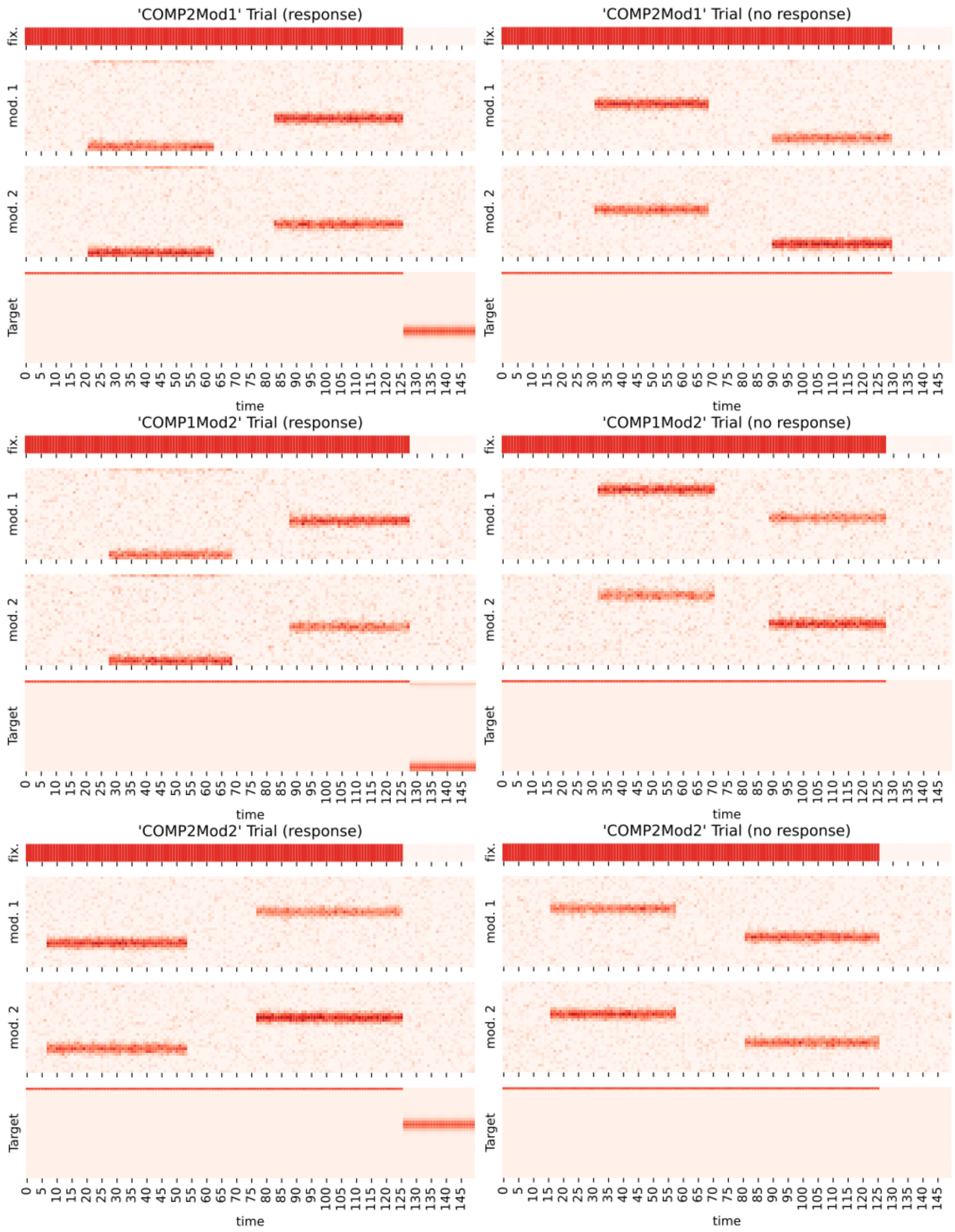
'COMP2Mod1' Trial (response)
'COMP2Mod1' Trial (no response)
'COMP1Mod2' Trial (response)
'COMP1Mod2' Trial (no response)
'COMP2Mod2' Trial (response)
'COMP2Mod2' Trial (no response)
fix.
mod. 1
mod. 2
Target
time
0 5 10 15 20 25 30 35 40 45 50 55 60 65 70 75 80 85 90 95 100 105 110 115 120 125 130 135 140 145

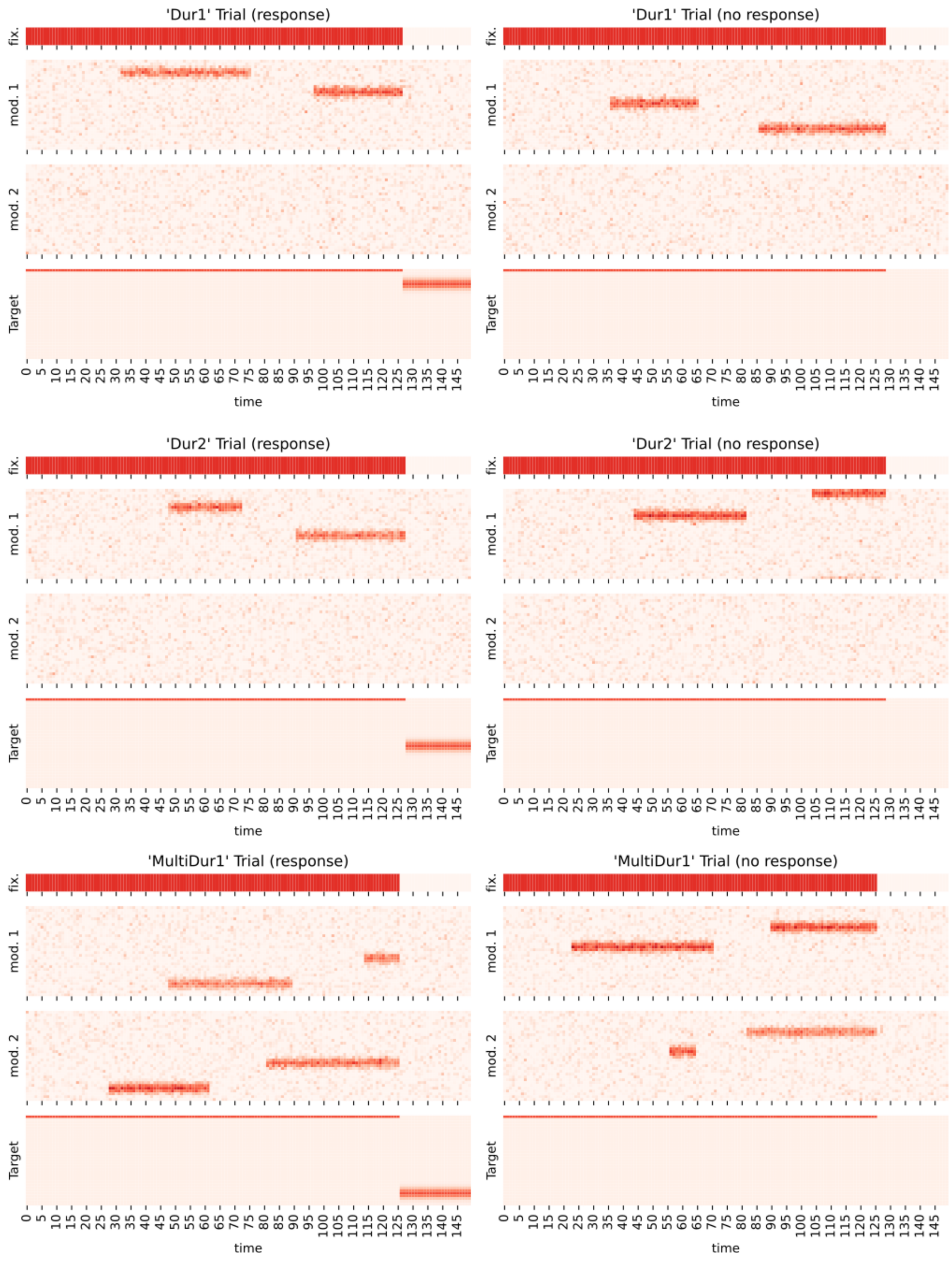
'Dur1' Trial (response)
'Dur1' Trial (no response)
'Dur2' Trial (response)
'Dur2' Trial (no response)
'MultiDur1' Trial (response)
'MultiDur1' Trial (no response)
fix.
mod. 1
mod. 2
Target
time

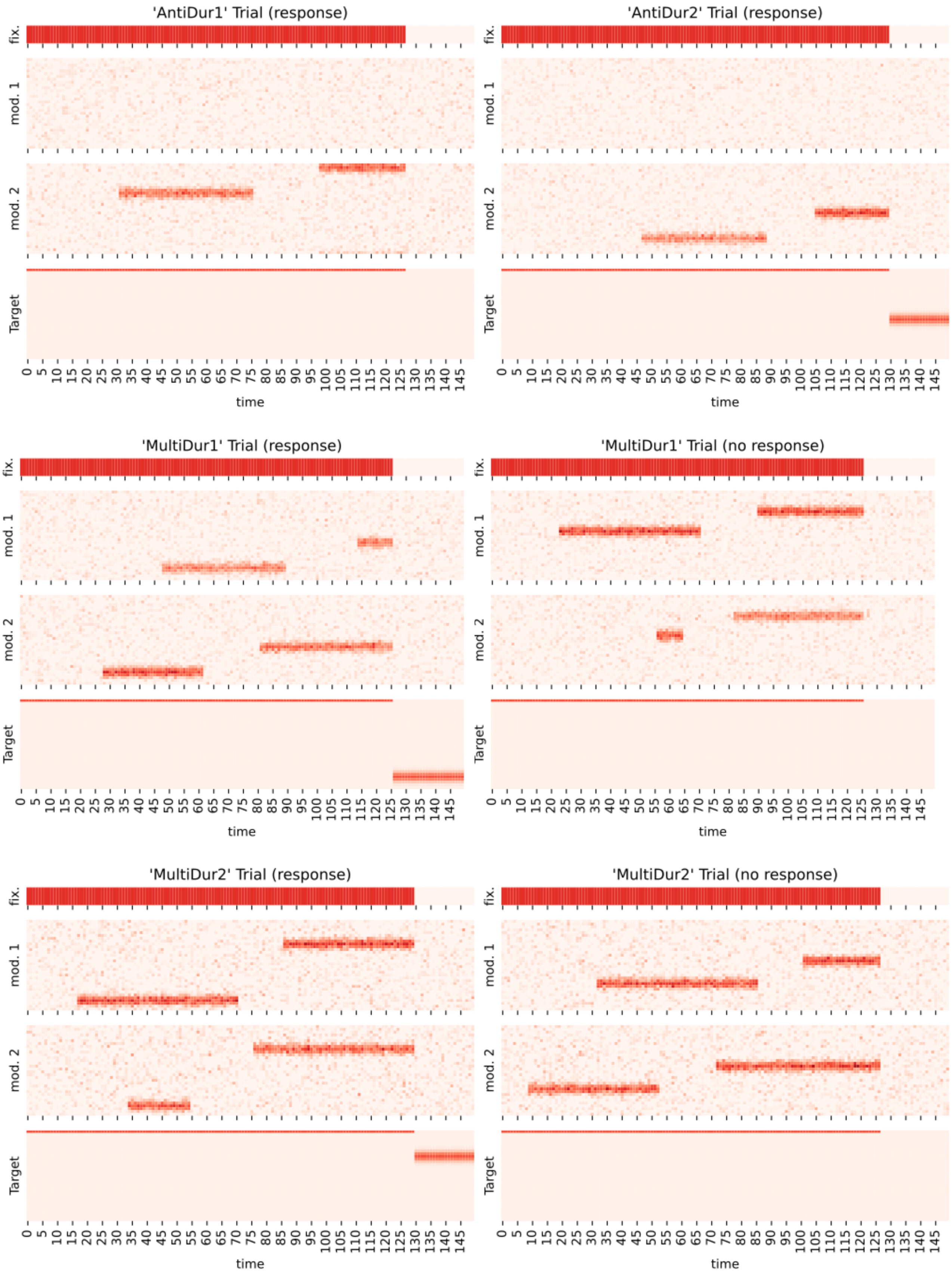
'AntiDur1' Trial (response)
'AntiDur2' Trial (response)
'MultiDur1' Trial (response)
'MultiDur1' Trial (no response)
'MultiDur2' Trial (response)
'MultiDur2' Trial (no response)
fix.
mod. 1
mod. 2
Target
time
0 5 10 15 20 25 30 35 40 45 50 55 60 65 70 75 80 85 90 95 100 105 110 115 120 125 130 135 140 145

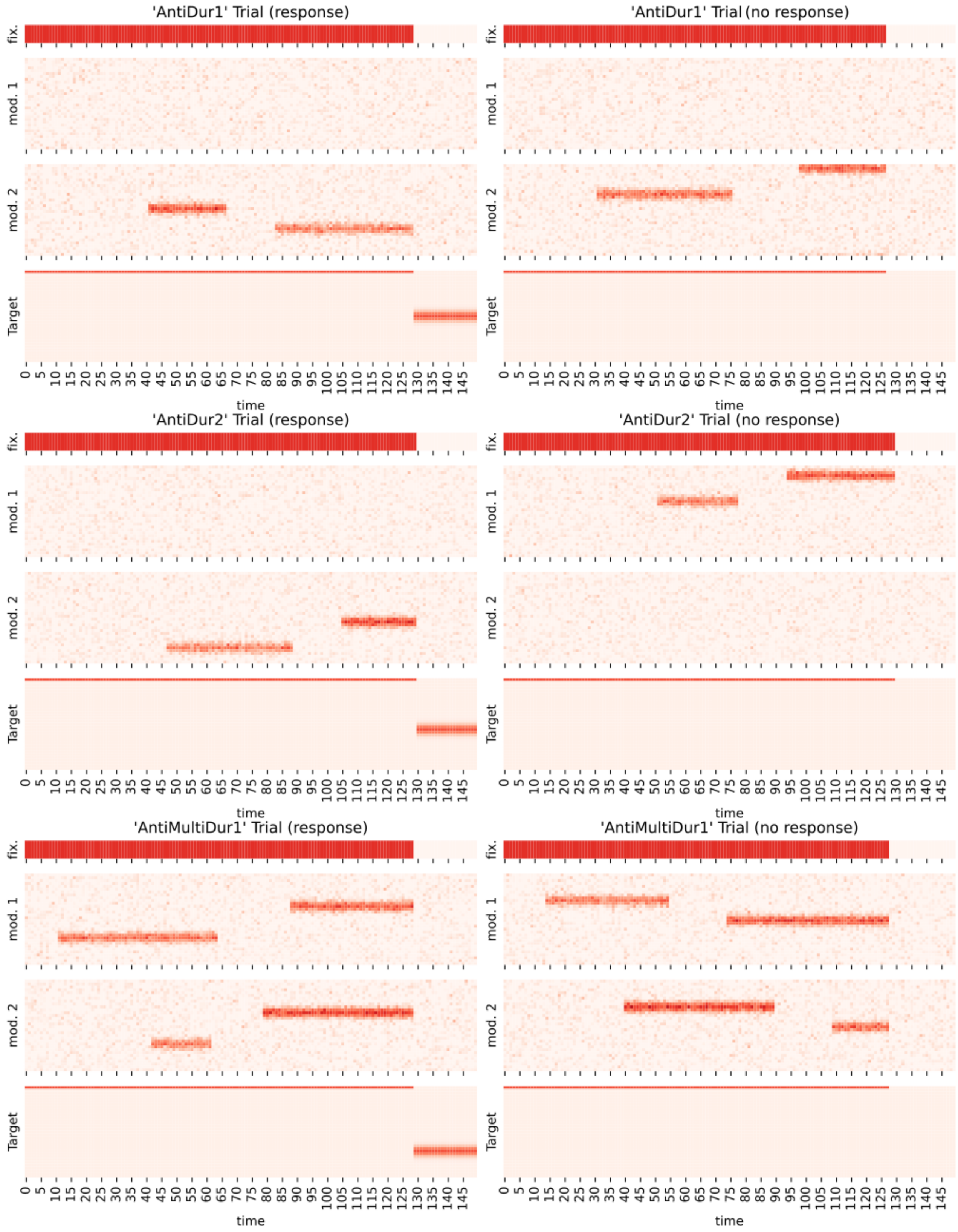
'AntiDur1' Trial (response)
'AntiDur1' Trial (no response)
'AntiDur2' Trial (response)
'AntiDur2' Trial (no response)
'AntiMultiDur1' Trial (response)
'AntiMultiDur1' Trial (no response)
fix.
mod. 1
mod. 2
Target
time

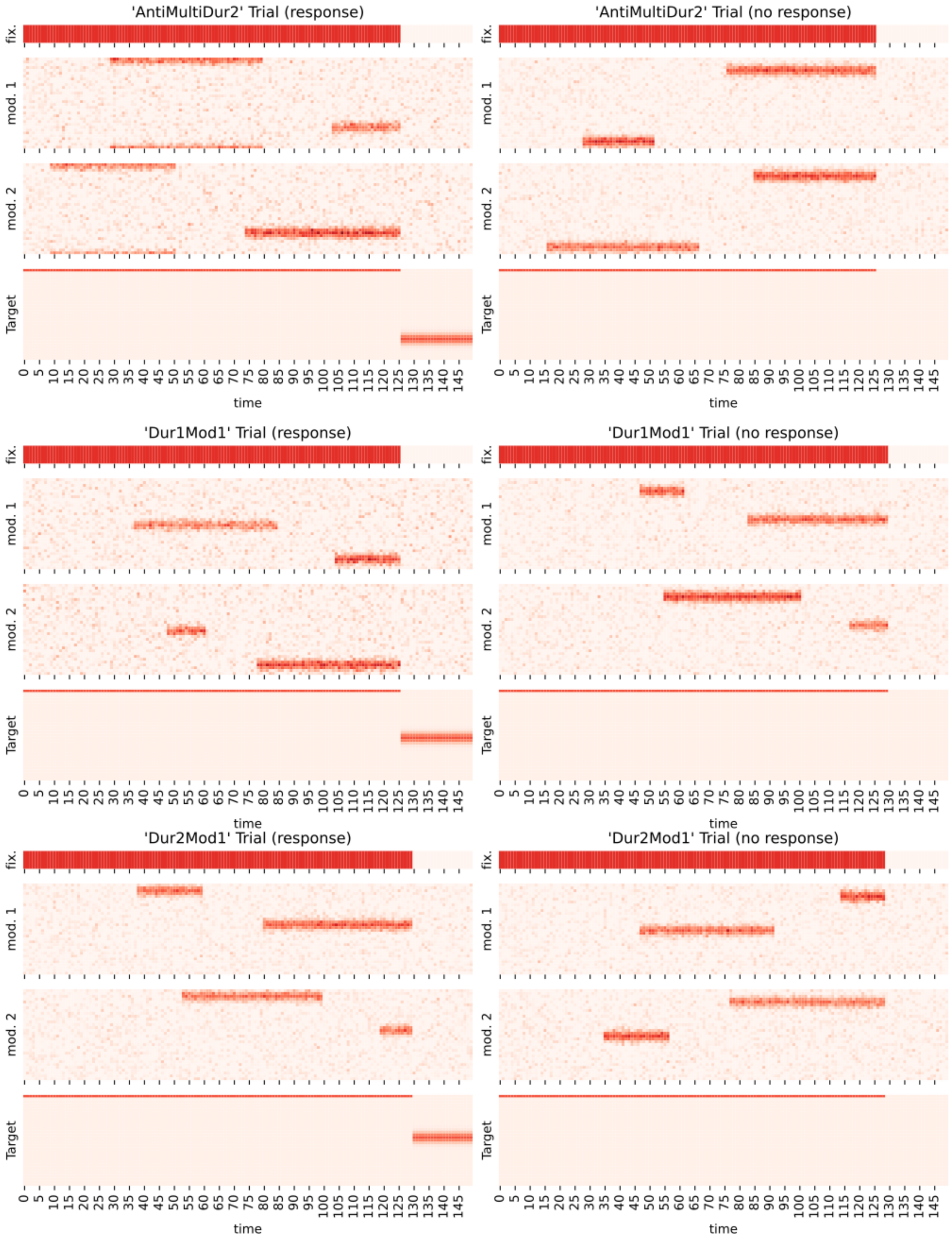
'AntiMultiDur2' Trial (response)
'AntiMultiDur2' Trial (no response)
'Dur1Mod1' Trial (response)
'Dur1Mod1' Trial (no response)
'Dur2Mod1' Trial (response)
'Dur2Mod1' Trial (no response)
fix.
mod. 1
mod. 2
Target
time
0 5 10 15 20 25 30 35 40 45 50 55 60 65 70 75 80 85 90 95 100 105 110 115 120 125 130 135 140 145

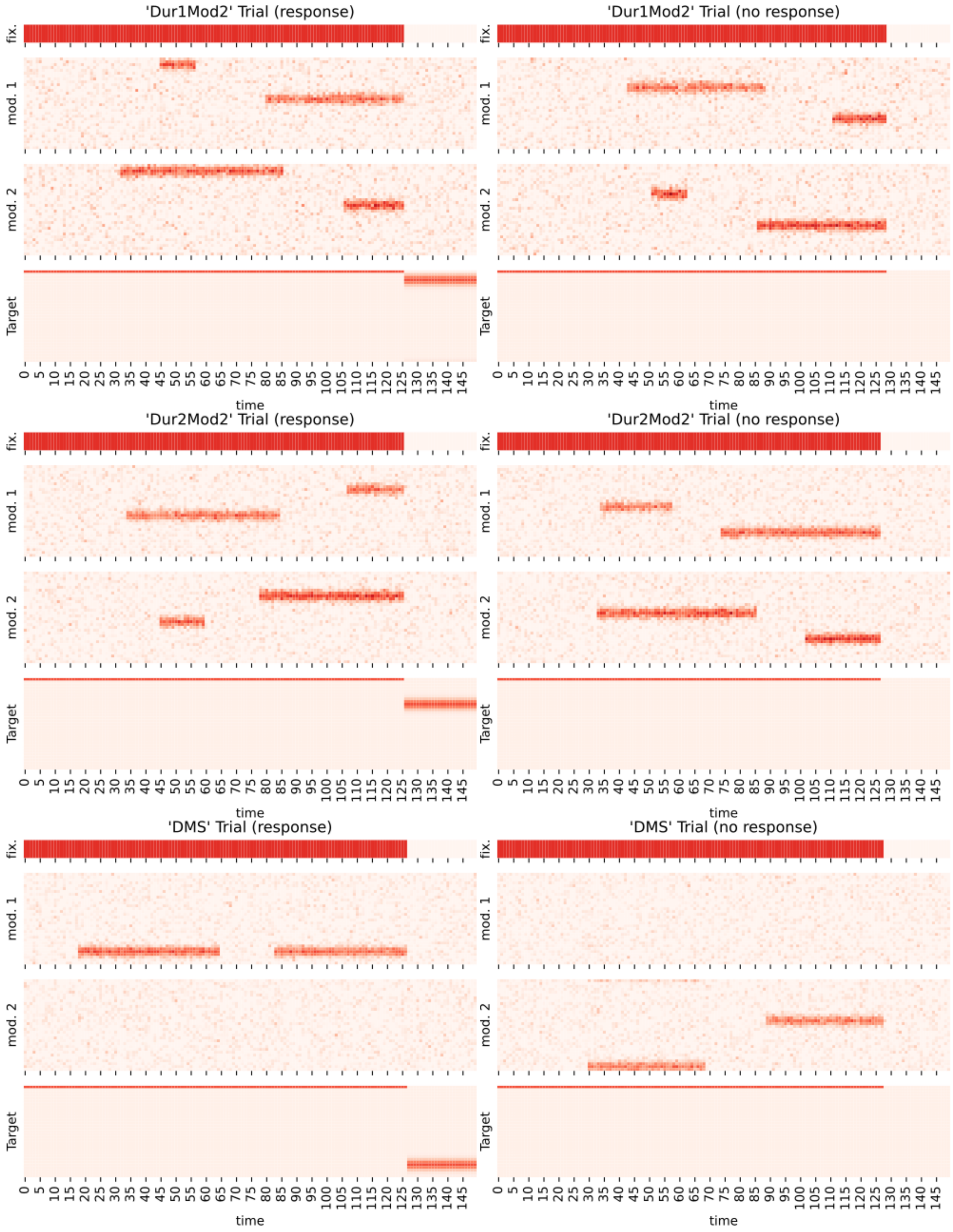
'Dur1Mod2' Trial (response)
'Dur1Mod2' Trial (no response)
'Dur2Mod2' Trial (response)
'Dur2Mod2' Trial (no response)
'DMS' Trial (response)
'DMS' Trial (no response)
fix.
mod. 1
mod. 2
Target
time

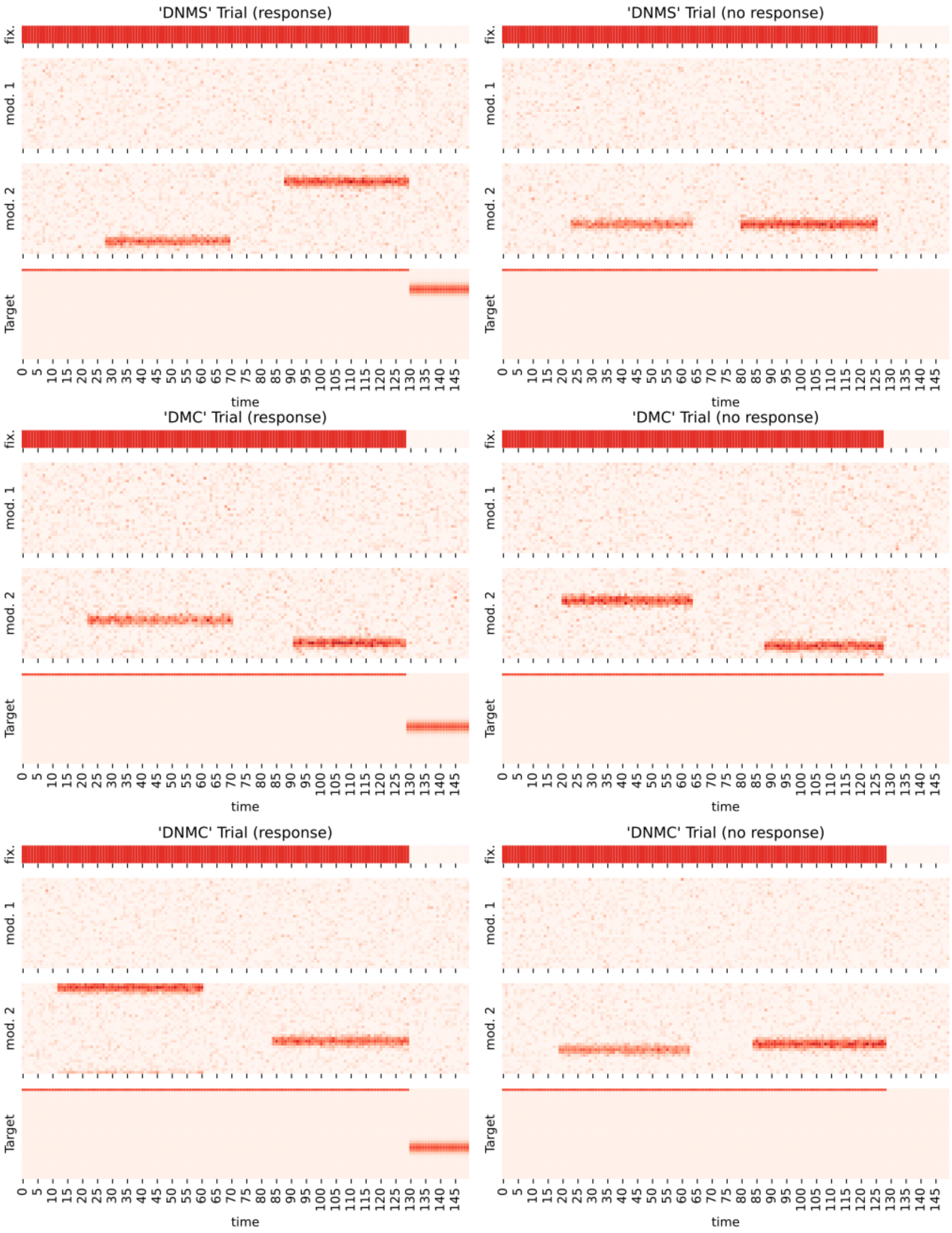
'DNMS' Trial (response)
'DNMS' Trial (no response)
'DMC' Trial (response)
'DMC' Trial (no response)
'DNMC' Trial (response)
'DNMC' Trial (no response)
fix.
mod. 1
mod. 2
Target
time
0 5 10 15 20 25 30 35 40 45 50 55 60 65 70 75 80 85 90 95 100 105 110 115 120 125 130 135 140 145
